# NMR structures and magnetic force spectroscopy studies of small molecules binding to models of an RNA CAG repeat expansion

**DOI:** 10.1101/2024.08.20.608150

**Authors:** Amirhossein Taghavi, Jonathan L. Chen, Zhen Wang, Krishshanthi Sinnadurai, David Salthouse, Matthew Ozon, Adeline Feri, Matthew A. Fountain, Shruti Choudhary, Jessica L. Childs-Disney, Matthew D. Disney

## Abstract

RNA repeat expansions fold into stable structures and cause microsatellite diseases such as Huntington’s disease (HD), myotonic dystrophy type 1 (DM1), and spinocerebellar ataxias (SCAs). The trinucleotide expansion of r(CAG), or r(CAG)^exp^, causes both HD and SCA3, and the RNA’s toxicity has been traced to its translation into polyglutamine (polyQ; HD) as well as aberrant pre-mRNA alternative splicing (SCA3 and HD). Previously, a small molecule, **1**, was discovered that binds to r(CAG)^exp^ and rescues aberrant pre-mRNA splicing in patient-derived fibroblasts by freeing proteins bound to the repeats. Here, we report the structures of single r(CAG) repeat motif (5’C<u>A</u>G/3’G<u>A</u>C where the underlined adenosines form a 1×1 nucleotide internal loop) in complex with **1** and two other small molecules via nuclear magnetic resonance (NMR) spectroscopy combined with simulated annealing. Compound **2** was designed based on the structure of **1** bound to the RNA while **3** was selected as a diverse chemical scaffold. The three complexes, although adopting different 3D binding pockets upon ligand binding, are stabilized by a combination of stacking interactions with the internal loop’s closing GC base pairs, hydrogen bonds, and van der Waals interactions. Molecular dynamics (MD) simulations performed with NMR-derived restraints show that the RNA is stretched and bent upon ligand binding with significant changes in propeller-twist and opening. Compound **3** has a distinct mode of binding by insertion into the helix, displacing one of the loop nucleotides into the major groove and affording a rod-like shape binding pocket. In contrast, **1** and **2** are groove binders. A series of single molecule magnetic force spectroscopy studies provide a mechanistic explanation for how bioactive compounds might rescue disease-associated cellular phenotypes.

**Graphical Abstract:** 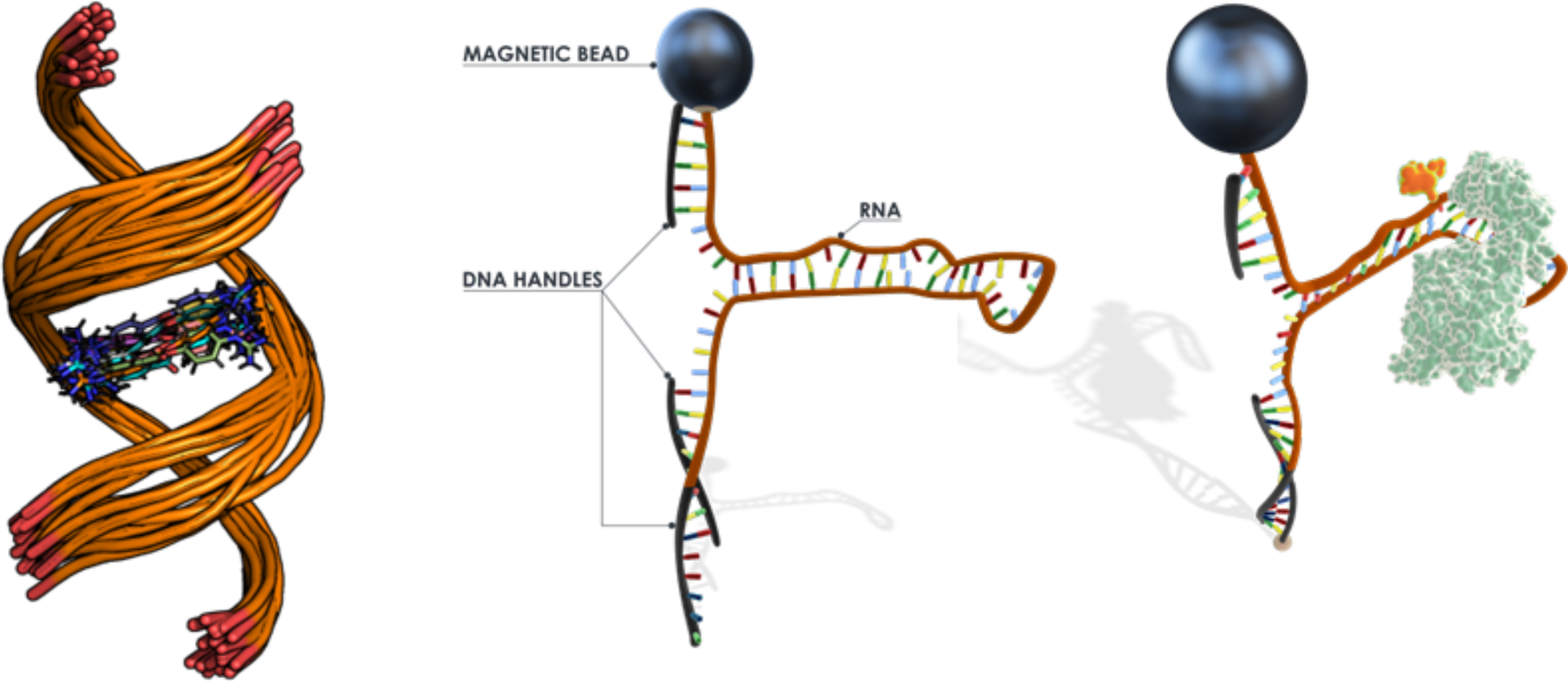

## INTRODUCTION

RNA functions in a variety of cellular processes, including pre-mRNA alternative splicing, transcription, translation, degradation, and RNA transport,(1) where these functions are predicated by the RNA’s structure. Misfolding or destabilization of RNA structures, formation of structure due to mutations or repeat expansions, and aberrant RNA expression each contribute to disease, for examples frontotemporal dementia and parkinsonism linked to chromosome 17 (FTDP-17) and other tauopathies,(2,3) Lewy-body dementia and synucleinopathies,(4,5) heart disease,(6) and cancer,(7) among others.

Repeat expansion disorders are a group of more than 40 neurological and neuromuscular diseases that develop when long, tandem RNA repeats surpass a certain threshold and fold into stable structures.(8-10) Huntington’s Disease (HD) and several of the spinocerebellar ataxias (SCAs) are caused by r(CAG) repeat expansions [r(CAG)^exp^], and the rate of disease progression after onset depends on repeat length.(11,12) For both HD and SCA3, r(CAG)^exp^ causes toxicity due to a gain-of-function where the nature of the aberrant function is dependent upon its location in the corresponding gene, for example in an open reading frame or an untranslated region (UTR).(13-16) This gain-of-function is traced to the structure formed by the mutant allele, where the RNA folds onto itself to form hairpin structures with an array of 1×1 nucleotide AA internal loops, or 5’C<u>A</u>G/3’G<u>A</u>C. Indeed, this gain-of-function caused by an aberrant RNA structure is observed in other microsatellite disorders.(17)

One therapeutic strategy for microsatellite diseases is to target the structures that confer gain-of-function with small molecules, thereby deactivating the toxic repeat expansion. Such an approach has been previously taken, affording small molecules that bind with nM – μM affinities and rescue disease-associated cellular defects.(18-20) Herein, we report the solution structure of the *apo* form of the 5’C<u>A</u>G/3’G<u>A</u>C motif that is formed by r(CAG)^exp^ as well as the solution structures of the complex formed by the r(CAG) repeat and three small molecules, as determined by nuclear magnetic resonance (NMR) spectrometry and restrained molecular dynamics (MD). Further, single molecule techniques were applied to study the binding of the three molecules to 5’C<u>A</u>G/3’G<u>A</u>C, including their abilities to inhibit formation of a complex between r(CAG)^exp^ and the RNA-binding protein (RBP) muscleblind-like 1 (MBNL1), a regulator of pre-mRNA alternative splicing, the sequestration of which causes splicing defects observed in HD and SCA3.(21,22)

## METHODS

### Compounds

Compound **1** was synthesized as described in (18), and compound **2** was synthesized as described in (23). Compound **3** was obtained from Pfizer.(24)

### Preparation of NMR samples

The oligoribonucleotide r(GACAGC<u>A</u>GCUGUC) (“r(CAG) RNA”) was purchased from GE Dharmacon, Inc.. The deprotected and desalted RNA was dissolved in NMR Buffer [5 mM KH_2_PO_4_/K_2_HPO_4_, 0.25 mM EDTA, pH 6.0], and folded by heating to 70 °C for 3 min. The concentration of r(CAG) RNA was 0.7 mM in free form and r(CAG)-**1** samples, 0.3 mM in r(CAG)-**2** samples, and 0.4 mM in r(CAG)-**3** samples. Compound **1** was added to r(CAG) samples to a final concentration of 0.7 mM, and **2** and **3** were added to r(CAG) samples to a final concentration of 0.6 mM. For single molecule studies r(CAG)_21_ RNA was purchased from IDT. MBNL1 protein was purchased from Abnova (H000041540P02, lot N2151).

### NMR spectroscopy

NMR spectra of samples in Shigemi tubes (Shigemi, Inc.) were acquired on Bruker Avance III 700 and 850 MHz spectrometers. WaterLOGSY (water-ligand observed via gradient spectroscopy) spectra were acquired on samples containing 300 µM of compound with or without 3 or 6 µM of RNA. Signals were phased to give negative NOEs for RNA binders. One-dimensional spectra were acquired at 25 °C with excitation sculpting to suppress the water signal.(25) Two dimensional NOESY and COSY spectra were acquired on free form and bound form RNAs at 5 °C or 9 °C and 25 °C. For the r(CAG)-**2** and r(CAG)-**3** complexes, spectra were also acquired at 35 °C. Proton chemical shifts were referenced to water. 2D NMR spectra were processed with NMRPipe (26) and assigned with SPARKY (**Tables S1-S4**).(27)

### Methods for obtaining distance and dihedral restraints

Distance restraints for pairs of protons were calculated by integrating NOE volumes in SPARKY(27) or manually assigned a range of distances based on relative NOE intensities (**Tables S3-S5**). NOE volumes were referenced to those calculated from fixed distances: H2′-H1′ (2.75 Å); cytosine or uracil H5-H6 (2.45 Å).(28) Hydrogen bonds in canonical AU, GC, and GU pairs were assigned to distances of 2.1 ± 0.3 Å. NOE intensities between H1′ and H6/H8 indicate that all residues adopted an *anti*-conformation. Therefore, the *χ* dihedral angle was constrained between 170° and 340° (*anti*) for all residues except terminal residues and loop adenosines or uridines.

### Modeling methods

Structures were calculated with a simulated annealing protocol, using a starting structure from Nucleic Acid Builder.(29) Restrained molecular dynamics simulations were carried out with AMBER (30) using the parm99χ_YIL force field.(31) Solvation was simulated with the general Born implicit model and 0.1 M NaCl.(32) The system was heated from 0 to 1,000 K in 5 ps, cooled to 100 K in 13 ps, and then to 0 K in in 2 ps. Force constants were 20 kcal mol^−1^ Å^−2^ for NOE restraints and 20 kcal mol^−1^ rad^−2^ for dihedral angle restraints. The simulated annealing procedure was repeated with different initial velocities to generate an ensemble of 100 structures. For each construct, the 20 structures with the fewest distance restraint violation energies were selected as the final ensemble of structures. For **1**- and **2**-bound r(CAG), ensembles of 60 structures were generated by simulated annealing, among which the 20 lowest energy structures without distance violations above 0.1 Å were selected as the final ensemble without further refinement. Root mean square deviations (RMSDs) of the ensemble of structures were calculated with VMD.(33)

### System preparation and molecular dynamics (MD) simulations

AMBER 18 (30) was used for MD simulations using the PARM99 (34) force field with revised *χ (31)* and *α*/*γ* (35) torsional parameters.. Each system was first neutralized with Na^+^ ions (36) and then solvated with TIP3P (37) water molecules in a truncated octahedral box with periodic boundary conditions extended to 10 Å using the LEAP (38) module of AMBER 18. The structures were minimized with the Sander module each in two steps. Positional restraints on RNA heavy atoms with restraint weights of 10 kcal mol^-1^ Å^-2^ were applied in the first step of minimization with 5000 steps of steepest-descent algorithm followed by 5000 steps of conjugate-gradient algorithm using the CPU implementation of Sander force field to avoid the truncation of forces or overflow of the fixed precision representation. The second round of minimization was performed without restraints with 10,000 steps of steepest descent. Minimization was followed by an equilibration protocol first in constant volume dynamics (NVT), where positional restraints were imposed on the RNA heavy atoms with restraint weights of 10 kcal mol^-1^ Å^-2^ while temperature was gradually increased from 0 K to 300 K within several nanoseconds using the Langevin (39) thermostat. A second round of equilibration was performed at constant pressure (NPT), where temperature and pressure coupling (40) were set to 300 K and 1.0 ps^-1^, respectively, while constraints were gradually removed. After minimization and equilibration, MD simulation with a 2 ps time step was performed using NPT dynamics with isotropic positional scaling. The reference pressure was set to 1 atm with a pressure relaxation time of 2 ps. SHAKE (41) was turned on for constraining bonds involving hydrogen atoms. An atom-based long-range cut-off of 10.0 Å was used in the production runs. The reference temperature was set to 300 K. The Particle Mesh Ewald (42) (PME) was used to handle the electrostatics and the Langevin (43) thermostat was applied with a coupling constant *γ* = 1.0 ps^-1^. Simulations were performed using the pmemd.cuda implementation (GPU accelerated) of AMBER18 with NMR restraints.

### Analyses

Trajectory analysis was completed with the CPPTRAJ module of AmberTools18.(44) Cluster analysis was done using the average-linkage hierarchical agglomerative method. Heavy atoms of 1×1 A/A loop residue were used in the cluster analyses with a root-mean-square deviation (RMSD) of 1.0 Å. Base pair step parameters, groove widths as well as bending angles and curvilinear helical axis were measured using Curves+ (45) and 3DNA.(46)

### Calculation of the potential of mean force (PMF)

We calculated the PC1 (principal component 1) and PC2 (principal component 2) of the loop and the immediate neighboring base-pairs, yielding four base-pairs in total using the CPPTRAJ module and used them as reaction coordinates to generate the free energy.

### Single molecule preparation for magnetic force spectroscopy

The RNA containing 21 CAG repeats [r(CAG)_21_] was annealed to two DNA splints in equimolar ratio in Annealing Buffer (10 mM Tris-HCl, pH7.4, 50 mM NaCl, and 1 mM EDTA). Once annealed, the structures were purified through MicroSpin S-200 HR spin columns (Cytiva) and were mixed with an equal volume of GenTegraRNA^TM^ (GenTegra) before storing at -20°C until further use.

The annealed RNA structure (4 fmol) was mixed with 3 µl of MyOne T1 streptavidin beads (Invitrogen) in Hybridization Buffer (10% PEG8000, 5× SSC Buffer) for 10 min at room temperature. The beads were washed twice in OB Buffer (1× PBS, 0.2% (w/v) bovine serum albumin (BSA), 0.1% (w/v) sodium azide) to remove unbound RNA structures, and the resulting beads containing the RNA were resuspended in OB Buffer before loading in the instrument.

### Magnetic force spectroscopy

All the experiments presented in this paper were acquired on a Stereo Darkfield Interferometry system (SDI) prototype instrument.(47) To attach the RNA molecules at the surface of the flow cell, a functionalized azide-coated coverslip (Susos) was grafted with a surface oligo (100 nM) harboring a 3′-DBCO group through click chemistry in the Click Buffer (500mM NaCl, 1 µM PEG-DBCO) for 2 h. The flow cell was then passivated with OB Buffer after assembly. The RNA bound to the MyOne beads was injected into the flow cell and left to hybridize for 30 min in OB Buffer. After the molecules were captured at the surface of the flow cell, the buffer was exchanged to the Test Buffer (20 mM HEPES, pH7.4, 100 mM KCl, 2 mM MgCl_2_, 5 mM DTT, and 1% (v/v) DMSO). The unbound structures were washed away, and the non-specific bound beads were removed by increasing the magnetic force above 25 picoNewton (pN). The experiments were performed at 22°C, and the data were recorded at a frequency of 30 Hz.

A control experiment was always recorded first in Test Buffer, which was used to normalize as well as to define the number of analyzable molecules. Then, experiments were performed with increasing concentration of either compound **1**, **2**, and **3**, or MBNL1 protein (in the absence or presence of 10 or 100 µM **1**, **2**, or **3**) in the Test Buffer, maintaining the concentration DMSO at 1% (v/v). In experiments where a compound of interest and MBNL1 protein were tested together, the compound and the protein were mixed in Test Buffer just prior to injecting into the flow cell.

Two types of experiment were performed on the instrument to extract the information: 1) Force ramp experiments, consisted of moving the magnet position slowly (0.125 mm/s) to increase the force applied to the beads from ∼0.2 pN to ∼28 pN, then maintaining the highest force for 1 s before moving back down to low force, with repetition of these steps for 100 cycles; and 2) For stepped force experiments, the magnet was maintained at a constant position corresponding to a constant force for 30 s, starting at ∼6 pN, then moved closer to the flow cell surface in a stepwise manner until the force reached ∼19 pN.

### Analysis of the force ramp experiments

The first step in the analysis is to select the beads with a functional RNA secondary structure. The data were first cleaned by removing signals with high noise (>1000nm). In order to be considered as an analyzable RNA molecule and used in downstream analyses, each molecules had to satisfy the following parameters: 1) unfolding force between 5-15 pN; 2) unfolding size between 5-30 nm in the DMSO control condition; 3) the total number of cleaned cycles must be above 20 cycles; 4) control condition has more than 80% of cleaned cycles; 5) the structure is present in more than half of the conditions tested; and 6) the width of the unfolding force distribution in control conditions is less than 2 pN. Following this selection, each cycle of each molecule was analyzed to determine the force required to unfold and fold the RNA structures as well as the size of the detected jump using python module HDBScan.(48) The structure was considered closed if all cycles did not contain unfolding or refolding events. For each analyzable structure, the unfolding and refolding probability were calculated by dividing the number of cycles with an abrupt change in bead vertical position (jump event) over the total number of cycles for a single molecule. The median unfolding and refolding probabilities of all structures were plotted against ligand or protein concentrations.

The unfolding force of the detected jumps in the DMSO control condition were used for normalization. The median unfolding force of each analyzable structure was used to normalize all cycles of all conditions for the given structure. The normalized unfolding and refolding forces were plotted to study the changes in force distribution upon ligand or protein binding. The median normalized unfolding force for each structure was used, and the median force for all structures was used against ligand or protein concentrations. The concentration with 50% maximum effect (EC_50_) was calculated and a sigmoid curve was fitted:

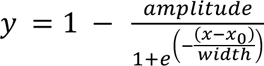

where *x*_0_ is the *f*1⁄2 (EC_50_) value.

To calculate the low-force fraction, a threshold was set for the 5% quantile of the force distribution in DMSO control conditions. Next, the number of cycles that have an unfolding or refolding force that is below the threshold was summed over the total number of cycles for that molecule. The median of all structures was plotted against ligand or protein concentrations. To calculate the EC_50_, the data were fitted with Hill’s equation with an offset:

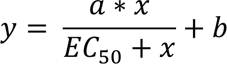

where a is the scaling factor and b is the offset.

### Analysis of stepped force experiments

Before analysis, the data were cleaned by removing any consecutive data points varying by greater than 40 nm (within a 20-frame rolling window) or discarding any force-step with less than 250 data points. To quantify the amplitude of the folding and unfolding of the structure, the variance (squared distance to mean value) per force step was measured using the following method: 1) the data were partitioned into 40-frame regions, over which the mean of the variance computed over a 20-frame rolling window was calculated; and 2) the estimated variance for that force step was the median variance of these partitions. For the control condition, the baseline was estimated based on a two-state model, and this baseline was removed for the same structure in all other conditions. After discarding RNA molecules with a low signal-to-noise ratio and baseline removal, the force of each structure was normalized with the force with maximum variance in the control condition. The variance was normalized with the maximum variance in the control condition. For all RNA molecules, the normalized variances were binned according to normalized forces with a bin size of 0.03. The median value for each bin was displayed and overlayed with a curve using third order splines. For each structure, the normalized force at maximum variance, as well as the maximum normalized variance height were calculated, and the median of all structures were plotted against MBNL1 protein concentration.

## RESULTS & DISCUSSION

### Design of a model of r(CAG)^exp^ suitable for NMR studies and selection of small molecules

The model RNA to study the structure of r(CAG) repeats was carefully designed to eliminate spectral overlap in NMR spectra. In particular, a self-complementary duplex was designed where the 5’C<u>A</u>G/3’G<u>A</u>C motif formed by r(CAG)^exp^ was flanked by five additional base pairs both 5’ and 3’, or r(GACAGC<u>A</u>GCUGUC)_2._ This design exemplifies the CAG repeat motif incorporating the A/A internal loop as a primary target for small molecules.

Compound **1** was previously reported to bind to the 5′C<u>A</u>G/3′G<u>A</u>C motif and inhibits formation of r(CAG)^exp^-MBNL1 complex both in vitro and in cells.(18) Further, optical melting studies showed that **1** increases the thermal stability of the RNA.(18) With the goal of improving the affinity and selectivity of **1**, a derivative of the compound, **2**, was synthesized.(23) In particular, the guanidines of **1** were replaced with imidazolines to reduce polarity while the ester linkage was replaced with an amide to increase metabolic stability and rigidity. Delocalization of electrons across the amide bond creates a partial double bond character which leads to increased rigidity.(49) Compound **3** binds an A bulge present in microtubule associated protein tau (*MAPT*) pre-mRNA that is part of a splicing regulatory element (SRE).(24) The solution structures of these three small molecules in complex with a model of the r(CUG) repeats that cause various microsatellite diseases were recently reported.(23)

### NMR analysis of *apo* form of r(CAG)

Analysis of the NMR spectra of the free (*apo*) form of the r(CAG) duplex (0.7 mM), collected in NMR Buffer (5 mM KH_2_PO_4_/K_2_HPO_4_, pH 6.0 and 0.25 mM EDTA), suggested that the RNA adopts an A-form geometry. Particularly, a sequential H6/H8-H1′ walk, in addition to interresidue H6/H8-H2′ nuclear Overhauser effects (NOEs), in 2D NOESY (Nuclear Overhauser Effect Spectroscopy) spectra were observed (**Figure S1; Tables S1 and S2**). Intrastrand and interstrand NOEs between adenine H2 protons and H1′ protons of nearby residues provided information about the conformation of the helix, especially the 5′C<u>A</u>G/3′G<u>A</u>C motif. Specifically, NOEs were observed from A2H2 to C3H1′ and C13H1′, from A4H2 to G5H1′ and G11H1′, and from A7H2 to G8H1′. Cross-peaks between adenine H2 and uracil imino protons were assigned for all AU pairs while cross-peaks between guanine imino and cytosine amino protons were assigned for all GC pairs except for the terminal GC pair (**Figure S2 and Table S1)**.

### Structure of *apo* form of a r(CAG) repeat using restrained MD

In agreement with the observations from 2D NOESY spectra, the 20 lowest energy structures from NMR-restrained MD simulations (via simulated annealing) each adopted an A-form conformation (**Figures S3 and Tables S6 and S7**). The root mean square deviation (RMSD) of all heavy atoms for the ensemble was 0.77 ± 0.26 Å, indicating good convergence (**Table 1**). The average helical rise and twist for the 20 structures were slightly less than average for, but otherwise consistent, with A-form RNA.(50,51) The average χ dihedral angles ranged from −149° to −169° for all residues, corresponding to *anti* conformations.(51)

**Table 1:**
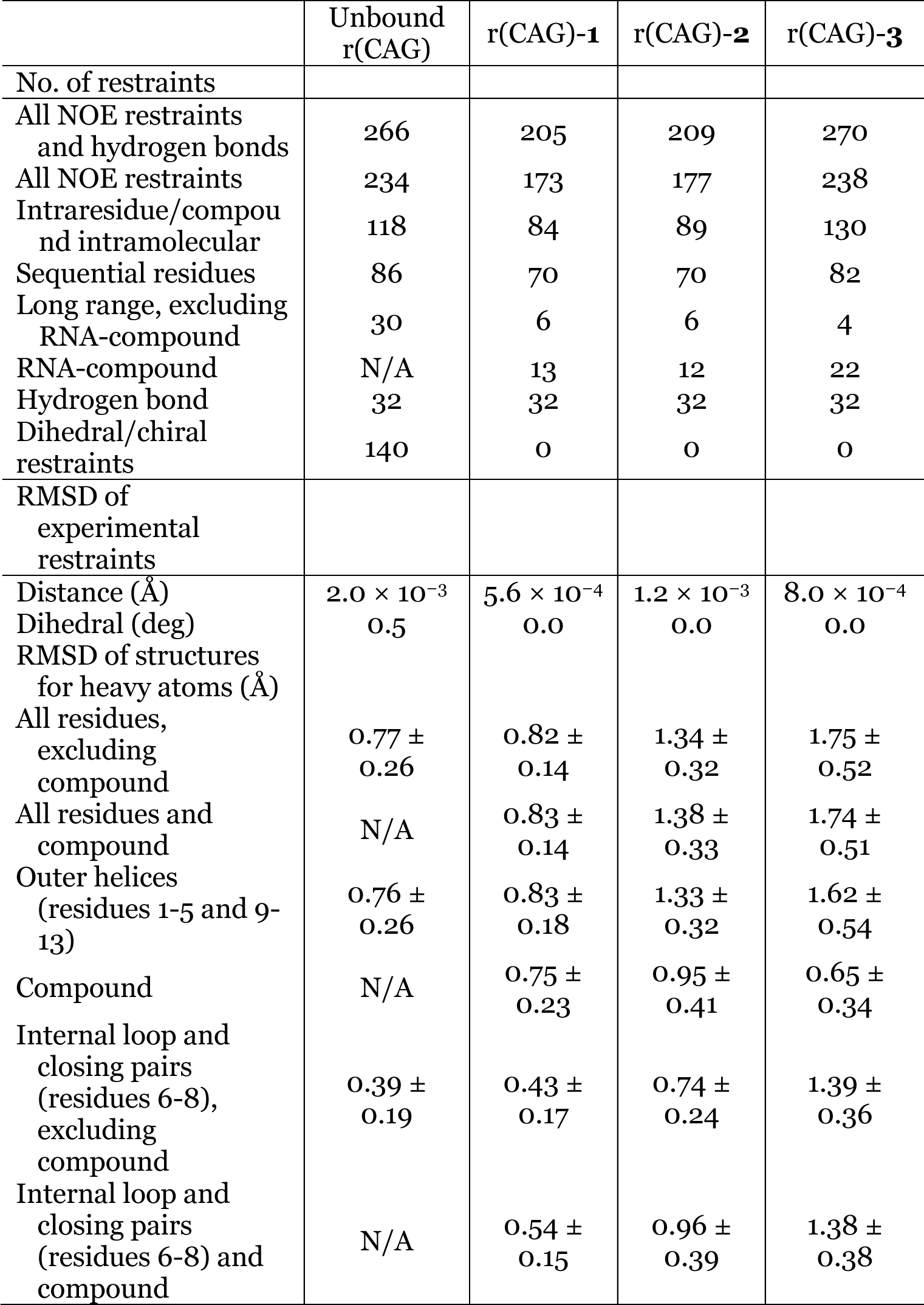
Structural refinement statistics for the average of 20 structures for unbound and ligand bound RNAs.

Within the 5′C<u>A</u>G/3′G<u>A</u>C motif itself, the helix was undertwisted at the 5′CA/3′GA and 5′AG/3′AC steps and overtwisted at the flanking 5′GC/3′CG steps. The AA mismatch adopted a *cis*-Watson Crick/Watson-Crick base pair stabilized by a single N6-H6···N1 hydrogen bond and stacking interactions with the neighboring (closing), canonical GC pairs (50). The average C1′-C1′ distance for the AA pair was 12.3 ± 0.1 Å, greater than ∼10.5 Å for typical A-form RNA helices, to accommodate the observed hydrogen bond. The AA mismatch had buckle, opening, shear, and stagger values close to those of the Watson-Crick pairs in the helix, but the stretch value differed more as compared to the other base pairs in the helix. Together, these values are consistent with a *cis* Watson Crick/Watson-Crick AA pair.(52,53) The nonplanarity of the AA mismatch was indicated by a large propeller twist. Altogether, the 1×1 nucleotide A/A internal loop induces distortions in the RNA helix to stabilize its formation.

Other *apo* structures of the 5’C<u>A</u>G/3’G<u>A</u>C motif have been previously reported, including NMR solution structures of six and three tandem motifs (repeats; represented as r(6×CAG) and r(3×CAG), respectively) (54,55) and X-ray crystal structures of two, five, and three tandem repeats.(56-58) The one hydrogen bond state of the AA pair, with elongated C1′-C1′ distances, observed herein is consistent with that observed in AA mismatches in the r(3×CAG) construct (55). NMR spectral studies of a hairpin with six r(CAG) motifs suggested that the AA mismatches adopt an *anti-anti* conformation undergo exchange between a zero-hydrogen bond and a one hydrogen bond state (N6-H6···N1).(54)

The X-ray crystal structures of r(CAG)-containing RNAs share some structural features with the NMR solution structures. The average helical twist angles of the Kiliszek *et al*. and Yildirim *et al*. structures are close to those observed for the *apo*-r(CAG) and r(3×CAG) NMR structures. All three reported X-ray structures contained mismatched adenines in the *anti-anti* conformation with elongated C1′-C1′ distances to prevent steric clashing.(56-58) However, these adenines interacted via a C2-H2···N1 hydrogen bond. Yildirim *et al*. also observed an N6-H6···N1 hydrogen bond in the *syn-anti* conformation,(58) observed herein, but without an elongated C1′-C1′ distance. As carbon does not form strong hydrogen bonds, the AA mismatches are more likely stabilized by a N6-H6···N1 hydrogen bond.

A closer inspection of the helical geometries around the AA loops revealed additional similarities among the X-ray crystal and NMR structures. In these structures, the values of the α- and γ-torsions in the 3′ adjacent guanosines of *anti-anti* AA mismatches deviated from the averages for A-form RNA (295°/−65° and 54°, respectively).(59) These values correspond to undertwisting of the 5′AG/3′AC steps. To compensate, the adjacent 5′GC/3′CG steps are overtwisted. A majority of the *anti-anti* AA mismatches in these structures contain at least one adenine with a λ-angle outside of the range for Watson-Crick pairs (typically 51°-60°),(60) indicating that these adenosines are shifted towards the major groove.(56) In summary, the X-ray and NMR structures of r(CAG) show that the AA mismatches distort the helix around the r(CAG) motif (**Supplemental Data Set S1**).

A comparison of the *apo*-r(CAG) and r(3×CAG) structures with crystal structures of fully base paired duplexes analyzed using X3DNA (52) showed no significant differences in the average values of helical parameters, including helical rise, and helical twist, and minor groove widths (**Table S1**). The average helical rise and twist of fully base paired structures (61-66) are within range of those observed for *apo*-r(CAG) and r(3×CAG) (**Tables S1**). Differences, however, are observed in the major groove of the 5’C<u>A</u>G/3’G<u>A</u>C internal loop, which is widened compared to fully paired structures. Additionally, the r(CAG)-containing structures have larger C1′-C1′ distances than the fully base paired constructs and disparities in α- and γ-torsions and λ-angles. Collectively, it appears that r(CAG)-containing structures are characterized by backbone distortions and larger C1′-C1′ distances and major groove widths within the r(CAG) motifs than fully base paired RNA constructs. These structural features may allow small molecules to recognize r(CAG) repeats selectively.

### NMR analysis of RNA-small molecule complexes: 1D imino ^1^H and WaterLOGSY spectra

To gain preliminary insight into the binding of **1** – **3** to the r(CAG) repeat duplex model, WaterLOGSY and imino ^1^H spectra were acquired. In the absence of the RNA, WaterLOGSY spectra of **1**, **2**, and **3** alone contained positive NOEs, indicating that the compounds did not aggregate in NMR conditions (**Figures S4-S6**). Addition of the RNA duplex to each compound afforded negative NOEs, resulting from binding of the compounds to the RNA. In summary, **1**, **2**, and **3** formed soluble complexes with the r(CAG) duplex.

Small molecule binding was also observed in RNA-observed, imino ^1^H spectra, where in each case binding can be traced to the 5’C<u>A</u>G/3’G<u>A</u>C motif (**Figures S7 – S9**). Addition of **1** to the r(CAG) repeat duplex model caused a downfield shift of G8H1 (starting at a 1:1 small molecule: RNA ratio; equivalent to G21), which along with C19 forms the loop closing base pairs in the self-complementary duplex (**Figure S7**). Interestingly, at **1**:RNA ratios of 2:1, 4:1, and 6:1, the overlapped G5H1 and U10H3 resonances shifted away from each other, indicating that the binding of **1** transmits structural changes near the internal loop. Thus, **1** appears to bind to the r(CAG) motif.

As observed upon the addition of **1**, changes in the imino ^1^H spectra of the r(CAG) duplex were observed upon addition of **2**, particularly the guanosines that form the loop’s closing base pairs (G8H1/G21 H1) (**Figure S8**). Throughout the titration, the resonance for G8H1 shifted downfield and eventually overlapped with G11H1. As observed in the titration of the r(CAG) duplex with **1**, the overlapped G5H1 and U10H3 resonances shifted away from each other upon compound binding. Shifting of U12H3 upfield at higher **2**:RNA ratios suggests that the small molecule stacked non-specifically on the end of the helix. The shifting of the resonances throughout the titration upon binding of **2** indicates that the small molecule forms interactions with the r(CAG) motif.

Addition of **3** to the r(CAG) duplex led to upfield shifts of G5H1 and G8H1 (closing base pairs) and downfield shifts of U10H3 and U12H3 (**Figure S9**). Compared to the peaks in the absence of small molecules, these peaks also broadened considerably, indicating that the r(CAG) motif may be destabilized and G8H1 may be more dynamic and possibly exchanging with solvent. At 3:1 **3**:RNA ratio, G11H1 also shifted downfield slightly. These results showed that **3** binds to r(CAG) and disrupts the internal loop motif (5’C<u>A</u>G/3’G<u>A</u>C) and the AA mismatch. As observed for **2** at higher small molecule:RNA ratios, shifting of the imino peaks near the end of the helix suggests that the compound stacks at the end of the helix non-specifically at higher concentration of small molecule.

In summary, the 1D imino ^1^H and WaterLOGSY spectra show that the small molecules bind to the r(CAG) duplex. Thus, 2D NMR spectra were acquired on these RNA-small molecule complexes to elucidate their structures.

### 2D NMR spectral analysis of a r(CAG) repeat bound to 1 and 2

The 2D spectra of the imino region of r(CAG) bound to **1** and **2** indicate formation of expected Watson-Crick base pairs in the RNA construct (**Figures S10 and S11**). Aromatic proton resonances of **1** appeared at 7.16, 7.20, 7.27, and 7.89 ppm (**Figure 1**) and those for **2** appeared at 7.51, 7.59, 7.96, and 7.71 ppm (**Figure 2**) in 2D NMR spectra of r(CAG) in complex with these compounds. In the 125 ms 2D NOESY exchangeable proton spectrum of both complexes acquired in 95% H_2_O, G8H1 formed intrastrand NOEs with C9H1′ and interstrand NOEs with C6 amino protons and A7H1′ (**Figures S10 and S11**). This clearly indicates that GC pairs flanking the AA mismatch were hydrogen bonded.

**Figure 1:**
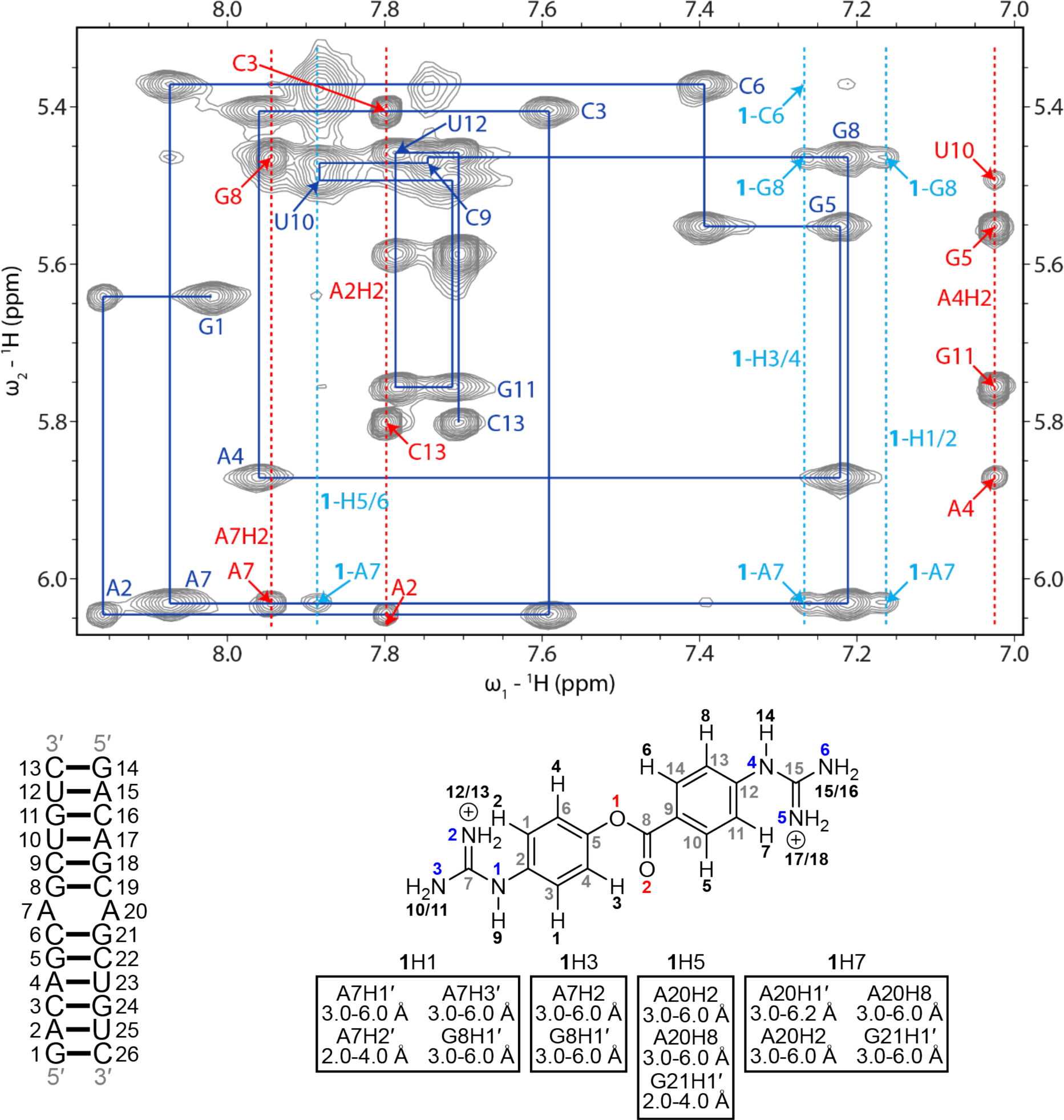
H6/H8-H1′ region of a 2D ^1^H-^1^H NOESY spectrum of the r(CAG)-1 complex. A sequential H6/H8-H1′ walk is shown with blue lines. Blue labels correspond to intraresidue H6/H8-H1′ NOEs. Adenine H2 resonances are labeled red with dashed lines and NOEs between r(CAG) and **1** are labeled light blue with dashed lines. RNA residues are only numbered from 1 to 13 in the NMR spectrum, and residues 14 to 26 are sequentially the same as 1 to 13. In the chemical structure of **1**, black numbers correspond to hydrogens, gray numbers correspond to carbons, blue numbers correspond to nitrogen, and red numbers correspond to oxygen. Intermolecular NOEs between r(CAG) and **1** and their corresponding distance restraints used for modeling are colored according to the atoms of **1**. The spectrum was acquired at 25 °C with 400 ms mixing time and 0.7 mM of RNA and **1**.

**Figure 2:**
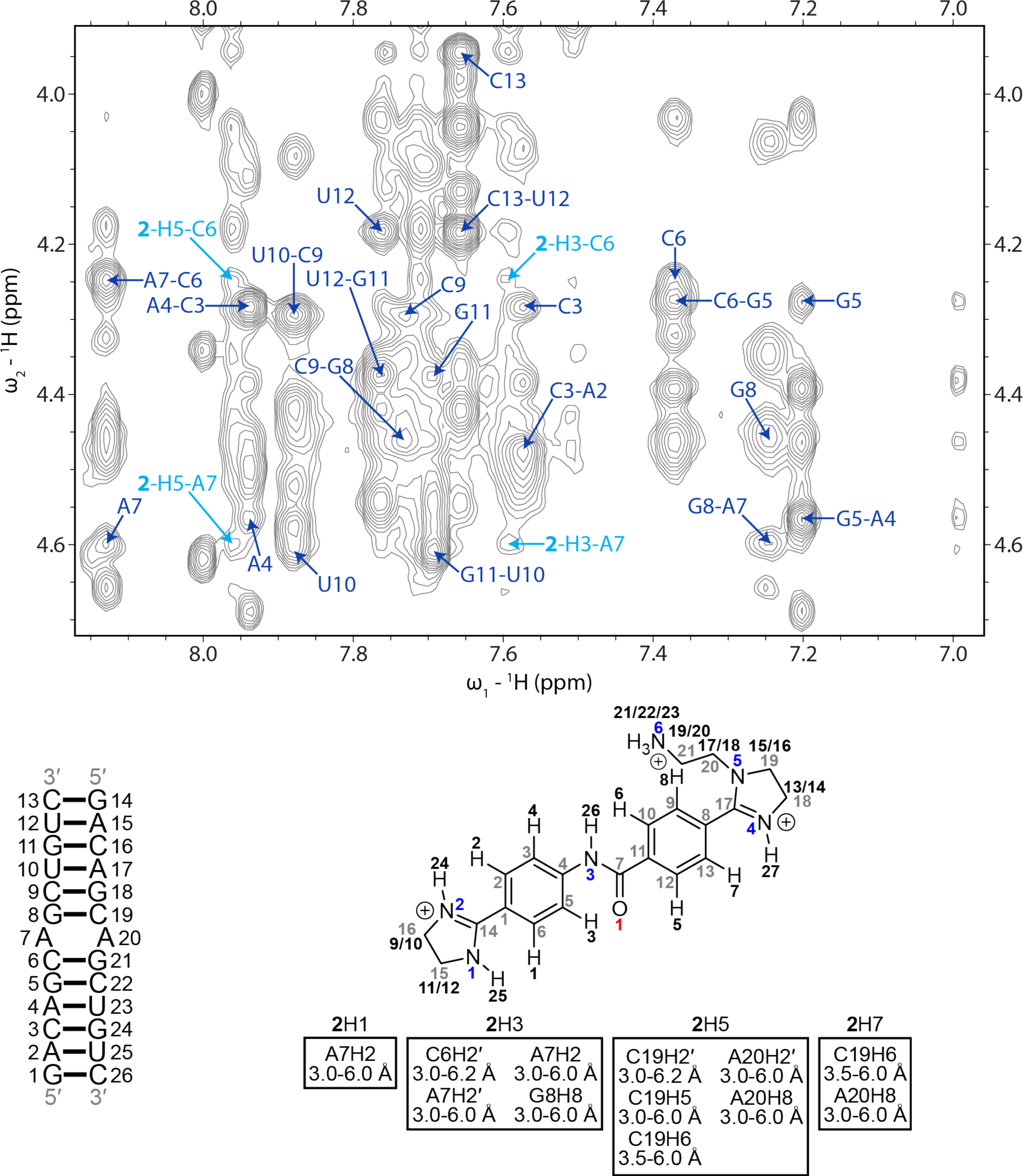
H6/H8-H2′/H3′/H4′ region of a 2D ^1^H-^1^H NOESY spectrum of the r(CAG)-2 complex. Blue labels correspond to intraresidue and interresidue H6/H8-H2′ NOEs. NOEs between r(CAG) and **2** are labeled light blue. RNA residues are only numbered from 1 to 13 in the NMR spectrum, and residues 14 to 26 are sequentially the same as 1 to 13. In the chemical structure of **2**, black numbers correspond to hydrogens, gray numbers correspond to carbons, blue numbers correspond to nitrogen, and red numbers correspond to oxygen. Intermolecular NOEs between r(CAG) and **2** and their corresponding distance restraints used for modeling are colored according to the atoms of **2**. The spectrum was acquired at 25 °C with 400 ms mixing time and 0.3 mM of RNA and 0.6 mM of **2**.

Intermolecular NOEs were observed in a 400 ms NOESY spectrum of **1** bound to the r(CAG) duplex. In particular, NOEs were observed from A7H1′ (loop nucleotide) and G8H1′ (loop closing base pair) to **1**-H1/H2 and **1**-H3/H4 (**Figure 1**). Within the H6/H8-H2′ region, NOEs were observed between **1**-H1/H2 and **1**-H3/H4 and A7H2′, A7H3′, and G8H2′. Additional NOEs were observed from **1**-H5/H6 to A7H1′ and A7H2′ and from **1**-H7/H8 to A6H2. Intermolecular NOEs between **1** and the terminal nucleotides suggests that end stacking of **1** occurs at higher concentrations of the compound.

Likewise, NOEs were observed between **2** and the nucleotides that comprise the 5’C<u>A</u>G/3’G<u>A</u>C internal loop. NOEs were observed from **2**-H3/H4 and **2**-H5/H6 to C6H2′ and A7H2′ (**Figure 2**). Additional NOEs were observed from **2**-H1/H2 and **2**-H3/H4 to A7H2, **2**-H3/H4 to G8H8, **2**-H5/H6 to C6H5 (loop closing base pair), C6H6, and A7H8, and **2**-H7/H8 to C6H6 and A7H8. In the r(CAG)-**2** spectrum, G8H1 also formed weak intermolecular NOEs with **2**. As in the spectra of r(CAG)-**1**, intermolecular NOEs between **2** and the terminal nucleotides indicate that higher concentrations of the compound results in end stacking.

Altogether, these data suggest that the r(CAG)-**1** and r(CAG)-**2** complexes conformationally comprise stable r(CAG) motifs with **1** and **2** within or near the AA mismatches.

### Structure of a r(CAG) repeat bound to 1

A total of 13 intermolecular NOE distance measurements were used as restraints to model the **1**-bound CAG complex (**Tables 1 and S8**). In the simulated annealing calculations, all 20 structures in the final ensemble of structures with the fewest distance restraint violations retained an A-form conformation in agreement with the NMR data (**Figure 1**). For the **1**-bound complex, the RMSD of all heavy atoms for the ensemble of 20 structures was 0.82 ± 0.14 Å (**Table 1**). Compound **1** bound within the major groove of the RNA and laid across the base pair plane, forming stacking interactions with the closing base pairs of the AA mismatch (**Figures 5A**). The mismatched adenines were partially displaced into the minor groove and formed two N6-H6···N1 hydrogen bond *trans* Watson-Crick/Watson-Crick base pair.(53) In addition, the amino groups of A7 and A20 formed hydrogen bonds with the G21 and G8 bases, respectively. The carbonyl oxygen was oriented towards the minor groove in all structures and formed dipole-dipole interactions with the 1×1 nucleotide A/A internal loop bases. In 15 of the structures, the guanidine(s) formed hydrogen bonds with the phosphate backbone of the opposite strand of the RNA, particularly at the A7 and/or A20 residues and, in some states, the C6 or G8 residues. Additionally, the guanidines formed electrostatic interactions (5 Å distance or less (67-69)) with the phosphate backbone at C6/C19, A7/A20, and G8/G21 in all of the structures. Van der Waals interactions between the **1** aromatic protons and A7 and A20 bases further stabilized the ligand-RNA complex.

**Figure 3:**
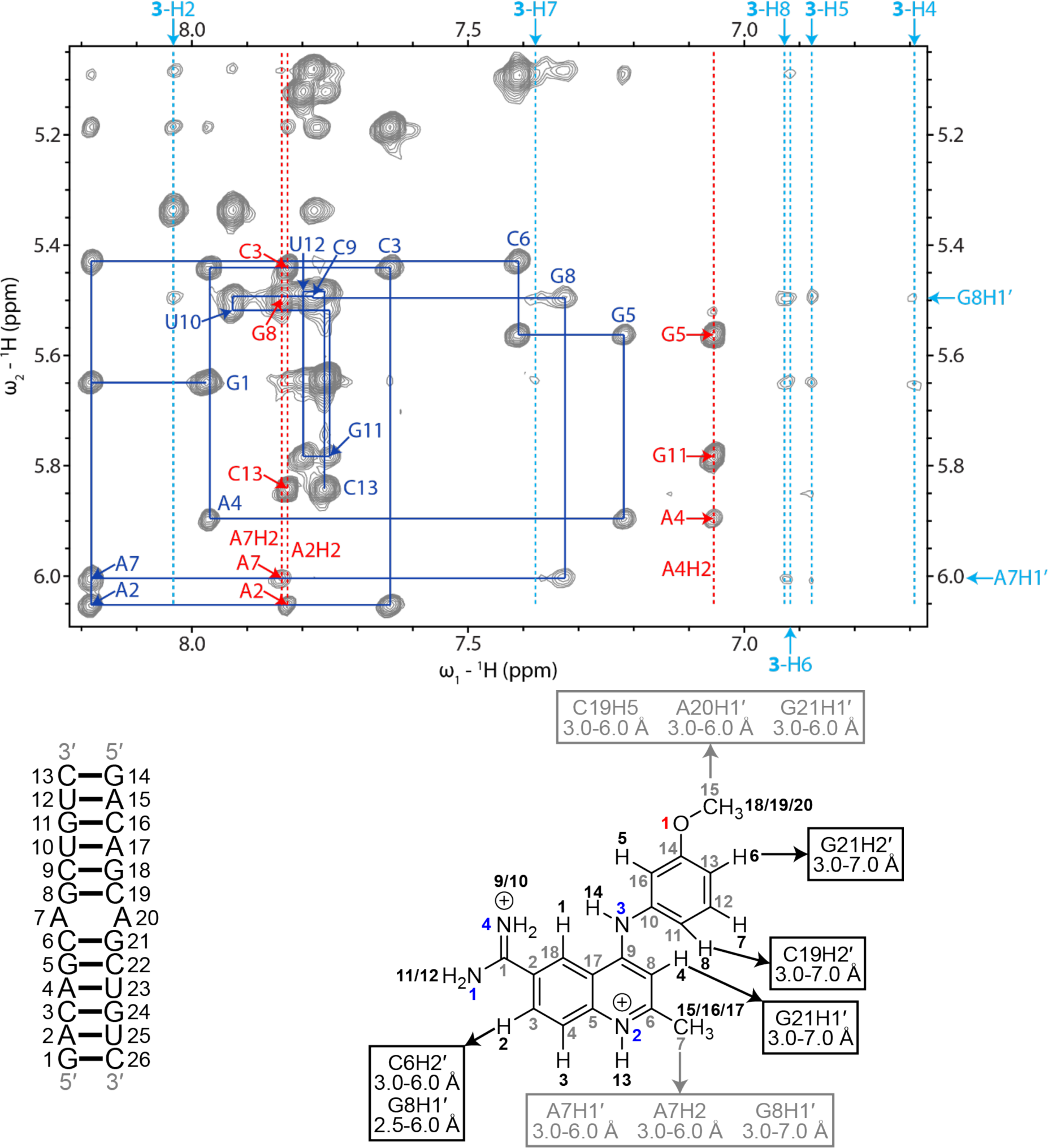
H6/H8-H1′ region of a 2D ^1^H-^1^H NOESY spectrum of the r(CAG)-3 complex. A sequential H6/H8-H1′ walk is shown with blue lines. Blue labels correspond to intraresidue H6/H8-H1′ NOEs. Adenine H2 resonances are labeled red with dashed lines and NOEs between r(CAG) and **3** are labeled light blue with dashed lines. RNA residues are only numbered from 1 to 13 in the NMR spectrum, and residues 14 to 26 are sequentially the same as 1 to 13. In the chemical structure of **3**, black numbers correspond to hydrogens, gray numbers correspond to carbons, blue numbers correspond to nitrogen, and red numbers correspond to oxygen. Intermolecular NOEs between r(CAG) and **3** and their corresponding distance restraints used for modeling are colored according to the atoms of **3**. The spectrum was acquired at 25 °C with 400 ms mixing time and 0.4 mM of RNA and 0.6 mM of **3**.

**Figure 4:**
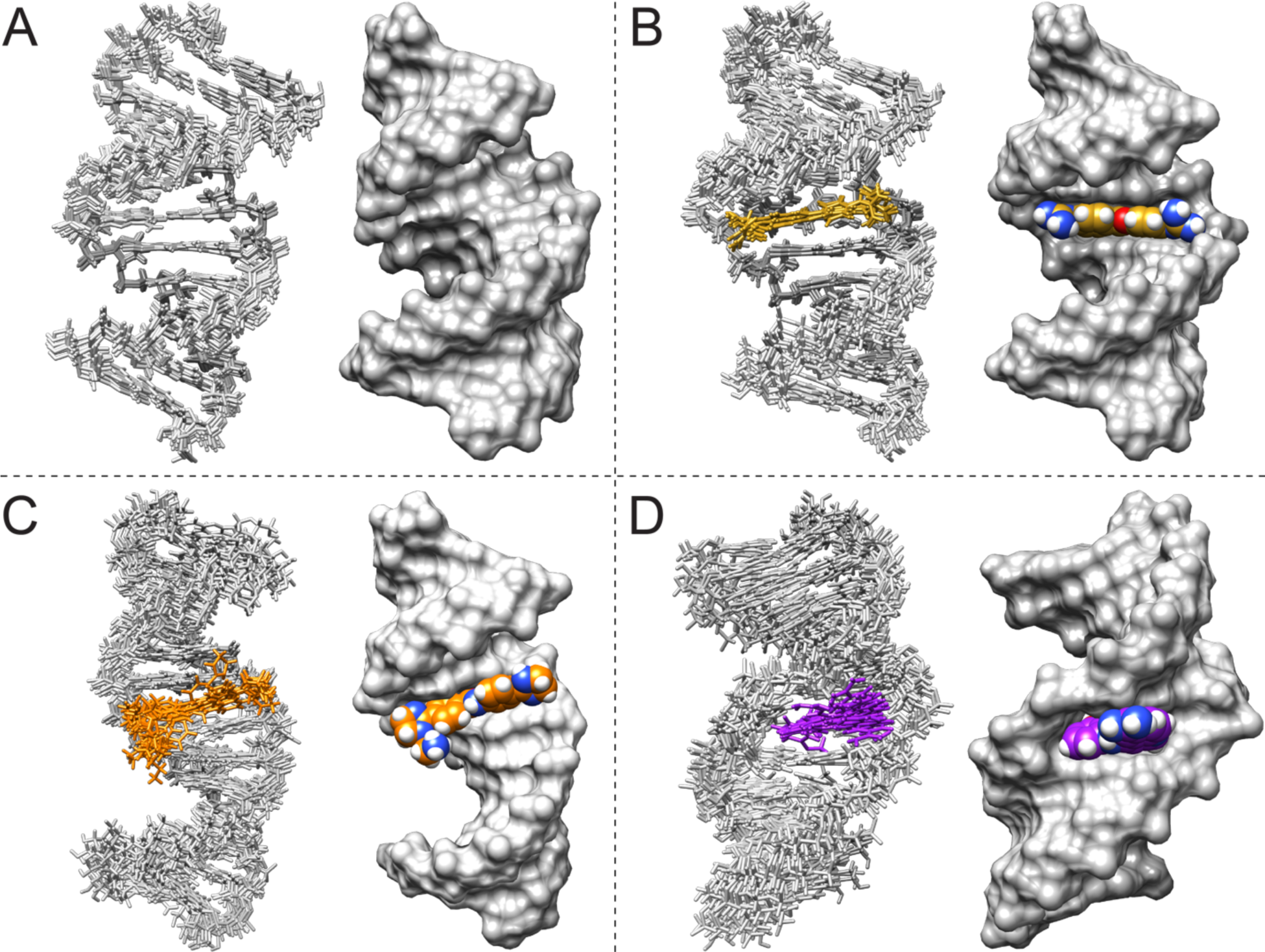
NMR solution structures of unbound and ligand-bound r(CAG) duplex model. (A) Structure of unbound r(CAG). (B) Structure of the r(CAG)-**1** complex. (C) Structure of the r(CAG)-**2** complex. (D) Structure of the r(CAG)-**3** complex. For each model, a surface representation (with the fewest distance restraint violations) and overlay of the 10 structures (stick representation) with the fewest distance restraint violations are shown.

**Figure 5:**
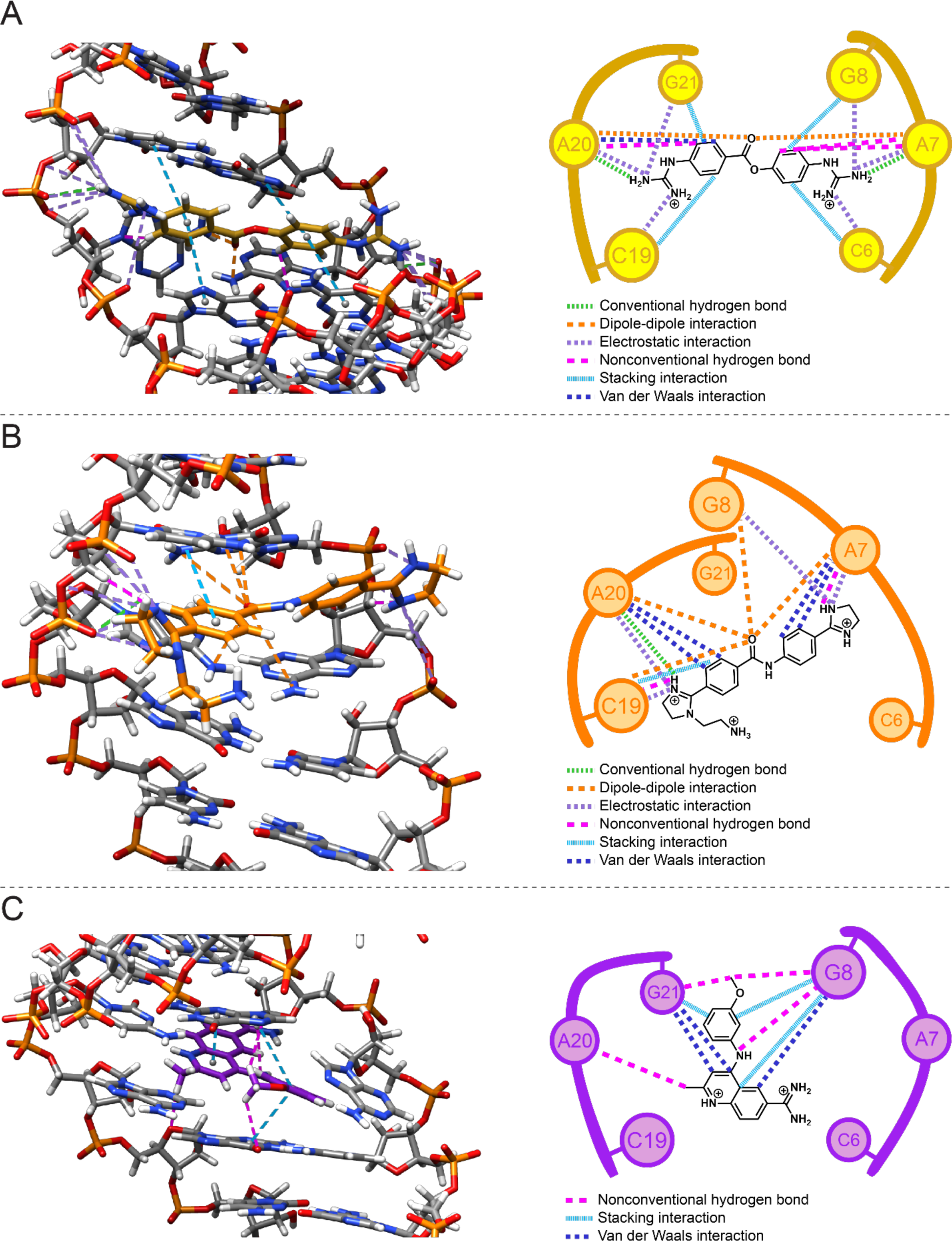
Schematic and 3D diagrams of ligand-RNA interactions in ligand-bound r(CAG) duplex model. (A) Interactions in the r(CAG)-**1** complex. (B) Interactions in the r(CAG)-**2** complex. (C) Interactions in the r(CAG)-**3** complex. For each model, a stick representation of the structure with the fewest distance restraint violations is shown. Light green dashed lines represent hydrogen bonding interactions, purple dashed lines represent electrostatic interactions, magenta dashed lines represent nonconventional hydrogen bonds, light blue dashed lines represent stacking interactions, and dark blue dashed lines represent van der Waals interactions. Van der Waals interactions are not shown in the 3D structures.

As summarized in **Table 1**, the binding of **1** induced various changes in the RNA’s structure, particularly the AA internal loop. In particular, a larger variation in the average helical rise and twist of the 20 **1**-bound r(CAG) structures indicated that ligand binding introduces additional dynamics to the RNA, while an increased bend angle (especially as indicated by the increase in roll values and the χ dihedral angles for the loop closing base pairs) and helix diameter are also indicative of differences in the overall helical structure (**Figure 6B**). These changes are consistent with shifting of imino proton resonances proximal to the AA mismatch upon titration of **1**. Further, the position and orientation of the aromatic rings of **1** nearby G8H1 could result in de-shielding and hence its change in chemical shift. Although the average C1′-C1′ distance for the AA mismatch upon **1**-binding was similar to that in the unbound structure, both the major and minor grooves of the 5’C<u>A</u>G/3’G<u>A</u>C motif were widened to accommodate ligand binding, the former to a greater degree. Unsurprisingly, the GC pairs adjacent to the AA mismatch were buckled more than other base pairs in the helix, and the AA mismatch had an average opening angle, average shear, and stretch distances typical for *trans* Watson-Crick/Watson-Crick AA pairs. of −0.5 ± 1.7 Å and −0.1 ± 0.8 Å, respectively. Additionally, the AA mismatch had a buckle of 14.3 ± 55.7° and propeller twist of 18.4 ± 51.7°, indicating that it is non-planar. Taken together, binding of **1** to the RNA induces distortions in the helix to accommodate the ligand-RNA interactions that stabilize the complex.

**Figure 6.**
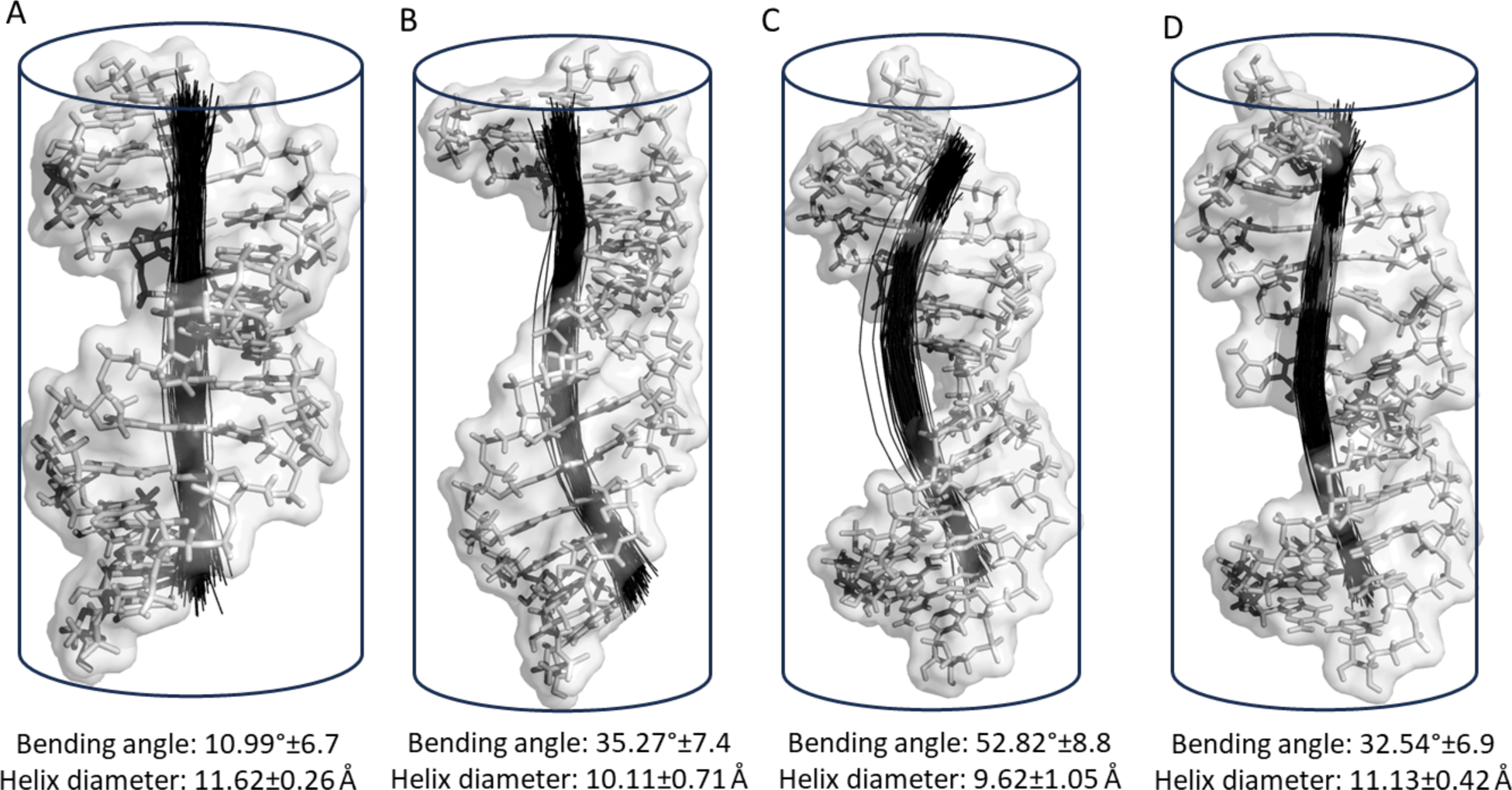
Model representations of the *apo* r(CAG) repeat and the repeats bound to 1, 2, and 3, showing bending of the helical axis and average diameter. (A) Model of the *apo*-r(CAG) repeat, showing no significant bending and a helix diameter consistent with A form RNA (1.2 nm). (B) The r(CAG) repeat bound to **1**, which causes helix bending and reduction of the average diameter of the structure. (C) Compound **2** causes major distortions of the helical axis and stretching of the structure by decreasing the helix diameter. (D) Although one of the A loop nucleotides is displaced as **3** intercalates in between the bases, there is no significant changes of the helix diameter as the main interactions are replaced by the intercalated compound. The change of the bending angle at the A/A loop is noteworthy.

### Comparison of the *apo* structure of r(CAG) repeat duplex and the repeat bound to 1

An examination of the *apo-*r(CAG) and r(CAG)-**1** revealed global changes in the helical structure and dynamics upon binding of **1**. This is evident from an increased bend angle, decreased helical diameter, and widening of the major groove, although minor groove widths did not significantly change. The AA pair in the r(CAG)-**1** structures was more dynamic than in the *apo*-r(CAG) structures, as indicated by large variations in buckle, opening, and propeller twist angles and χ-torsions. Loss of stacking of the AA mismatches with the closing GC pairs was indicated by ring overlap areas of 0.0 ± 0.0 Å^2^ and 0.4 ± 0.2 Å^2^ for the 5′CA/3′GA and 5′AC/3′AC steps, respectively. As with the *apo*-r(CAG) structures, the α-torsions of the 3′ guanines in r(CAG) took on unusual values (average of 138.1 ± 69.2°, with most values clustered around 160°). λ-angles for the AA mismatch in the r(CAG)-**1** structures also deviated from and were below the normal range for Watson-Crick pairs. Taken together, the *apo*-r(CAG) and r(CAG)-**1** structures retain a balance of hydrogen bonding and stacking interactions, but with additional dynamics and distortions in the r(CAG)-**1** structures.

### Comparison of the NMR-restrained structure of the r(CAG) repeat duplex bound to 1 and previously reported unrestrained structures

The P. Carloni group previously generated models of r(2×CAG) bound to **1** using unrestrained docking and MD simulations.(70) The group later reported a structure of furamidine bound to r(CAG) using MD simulations.(71) In all of the structures, the ligands bind in the major groove of the RNA, which was also widened, as observed with the **1**-r(CAG) NMR-refined structure reported herein. The buckle values of both closing GC pairs were negative in the Carloni group’s structures, whereas the buckle values of the closing GC pairs in the NMR-restrained structures alternated between positive and negative. Taken together, the compounds in these structures induced minor distortions to the helix within the r(CAG) motif and did not significantly distort the helical regions away from r(CAG). In the lowest free energy structure of r(2×CAG)-**1**, a π-stacking interaction as observed between one ring of **1** (β) and the A4 base and a T-shaped π-stacking interaction between the other ring (α) of **1** and the A14 base. In the r(CAG)-**1** NMR structures, **1** partially intercalated between the adjacent CG pairs and could be the source of the observed bending in the helix. Thus, the adenines did not form stacking interactions with **1**, and **1** instead formed stacking interactions with either or both of the closing GC pairs of 5’C<u>A</u>G/3’G<u>A</u>C.

In the Carloni group’s structures, salt bridges formed between each guanidine of **1** and a closing guanine of each r(CAG), and hydrogen bonds formed between the N6 atom (N23 in the Carloni group’s studies) O5′ of one closing guanosine, and between a carbonyl oxygen of **1** and a closing cytosine amino group. Similarly, the guanidines of **1** formed hydrogen bonds and electrostatic interactions with the phosphate backbone of one of the closing guanosines, in addition to an adenosine and both of the closing cytosines of r(CAG). Rather than interacting with the closing GC pair, the carbonyl oxygen of **1** instead formed electrostatic interactions with the A7 and A20 bases. As with the **1**-bound NMR structures, the mismatched adenines remained base paired in the Carloni group’s structures but with different configurations. Specifically, an A7N1-A14H2 hydrogen bond in the unbound state was replaced by an A7N1-A14H6 hydrogen bond upon binding of **1**. In summary, the unrestrained and NMR-restrained r(CAG)-**1** structures share structural features, with the compound adopting a similar orientation in the binding site and forming stacking, hydrogen bonding and electrostatic interactions with the RNA.

### Comparison of 1 bound to a r(CAG) repeat duplex model vs. a r(CUG) repeat duplex model

We reported the structure of **1** bound to a model of r(CUG) repeat expansions, as elucidated by NMR spectrometry and restrained MD.(23) Compound **1** in the r(CAG)-**1** and r(CUG)-**1** structures shared a similar binding mode, where the compound laid in the major groove, between the closing GC pairs. In these structures, the aromatic groups of **1** formed stacking interactions with the closing GC pairs and the guanidine tails formed hydrogen bonds and electrostatic interactions with the RNA backbone. While the mismatched adenines in the r(CAG)-**1** structures were displaced into the minor groove and form a base pair, in addition to hydrogen bonds with the closing GC pairs, the uracils in the r(CUG)-**1** structures were shifted away from each other in the base pair plane and formed stacking interactions with the closing GC pairs. Displacement of the mismatched adenines into the minor groove in the r(CAG)-**1** structures could allow **1** to lay closer to the minor groove of r(CAG) than r(CUG) in the r(CUG)-**1** structures, facilitating a greater number of stacking and electrostatic interactions with the RNA. Although the bend angles of the *apo*-r(CAG) (10.99 ± 6.7°) and -r(CUG) (11.88 ± 5.01°) structures were similar, binding of **1** to r(CAG) induced a significantly higher bend angle than observed in the r(CUG)-**1** structures.

A closer inspection of the r(CAG)-**1** and r(CUG)-**1** structures revealed that the helical rise and major groove widths were close to each other in the r(CAG)-**1** and r(CUG)-**1** structures, but the r(CAG)-**1** structures had greater variations in helical twist than the r(CUG)-**1** structures. Additionally, binding of **1** to the *apo* structures resulted in elongated C1′-C1′ distances in the mismatched bases. Nonplanarity of the AA mismatch was characterized by relatively large buckle and propeller twist angles and nonplanarity of the UU mismatch was characterized by a large stagger distance.(72) In summary, **1** adopts a similar binding mode in the r(CAG)-**1** and r(CUG)-**1** structures, but with differences in localized geometries and RNA-compound interactions.

### Structure of a r(CAG) repeat bound to 2

A total of 12 intermolecular NOEs were used as distance restraints to model the r(CAG)-**2** complex (**Tables 1 and S9**). The RMSD of all heavy atoms for the ensemble of 20 structures was 1.34 ± 0.32 Å (**Table 1**). Compound **2** laid in the major groove of the RNA, across the base pair plane, and formed stacking interactions with A7, A20, and one of the closing GC pairs (**Figures 5B**). In most of the structures, the mismatched adenine bases were stacked in the helix and formed a one hydrogen bond N6-H6···N1 *cis* Watson Crick/Watson Crick pair.(53) The **2** carbonyl oxygen was oriented into the major groove and formed dipole-dipole interactions with 1×1 nucleotide A/A internal loop and closing GC pairs while the amide nitrogen was oriented out of the major groove. Although no hydrogen bonds were observed in the reported structures, there were several hydrogen bond donors A20-NH2 and C19-NH2 near the carbonyl. The imidazoline(s) formed hydrogen bond(s) with the r(CAG) backbone or bases in 14 of the structures and electrostatic interactions with the RNA backbone, particularly at A7, G8, C19, and A20, in all 20 structures. Van der Waals interactions were formed between the **2** aromatic protons and adenine mismatch bases in all of the ensemble structures.

As with binding of **1** to r(CAG), binding of **2** to r(CAG) induced changes globally in the geometry of the helix (**Table 1**). Although the average helical rise and twist remained close to those of *apo*-r(CAG), the bend angle increased significantly, and helix diameter decreased significantly. Additionally, the major and minor grooves widened upon binding of **2**, consistent with shifting of the G5 and G8 imino resonances in the NMR titration. Additionally, the downfield shifting of G8H1 may be attributed to ring current effects of **2** as a result of the location of G8H1 relative to the aromatic rings of **2**. Locally, the average χ dihedral angles of the AA mismatch fell within the range expected for an *anti-anti* conformation. Average C1′-C1′, shear, stagger distances for the AA mismatch that were close to those in *apo*-r(CAG) were consistent with the AA mismatch retaining a similar hydrogen bonding pattern. However, large variations in the buckle, opening, and propeller twist angles indicate that it is dynamic. Taken together, binding of **2** to r(CAG) induces distortions in r(CAG).

### Comparison of r(CAG)-2 and the *apo*, r(CAG)-1, and r(CUG)-2 structures

Binding of **1**- and **2**-to r(CAG) and r(CUG) affected the overall helical geometries of the RNAs. The r(CAG)-**2** structures had a significantly higher bend angle than the r(CAG)-**1** and r(CUG)-**2** structures. While the helical rise and twist of the *apo*- and **1**-, and **2**-bound r(CAG) and r(CUG) structures were close to each other, significant variations in the helical twist were observed in the r(CAG)-**1** and r(CUG)-**2** structures. The major groove, which was 19-20 Å in the *apo*-r(CAG) and -r(CUG) structures, widened to a maximum of 21-23 Å in the **1**- and **2**-bound r(CAG) and -r(CUG) structures.

A closer inspection of the ligand binding sites in the **1**- and **2**-bound r(CAG) and r(CUG) structures revealed some similarities in the binding modes of the compounds. As in the r(CAG)-**1** and r(CUG)-**2** structures, the r(CAG)-**2** structure was stabilized by hydrogen bonds and electrostatic interactions between the guanidines or imidazolines and the phosphate backbone of the RNA. The mismatched adenines in the r(CAG)-**1** and -**2** structures formed stable base pairs, as indicated by average hydrogen bond counts of 2.0 ± 0.0 and 1.0 ± 0.0, respectively. On the contrary, base pairing of the mismatched uracils in r(CUG)-**1** was largely disrupted, as indicated by an average hydrogen bond count of 0.2 ± 0.4. The same uracils of r(CUG)-**2** alternated between one and two-hydrogen bond base pairs and between *cis*- and *trans*-Watson Crick/Watson Crick base pairs in the minor groove, as indicated by an average hydrogen bond count of 1.6 ± 0.5. This may result from the lower conformational stability of UU pairs compared to AA pairs, particularly in these ligand-bound structures, as indicated by greater variations in the hydrogen bond counts of the UU pairs. As observed in the r(CUG)-**1** structures, the compound in the r(CAG)-**2** structures laid farther in the major groove than **1** in the r(CAG)-**1** structures or **2** in the r(CUG)-**2** structures. This accommodates stacking of the mismatched bases in the helix, as indicated by ring overlap areas of 1.4 ± 0.8 Å^2^ and 1.6 ± 0.8 Å^2^ for the 5′CA/3′GA and 5′AC/3′AC steps, respectively in the r(CAG)-**2** structures 1.5 ± 1.0 Å^2^ and 2.0 ± 0.9 Å^2^ for the 5′CU/3′GU and 5′UC/3′UC steps, respectively in the r(CUG)-**1** structures. Such stacking interactions occurred at the expense of stacking interactions between the compounds and RNA. However, unlike in the r(CUG)-**1** structure, the mismatched bases in the r(CAG)-**2** structures remained base paired upon binding of the compound. Additionally, the C1′-C1′ distance of the AA mismatch in r(CAG)-**2** structures was close to that in the *apo*-r(CAG) structures, whereas the C1′-C1′ distance of the UU mismatch decreased in the r(CUG)-**2** structures upon binding of compounds. In summary, binding of **1** and **2** to r(CAG) introduced new hydrogen bonding and stacking interactions to the helices without disrupting base pairing between the mismatched adenines. These results are in line with the single molecule observations, *vide infra*, where no significant mechanical changes were observed upon ligand interactions indicating the preservation of base stacking.

### NMR analysis of a r(CAG) repeat bound to 3

The r(CAG) duplex-**3** structures were modeled using 22 intermolecular RNA-compound NOEs. In a 400 ms spectrum acquired at 35 °C, NOEs were observed from the methyl group and aromatic protons of the methoxybenzene moiety to C6H2′, A7H1′, and G8H1′ (**Figure 3**). On the quinoline ring, NOEs were observed from H2 to C6H2′ and G8H1′, H4 to G8H1′, and the methyl group to C6H1′, C6H2′, A7H1′, and G8H1′. In a 95% H_2_O spectrum, the G8H1 resonance was not observed either because of broadening or overlap with G11H1 (**Figure S12**). These NOEs suggest stacking of the compound between the closing GC pairs of AA internal loop. Additional NOEs between the terminal nucleotides and **3** arise from end stacking of **3** as a result of the high concentration of **3** relative to the RNA.

### Structure of a r(CAG) repeat bound to 3

In the final ensemble of 20 structures with the fewest distance restraint violations, the RMSD of all heavy atoms was 1.75 ± 0.52 Å. The **3**-RNA complex adopted an A-form geometry, with the compound in the minor groove. The quinoline and methoxybenzene groups of the compound formed stacking interactions with the closing GC pairs, displacing the adenines into the major groove, where they did not base pair. A7 was in the *syn* conformation (O4′-C1′-N9-C4 from −90° to 90°) in 17 of the 20 structures, and A20 was in the *syn* conformation in one of the structures. The methyl groups of **3** were oriented towards the major groove and formed nonconventional hydrogen bonds with G8, A20, and G21. Additionally, N3 of **3** formed a nonconventional hydrogen bond with G8 and the C8, C9 and C18 aromatic protons of **3** interacted with G8 and G21 via van der Waals interactions (**Figure 5C**).

Binding of **3** to r(CAG) resulted in changes in the RNA structure (**Table 1**). Large variations in helical rise and twist indicate that the complex is dynamic. Decreases in the helical rise, twist, and diameter and increases in bend angle and widening of the major and minor groove widths indicate changes in the overall geometry of the helix. Within the AA mismatch, increases in C1′-C1′ distance and opening angle and decreases in shear and stretch distances are consistent with disruption of hydrogen bonding. The disruption of the AA base pair resulted in dynamics within the r(CAG) motif, on the basis of large variations in the buckle and propeller twist angles and stagger distance of the AA mismatch. These observations are also consistent with shifting of imino resonances, particularly around the r(CAG) motif, in the NMR titration experiments. In summary, binding of **3** to r(CAG) results in loss of base pairing within r(CAG) and alters local structure within and global structure of the helix.

### Comparison of the r(CAG)-3 structure with the *apo*, r(CUG)-3, and tau RNA-3 structures

In NMR-refined structures of **3** bound to a tau splicing regulatory element RNA,(24) **3** bound to r(CAG), and **3** bound to r(CUG),(23) **3** formed stacking interactions with the RNAs and disrupted base stacking of loop bases. However, the stacking configuration of **3** with the RNA in the **3**-tau RNA structures differed from the **3**-r(CAG) structures, with both rings of the quinoline but not the methoxybenzene stacking with the closing GC pair. At the A-bulge site of the unbound tau RNA, the unpaired adenine was stacked between its closing GC pairs, inducing a kink in the opposite strand of the helix.(73) Stacking of **3** in the tau RNA displaces the A-bulge from the helix into the minor groove, increasing coaxial stacking in the opposite strand and bringing the helix to a more A-form like conformation. The reduction in kink was evident from a decrease in the tilt of the 5′GC/3′CG step. Binding of **3** to the major groove of the tau RNA also facilitated van der Waals and/or nonconventional hydrogen bonding interactions between the methoxybenzene group and the residues adjacent to the kink. Thus, binding of **3** to the tau RNA causes the helix to adopt an A-form like conformation. In contrast to the tau A-bulge site, the r(CAG) motif did not contain a kink as a recognition element for the compound. Thus, binding of **3** in the major groove of 5’C<u>A</u>G/3’G<u>A</u>C was not necessary to stabilize a kink motif. Instead, the complex was stabilized by stacking of the compound between the closing GC pairs to facilitate continuous stacking of the helix. The continuous stacking within the **3**-bound r(CAG) motif was indicated by a small average tilt value for the 5′C6G21/3′G8C19 step, which excludes the unstacked adenines of the 1×1 A/A loop. Stacking interactions formed between the adenine bases and the cytosines of the closing GC pairs in the **3**-r(CAG) structures, as indicated by the ring overlap areas of the 5′C6A7/3′G21A20 and 5′A7G8/3′A20C19 steps. In comparison, in the r(CUG)-**3** structure, where U7 and U20 were not stacked, there was no ring overlap areas of the 5′C6U7/3′G21U20 and 5′U7G8/3′U20C19 steps. Thus, the stacking interactions of the mismatched adenines with the closing GC pairs in the major groove may favor binding of **3** in the minor groove of r(CAG).

In the tau RNA-**3** and r(CUG)-**3** structures, the amidine of **3** was in the major groove of the RNA and N2 and N3 of the compound laid between the closing GC bases and, in the **3**-tau RNA structure, close to the minor groove A-bulge. Thus, they are better positioned to form hydrogen bonds with the RNA. In contrast, the positioning of these hydrogen bonding groups in the minor groove of r(CAG) prevented such interactions with the RNA. In the tau RNA-**3**, r(CAG)-**3**, and r(CUG)-**3** structures, the C15 methyl group formed van der Waals and/or nonconventional hydrogen bonding interactions with the RNA backbone. The bulky C7 methyl group interacts with the RNA in the major groove of the r(CAG)-**3** structures and minor groove of the r(CUG)-**3** structures but did not interact with the RNA the tau-**3** RNA structures. Thus, the positioning of **3** in the minor groove of the r(CAG) motif facilitated nonconventional hydrogen bonding, stacking, and van der Waals interactions of **3** with the RNA in place of conventional hydrogen bonding interactions.

### Comparison of r(CAG)-1, -2, and -3 structures

In the r(CAG)-**1** and -**2** structures, **1** and **2** displaced the adenines into the minor groove with retention of hydrogen bonding between adenines. In the r(CAG)-**1** structures, the A7 and A20 bases were rotated so that their amino groups were in close proximity to hydrogen bond with G21- and G8-N3, respectively, on the opposite strands. For the r(CAG)-**2** structures, the adenines were partially stacked on the adjacent CG base pair and the adenine to G-N3 hydrogen bonds were not observed. In both r(CAG)-**1** and -**2**, the carbonyl was oriented towards the minor groove. For the r(CAG)-**1** structures, the carbonyl exhibited hydrogen bonds to the amino group of the displaced A7. The carbonyl in r(CAG)-**2** did not hydrogen bond to the displaced adenines in the structure with the fewest restraint violations. However, in five of the r(CAG)-**2** structures, the carbonyl did form hydrogen bonds with the A7 and/or A20 amino groups. In the r(CAG)-**3** structures, **3** was intercalated between the closing CG pairs, displacing the adenines into the major groove with one adenine adopting a *syn* conformation. The displaced adenines were partially stacked on the adjacent cytosines of the r(CAG) motif and were no longer hydrogen bonded.

All three compounds contain positively charged groups. The positively charged guanidines and imidazolines of **1** and **2**, respectively, had hydrogen bonding and electrostatic interactions with the negatively charged phosphates on opposite strands. In the case of r(CAG)-**1**, the positively charged guanidines had electrostatic interactions with the A7 and A20 phosphates. In r(CAG)-**2**, the imidazolines had electrostatic interactions with the A20 and G8 phosphates. These interactions add to specificity in binding of **1** and **2** to r(CAG). In contrast, the positively charged amidine of **3** protruded out of the minor groove with little to no interactions with phosphates in the RNA backbone.(74) In summary, **1** and **2** formed significant stacking, hydrogen bonding and electrostatic interactions with the r(CAG) and retained the AA mismatch hydrogen bonding. For r(CAG)-**3**, binding was dominated by stacking interactions with few observed hydrogen bonds and electrostatic interactions. Additionally, the bulky methyl substituents of **3** filled the binding cavity formed by the displaced adenines. The displaced adenines were no longer hydrogen bonded therefore making the loop more susceptible to unfolding.

Zooming in on the binding interactions of the first five lowest energy structures from the NMR ensemble for **1**, **2**, and **3**, it was observed that a combination of multiple hydrogen bonds and stacking interactions (**Supplemental Data Set 1**) are the main stabilizing forces of the compounds bound states (**Figures 5 and S14**). Compounds **1** and **2** can both form stacking interactions with the closing GC base pairs of the 1×1 nucleotide A/A internal loop. Nonconventional hydrogen bonds are also observed forming between **1** and A7/A20, between **2** and A7/C19 and between **3** and G8/G21/A10 (**Figure 5**). Although **1** and **2** show stacking interactions with the immediate neighbors of the 1×1 nucleotide A/A internal loop but the degree of overlap is not as significant as **3** as the later interacts in an intercalating way optimizing the degree of overlap between flat surfaces hence enhancing the π-π interactions (**Figure S13**). Shape complementarity between **1**, **2**, and **3** and r(CAG) is also evident from the large bend angles of these ligand-bound structures and results in structural specificity. In summary, **1 – 3** contain a variety of functional groups that facilitate selective binding to their target RNAs.

To investigate the global differences in the architectures of the four structures we completed the calculation of global formations by running 10 µsec long MD simulations. Conformational flexibility or deformability of RNA is important for its function.(75) These deformations specially affect binding affinity of proteins through shape-dependent recognition mechanism.(76) In order to measure the global deformation of the RNA before and after ligand binding, the overall bend angle (global curvature) was calculated using *Curves+*.(45) The global curvilinear helical axis was calculated from the average local axes applying a polynomial smoothing. We used a 10 µsec long MD trajectory, ignoring the first 2 µsec of the simulations, to calculate the overall bending angle.

The *apo* form of the r(CAG) repeat duplex model had a bend angle of 10.99° with no structural distortions as expected for an A-form RNA (77) (**Figure 6A**). Subsequent bending measurements of the RNA bound to **1** showed an increase in bending angle compared to *apo* form (35.27°) (**Figure 6B**). The most significant distortions, however, were caused by **2**, with a global bending measured at 52.82° (**Figure 6C**). Among the studied compounds, **3** was the only compound that intercalates into the binding pocket provided by the displaced A/A nucleotides. Notably, the induced global bending by **3** (32.54°) was not as drastic as the minor groove binders, implying some of the lost interactions due to the displaced A/A nucleotides are replaced by the inserted compound (**Figure 6D**). Compound **2** also had similar electrostatic interactions with the RNA compared to **1**, where such interactions extend across the A/A loop. The overall conformational changes induced by these compounds have important implications for RNA-protein interaction as manifested by single molecule studies.

### Small molecule binding alters the dynamics of the 1×1 nucleotide A/A internal loop

To understand the effect of the compound binding on the dynamics of the 1×1 nucleotide A/A internal loop, a 10 µsec long MD simulations with NMR restraints was performed and a 2D PMF (potential of mean force) plot was created based on the PC1 (principal component 1) and PC2 (principal component 2) of the 1×1 nucleotide A/A internal loop and the most immediate neighboring base pairs on both sides. It was observed that the *apo* form of the r(CAG) duplex RNA adopted two minima along the free energy landscape, corresponding to one and two hydrogen bond states (**Figure S14**). Subsequent cluster analysis showed two major populations (State 1 and 2), each forming one hydrogen bond in the 1×1 nucleotide A/A internal loop; multiple sugar phosphate backbone hydrogen bonds (sugar-sugar (S-S) or sugar to phosphate (S-P)) in State 2 were observed (**Figure S14A**). These classes of hydrogen bonds have been observed in many RNA structural motifs.(78)

PMF calculations of the r(CAG)-**1** complex revealed one minimum (**Figure S14B**) corresponding to a structure with no hydrogen bonds in the 1×1 nucleotide A/A internal loop but multiple backbone hydrogen bonds, both observations consistent with NMR structures. These data indicate that the bound compound has restricted the conformational changes around the 1×1 nucleotide A/A internal loop as compared to the *apo* form (**Figure S14B; Right**). Compound **2** caused more drastic structural changes, in agreement with the NMR ensemble, creating three distinct minima (States 1 – 3) in the free energy landscape (**Figure S14C**). States 1 and 3 formed no hydrogen bonds in the 1×1 nucleotide A/A internal loop while State 2 formed one hydrogen bond. Multiple sugar-sugar and sugar-phosphate backbone hydrogen bonds were also observed in all three states, with two bifurcated hydrogen bonds in state 2 (**Figure S14C**).

As **3** intercalates between the adenosine loop nucleotides and displaces them from the helical axis, it is not surprising that differences were observed in PMF calculations, which afforded three minima with one distinct minimum corresponding to State 1 (**Figure S14D**). The two other minima were not as highly populated as the first minimum. In State 1, the A nucleotides remained within the helical axis although hydrogen bonds formed between them were broken. In States 2 and 3, one adenosine was displaced from the helical axis while the other remained within the helical axis. States 2 and 3 also formed multiple sugar-sugar and sugar-phosphate hydrogen bonds (**Figure S14D;** States 2 and 3).

### Comparison of other small molecules bound to r(CAG) repeats

The X-ray crystal structures of other small molecules in complex with a tandem r(CAG) repeat [r(2×CAG)] have been reported, including cyclic *bis*-naphthyridines (NAs) and cyclic mismatch-binding ligands (CMBLs), which adopt similar binding modes.(19) The CMBLs intercalate with the adenines and closing GC pairs and disrupt base pairing of the 1×1 A/A nucleobases. Each naphthyridine moiety forms a base pair with an adenine via two hydrogen bonds while the linkers interact with the adenines via water-mediated hydrogen bonds. The pseudo-canonical base pairs are stacked in the helix and displace the central cytosines and guanines into the minor groove, where they stack but do not form base pairs. In common with the naphthyridine-bound r(2×CAG) structures, **1** – **3** primarily recognize 1×1 A/A loops via stacking interactions.

Although intercalators might bind nonspecifically with DNAs and RNAs,(79) target discrimination may be achieved by adding hydrogen bonding groups to the compounds.(19,80) Alternatively, filling the binding cavity with functional groups that form other types of interactions can result in entropic gains by displacing water molecules from the binding cavity and into bulk water.(80,81) For target specificity, the NAs and CMBLs form a pattern of hydrogen bonds with the r(CAG) motifs, influenced by the linker length and geometry. Rather than relying on hydrogen bonding interactions for target specificity, **1** – **3** use a combination of hydrogen bonding, electrostatic, dipole-dipole, and van der Waals interactions.

### Single molecule studies of r(CAG) vs. r(CUG)

The NMR-restrained structures along with MD and PMF calculations to explore dynamics suggest that the small molecules might affect MBNL1 binding differently. As **1** and **2** binding helps to stabilize the loop, we hypothesized that ligand binding might inhibit the binding of MBNL-1. In contrast, **3** displaced the loop adenines, which were no longer hydrogen bonded, perhaps making the loop more susceptible to unfolding and MBNL-1 binding. We therefore explored the binding of the three small molecules to both r(CAG) repeats using magnetic force spectroscopy (MFS) single molecule studies. In MFS, RNA molecules are tethered to a flow cell surface and attached to micron-scale paramagnetic beads. They are then subjected to a controllable magnetic force while the beads’ vertical positions are tracked with high precision. The platform thus provides information about the dynamics of molecular unfolding/refolding and allows the impact of ligand binding to be studied. In an experiment set up known as a force ramp, the magnetic force is slowly increased until the RNA becomes unfolded. This is detected as a sudden change in the bead position. Then, the applied force is gradually reduced allowing the RNA to refold. This non-destructive cyclical process is repeated up to 100 times across hundreds of individual molecules, to maximize data acquisition.

Using r(CAG)_21_ in ramp experiments, we observed that the RNA unfolds and refolds without any hysteresis (**Figures 7A & S15A**). Hysteresis indicates that some energy is dissipated (lost) in the process of stretching and relaxing of the molecule. This can be due to conformational changes that do not revert completely.(82) This observation differed from the r(CUG)_21_, where a small but consistent hysteresis has been observed (**Figure 7A**). To investigate further, we compared the distribution of the normalized unfolding and refolding forces of the two RNAs across over 300 molecules. For r(CAG)_21_, the unfolding and refolding forces overlapped (p = 0.33) and occurred at around 11 pN. In contrast, r(CUG)_21_ RNA unfolded at around 14 pN and refolded at around 13.5 pN, which confirmed the observation of a hysteresis (p = 0.00058) (**Figure S15B**). Previous studies have shown that RNAs with multiple r(CUG) repeats are thermodynamically more stable than RNAs with an equal number of r(CAG) repeats (Δ*G*^37°^_*diff*_ ∼1.22 kcal/mol).(83) Therefore, our observation that the r(CUG)_21_ RNA unfolds at a higher forces matches with the thermodynamic studies.

**Figure 7.**
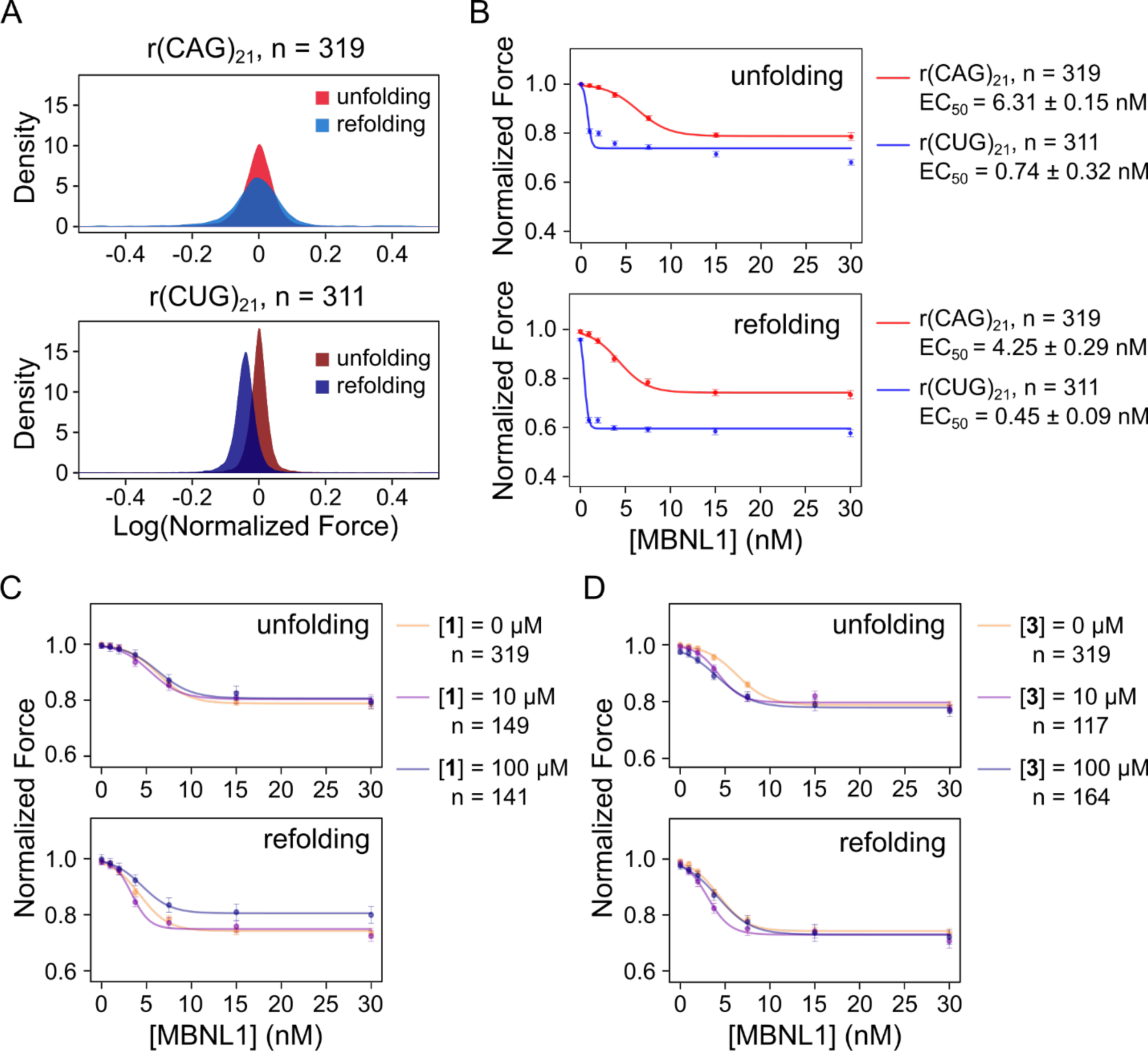
The effect of compounds on r(CAG)21 RNAs obtained by magnetic force microscopy. (A) Normalized force distribution for r(CAG)_21_ (top) and r(CUG)_21_ (bottom) RNAs in force ramp experiments. The numbers of molecules analyzed (n) are shown. (B) The median normalized unfolding (left) and refolding (right) force with increasing concentration of MBNL1 proteins, comparing r(CAG)_21_ and r(CUG)_21_ RNAs. (C-E) The median normalized unfolding (left) and refolding (right) force with increasing concentration of MBNL1 protein in the presence of 0/10/100 µM of **1** (C) and **3** (D). The numbers of molecules analyzed in each condition are shown, and the data are shown as median force +/-SEM.

MBNL1 protein binds to RNA r(CAG) repeats as well as r(CUG) repeats, albeit with a lower affinity to the former(84,85), and formation of r(CUG)^exp^- and r(CAG)^exp^-complexes cause disease.(86) As aforementioned, **1** inhibits formation of the r(CAG)^exp^-MBNL1 complex and rescues MBNL1-associated splicing defects in patient-derived cells.(18) Therefore, the effects of MBNL1 binding on the two RNAs in the force ramp assay were also studied.

MBNL1 protein binding caused a decrease in the median unfolding and refolding force in both RNA repeats (**Figure 7B, S15C**). In addition, for a given single RNA molecule, it could be observed that MBNL1 prevented the RNA from unfolding and refolding for some of the force cycles to which it was subjected, thereby causing a decrease in the unfolding and refolding probabilities (**Figure S15D**). These observations suggest that MBNL1 preferentially binds to single-stranded r(CAG) or r(CUG) repeats when they are unfolded, and therefore decreases the refolding force. In subsequent cycles, this partially folded secondary structure would require less force to unfold, causing a global decrease in both the unfolding and refolding force and probability. The observation of preference of MBNL1 for single-stranded RNA is in agreement with structures of MBNL1-RNA complexes where MBNL1 binding causes partial unwinding of the RNA (87-89) as well as *in vitro* footprinting assays.(90)

While the same trends in the data were observed for both RNAs upon MBNL1 binding, a difference in their binding affinities was observed (**Figure 7B**), as expected. At the lowest MBNL1 concentration tested (0.95 nM), the protein had only a negligible effect on r(CAG)_21_, while having a noticeable impact on the r(CUG)_21_ unfolding and refolding forces (0.994+/- 0.00665 and 0.0981 +/- 0.0106 for normalized unfolding and refolding force, respectively, for r(CAG)_21_, and 0.807 +/- 0.00989 and 0.930 +/- 0.0106 for normalized unfolding and refolding force, respectively, for r(CUG)_21_). (**Figure S15C**). The unfolding probability (*i.e.*, the fraction of cycles showing an unfolding/refolding event) also decreased slightly more at lower concentration of MBNL1 protein, suggesting that at these concentrations, more MBNL1 protein binds to r(CUG)_21_ compared to r(CAG)_21_. At higher concentrations, however, the unfolding probability reduced to a similar extent for both RNAs, indicating that there is no difference in behavior once MBNL1 protein has saturated the RNA (**Figure S15D**).

This difference in affinity was confirmed by a second experiment, referred to as stepped force in which the force is held constant for a fixed period before being increased stepwise until the RNA structure remains unfolded. During a given force step, the RNA may transition between different states and the amplitude of the transition can be assessed from the variance in bead displacement. As the force is increased stepwise, there is a progressive increase in the displacement variance due to the increasing number of repeats that unfold, until a maximum is attained, after which the displacement variance decreases until the molecules remain fully unfolded. For both r(CUG)_21_ and r(CAG)_21_, increasing MBNL1 protein concentration progressively decreased the maximum transition variance, suggesting that the bound protein reduces the available length of RNA that can refold. MBNL1 protein also slightly reduced the force at which the maximum variance occurred, indicating that protein binding decreases the stability of the RNA structure. At higher concentrations, MBNL1 protein almost completely abolished the transition variance, likely due to the protein binding to all available sites and preventing the RNA from forming a structure (**Figure S15E**). The concentration of MBNL1 protein where changes in the force and height of maximum concentration were first observed were different for the two RNAs, however, 0.95 nM and 15 nM for r(CUG)_21_ and r(CAG)_21_, respectively. Further, saturation of MBNL1 binding was observed at a lower concentration for r(CUG)_21_ than for r(CAG)_21_, 3.75 nM and 30 nM, respectively (**Figure S15E**). Collectively, these stepped force studies are in agreement with the observations from force ramped assays, showing that MBNL1 protein bound to r(CUG)_21_ with higher affinity than it did to r(CAG)_21_ in MFS single molecule studies.

### Effects of 1-3 on RNA repeat stability

After establishing the differences between r(CUG)_21_ and r(CAG)_21_ folding and unfolding under mechanical force in the presence and absence of MBNL1, we next studied the effect of compound binding to r(CAG)_21_. No effect on the unfolding and refolding force (force ramp experiments) was observed for any compound, indicating that the binding of **1** – **3** did not change the stability of the RNA’s structure in a way which can be captured by MFS single molecule study (**Figure S16**). The unfolding and refolding probabilities also showed no drastic changes with increasing concentrations of **1** (**Figure S16A**), confirming that **1** binding to r(CAG)_21_ did not impact unfolding and refolding. Increasing concentrations of **2** slightly decreased the unfolding and refolding probabilities (**Figure S16B**), indicating that **2** might either prevent the RNA from refolding properly so that it remains single stranded or prevent full unfolding. For **3**, we also observed a dose-dependent reduction of unfolding and refolding probability, with the effect more marked for refolding, although the effect is modest (**Figure S16C**). These data suggest that **3** might prevent the RNA from refolding properly so that it remained single stranded.

### Effect of compounds 1-3 on MBNL1 protein binding to r(CAG)_21_

To assess whether compound binding affects the binding of MBNL1 protein to r(CAG)_21_, experiments were performed at a constant concentration of each of the three compounds in the presence of increasing concentrations of MBNL1.

*Compound **1**.* Small molecule **1** was tested in force ramp experiments at concentrations of 10 and 100 µM and varying concentrations of MBNL1 (0.95 nM to 30 nM; **Figure 7C**). At both concentrations, no discernable effect on the reduction of the median normalized unfolding force of r(CAG)_21_ as a function of MBNL1 protein concentration was observed. However, at the higher concentration, **1** seemed to partially counteract MBNL1’s effect of reducing the force at which the RNA repeats refolded. This effect was apparent even at saturating concentrations of MBNL1 (>10 nM), possibly due to less of the protein binding to the RNA, allowing it to refold at a higher force (**Figure 7C**). Further, **1** did not have any significant effect on the unfolding and refolding probabilities, suggesting that it counters MBNL1’s destabilization of the structure, not by acting on the rate of binding of the MBNL1 protein, but rather by decreasing the number of available r(CAG) repeats for MBNL1 to bind (**Figure S17A**). Thus, the NMR structures elucidated herein and the observations from single molecule studies suggest that binding of small molecules to the internal loops reduces their accessibility to MBNL1 binding. Direct binding of **1** to the r(CAG) repeats is consistent with changes in the G8 imino proton resonance in 1D NMR spectra upon addition of **1** to the r(CAG) duplex model and with NOEs between the compound and 5’C<u>A</u>G/3’G<u>A</u>C residues in 2D NMR spectra. This observation provides a mechanistic view on how **1** prevents binding of the MBNL1 to repeat expansions and hence improve splicing defects.

*Compound **2**.* Likewise using force ramps experiments, we observed that 100 µM of **2** (but not at 10 µM) enhanced the effect of MBNL1 protein binding as evidenced by the reduction in the median unfolding force in response to MBNL1 protein (**Figure S17B**) and analysis of the unfolding probability (**Figure S17C**). Collectively, the unfolding data suggest a mechanism whereby **2** increases the binding rate of MBNL1 protein to the RNA structure. For the refolding of r(CAG)_21_, however, **2** showed variable effects. Specifically, while **2** did not alter the refolding probability (**Figure S17C**), it had contradictory impacts on the refolding force, diminishing MBLN1 binding at 10 µM but enhancing it at 100 µM (**Figure S17B**). The larger bend in the helix induced by **2** than **1** would cause the RNA to become more distorted when multiple r(CAG) sites are bound to **2**.

*Compound **3**.* In force ramp experiments, **3** appeared to enhance the effect of MBNL1 protein binding during RNA unfolding but had a less consistent impact on refolding (**Figure 7D**). At MBNL1 protein concentrations below 10 nM, 10 µM of **3** enhanced the reduction in the unfolding force, although the minimum unfolding force reached at saturating protein concentrations was unchanged. These observations indicate that a lower MBNL1 protein concentration is required to reach binding saturation, and therefore **3** enhances MBNL1 binding to r(CAG)_21_. Increasing the concentration of **3** (to 100 µM) did not further enhance the effect of MBNL1 protein binding to the RNA (**Figure 7D**). At lower MBNL1 concentrations, the unfolding probability also showed a slightly larger reduction, possibly due to more MBNL1 protein binding at a given concentration (**Figure S17D**). While the effect of MBNL1 on refolding of the RNA was enhanced by **3** at 10 µM, this effect was not observed at 100 µM (**Figure 7D**). Interestingly, **3** decreased the minimum refolding probability reached (**Figure S17D**), indicating that the compound may stabilize the ternary complex, therefore increasing the number of force cycles with bound protein where the RNA could not be refolded properly. Alternatively, **3** could also diminish RNA folding (as seen by the decrease of the folding probability in the presence of **3** alone). Therefore, more RNA molecules would remain single stranded, allowing more MBNL1 protein to bind.

Overall, these observations confirm that **1** prevents the MBNL1 protein from binding to r(CAG)_21_. Since **1** has a greater effect on RNA refolding, **1** binding could compete or inhibit the binding of MBNL1 protein. The effect of **2** was more variable in these experiments and therefore difficult to interpret. In contrast, **3** seemed to enhance MBNL1 protein binding to r(CAG)_21_, mostly likely at the unfolding stage. One hypothesis is that the binding of **3** displaces the adenines from the binding site, allowing the RNA to be more easily unfolded, and impeding refolding which is supported by the mode of binding of **3** obtained from NMR structure. Therefore, the RNA would be more likely to be single stranded and allow more/faster MBNL1 protein binding.

### Outlook

Herein, we report the structure of a model of r(CAG) repeats bounds to three small molecules. The atomistic details describing how each ligand interacts with the RNA enable optimization by structure-based design and iterative medicinal chemistry (91) and for virtual screening campaigns. The success of both structure-based design and virtual screening depends on the availability of high-resolution 3D structures and obtaining such structures is especially challenging for RNA due to its highly dynamic nature.(92-94) Exploiting conformational dynamics, however, could also be a source of specificity for RNA targets.(95) The alternate structures present in an ensemble provide binding pockets which might be not be available in a single structure. Indeed, ensemble molecular docking of RNA has been developed to design small molecules and improve hit rates (96) as well as to reduce the number of false positives in virtual screens.(97)

Our complementary single molecule studies also provide mechanical insights on how small molecules can interact with RNA repeat expansions and inhibit binding of proteins such as MBNL1. Overall, the dynamical and mechanical overview provided for the first time for r(CAG) model bound to small molecules provides the fundamentals of design principles of small molecules that can rescue MBNL1 activity as a potential therapeutic approach.

## SUPPORTING INFORMATION AVAILABLE

(I) H6/H8-H1′ region of a 2D ^1^H-^1^H NOESY spectrum of unbound r(CAG). (II) Imino proton region of a 2D ^1^H-^1^H NOESY spectrum of unbound r(CAG) and r(CAG)-**1**, r(CAG)-**2**, and r(CAG)-**3** complexes. (III) 1D ^1^H and WaterLOGSY NMR spectra of **1**, **2**, and **3**. (IV) 1D ^1^H imino region titration with **1**, **2**, and **3**. (V) Structure of the unbound r(CAG) motif. (VI) Comparison between r(CAG)_21_ and r(CUG)_21_ RNA behavior by magnetic force microscopy. (VII) The effect of different compounds binding to the r(CAG)_21_ RNA in ramp and force step experiments. (VIII) The effects of compounds **1**, **2**, and **3** on MBNL1 protein binding. (IX) ^1^H NMR chemical shifts of the unbound r(CAG) duplex and r(CAG)-**1**, r(CAG)-**2**, and r(CAG)-**3** complexes. (X) NOE restraints used for modeling of the unbound r(CAG) duplex. (XI) Dihedral restraints used for modeling of the unbound r(CAG) duplex. (XII) NOE restraints used for modeling of the r(CAG)-**1**, r(CAG)-**2**, and r(CAG)-**3** complexes. (XIII) Summary of select helical and base pair parameters for *apo*- and ligand-bound RNA constructs (**Supplemental Data Set S1**).

## FUNDS

This work was supported by the National Institutes of Health grant R01 CA249180 (to M.D.D.), the American Chemical Society Division of Medicinal Chemistry and Muscular Dystrophy Association (MDA), Development Grant 963835 (to A.T.) and the Huntington Disease Society of America and the National Ataxia Foundation (to J.L.C.).

### Notes

The authors declare the following competing financial interest(s): M.D.D. is a founder of Expansion Therapeutics.

## ABBREVIATIONS

1D: one-dimensional
2D: two-dimensional
CMBL: cyclic mismatch-binding ligands
COSY: correlation spectroscopy
DM1: myotonic dystrophy type 1
HD: Huntington’s Disease
MAPT: microtubule associated protein tau
MBNL1: muscleblind-like 1 protein
MD: molecular dynamics
MFP: magnetic force spectroscopy
NA: naphthyridine-azaquinolone
NMR: nuclear magnetic resonance
NOE: nuclear Overhauser effect
NOESY: nuclear Overhauser effect spectroscopy
RMSD: root-mean-square deviation
SCA: spinocerebellar ataxia
UTR: untranslated region

## ACKNOWLEDGMENTS

We thank Xiangming Kong for helping with NMR experiments. The authors acknowledge University of Florida Research Computing for providing computational resources and support that have contributed to the research results reported in this publication (URL: http://www.rc.ufl.edu.).

This study made use of NMR spectrometers at The Scripps Research Institute, Campus Chemical Instrument Center NMR facility at Ohio State University, Minnesota NMR Center, and the National Magnetic Resonance Facility at Madison (NMRFAM). Purchase of the 600 MHz NMR spectrometer at The Scripps Research Institute was supported in part by the National Institutes of Health grant S10OD021550. NMR instrumentation, helium recovery equipment, and computers at NMRFAM were purchased with funds from the University of Wisconsin-Madison, the NIH P41GM136463, R24 GM141526, P41 GM103399, S10 RR023438, S10 RR025062, and S10 RR029220.

