## Supplementary File 1 for "NMR structures and magnetic force spectroscopy studies of small molecules binding to models of an RNA CAG repeat expansion"

### Contents Page


**Supplementary Figures** 4

Figure S1 H6/H8-H1′ region of a 2D $^{1}$H NOESY spectrum of unbound r(CAG). 4

Figure S2 Imino proton region of a 2D $^{1}$H NOESY spectrum of unbound r(CAG). 5

Figure S3 Structure of the unbound r(CAG) motif. 6

Figure S4 1D $^{1}$H and WaterLOGSY NMR spectra of **1**. 7

Figure S5 1D $^{1}$H and WaterLOGSY NMR spectra of **2**. 8

Figure S6 1D $^{1}$H and WaterLOGSY NMR spectra of **3**. 9

Figure S7 1D $^{1}$H imino region titration with **1**. 10

Figure S8 1D $^{1}$H imino region titration with **2**. 11

Figure S9 1D $^{1}$H imino region titration with **3**. 12

Figure S10 Imino proton region of a 2D $^{1}$H NOESY spectrum of the r(CAG)-**1** complex. 13

Figure S11 Imino proton region of a 2D $^{1}$H NOESY spectrum of the r(CAG)-**2** complex. 14

Figure S12 Imino proton region of a 2D $^{1}$H NOESY spectrum of the r(CAG)-**3** complex. 15

Figure S13 Stabilizing interactions of the compounds extracted for the lowest energy NMR structures. 16

Figure S14 Potential of mean force (PMF) with respect to PC1 and PC2 of the RNA complex with **1**, **2** and **3**. 17

Figure S15 Comparison between $r(CAG)_{21}$ and $r(CUG)_{21}$ RNA behavior by magnetic force microscopy. 18

Figure S16 The effect of different compounds binding to the $r(CAG)_{21}$ RNA in ramp and force step experiments. 19

Figure S17 The effect of compounds **1**, **2** and **3** on MBNL1 protein binding. 20

**Supplementary Tables** 21

Table S1 $^{1}$H NMR chemical shifts of the unbound r(CAG) duplex. 21

Table S2 $^{1}$H NMR chemical shifts of r(CAG)-**1** complex. 22

| | | |
|---|---|---|
| Table S3 | $^{1}$H NMR chemical shifts of r(CAG)-**2** complex. | 23 |
| Table S4 | $^{1}$H NMR chemical shifts of r(CAG)-**3** complex. | 24 |
| Table S5 | NOE restraints used for modeling of the unbound r(CAG) duplex. | 25 |
| Table S6 | Dihedral restraints used for modeling of the unbound r(CAG) duplex. | 32 |
| Table S7 | NOE restraints used for modeling of the r(CAG)-**1** complex. | 36 |
| Table S8 | NOE restraints used for modeling of the r(CAG)-**2** complex. | 41 |
| Table S9 | NOE restraints used for modeling of the r(CAG)-**3** complex. | 46 |
| Table S10 | Summary of select helical and base pair parameters for apo- and ligand-bound RNA constructs. | 52 |

**References** 53

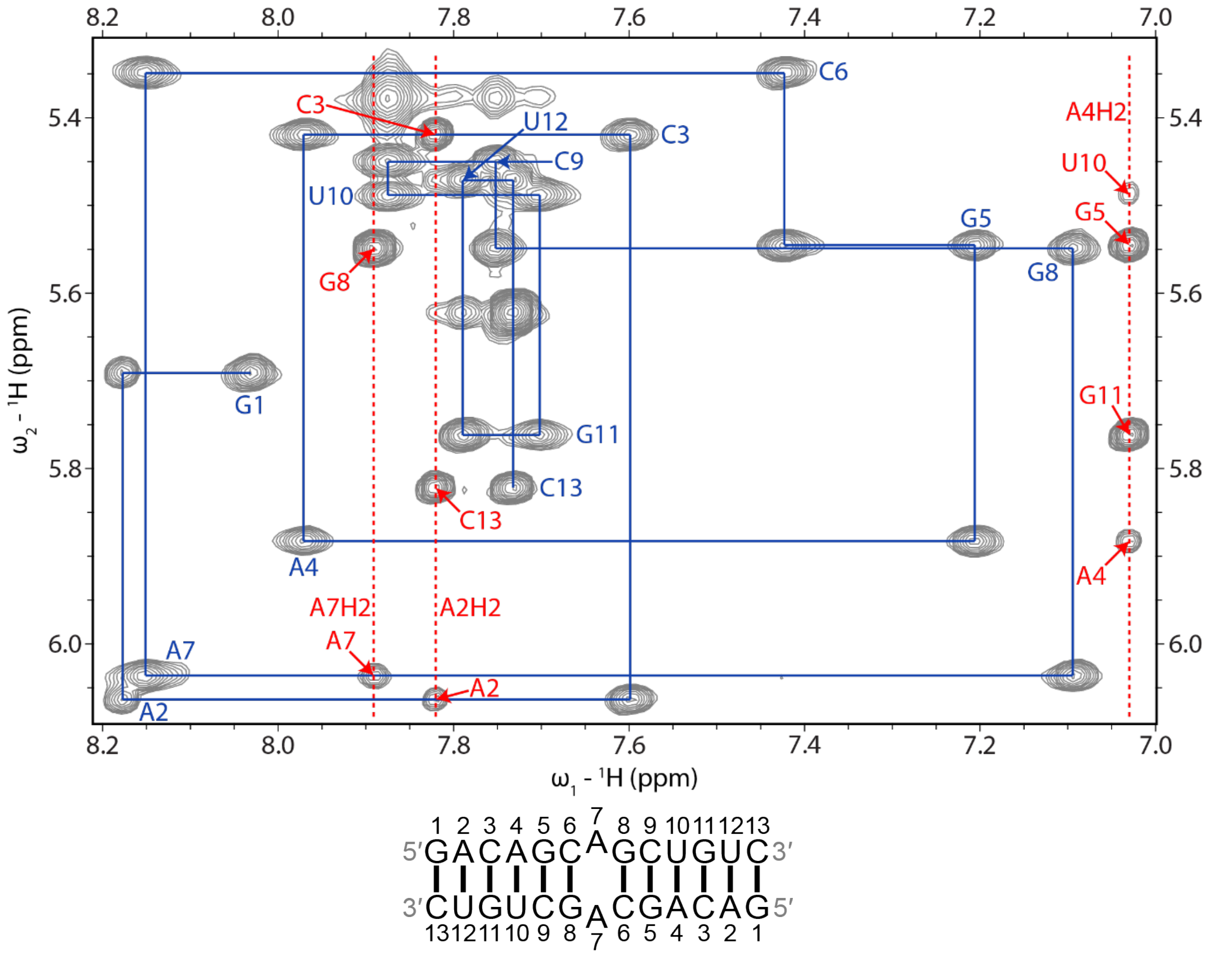


**Figure S1: H6/H8-H1′ region of a 2D $^1$H NOESY spectrum of unbound r(CAG).** A sequential H6/H8-H1′ walk is shown with blue lines. Blue labels correspond to intraresidue H6/H8-H1′ NOEs. Adenine H2 resonances are labeled red with dashed lines. The spectrum was acquired at 25 °C with 400 ms mixing time and 0.7 mM of RNA.

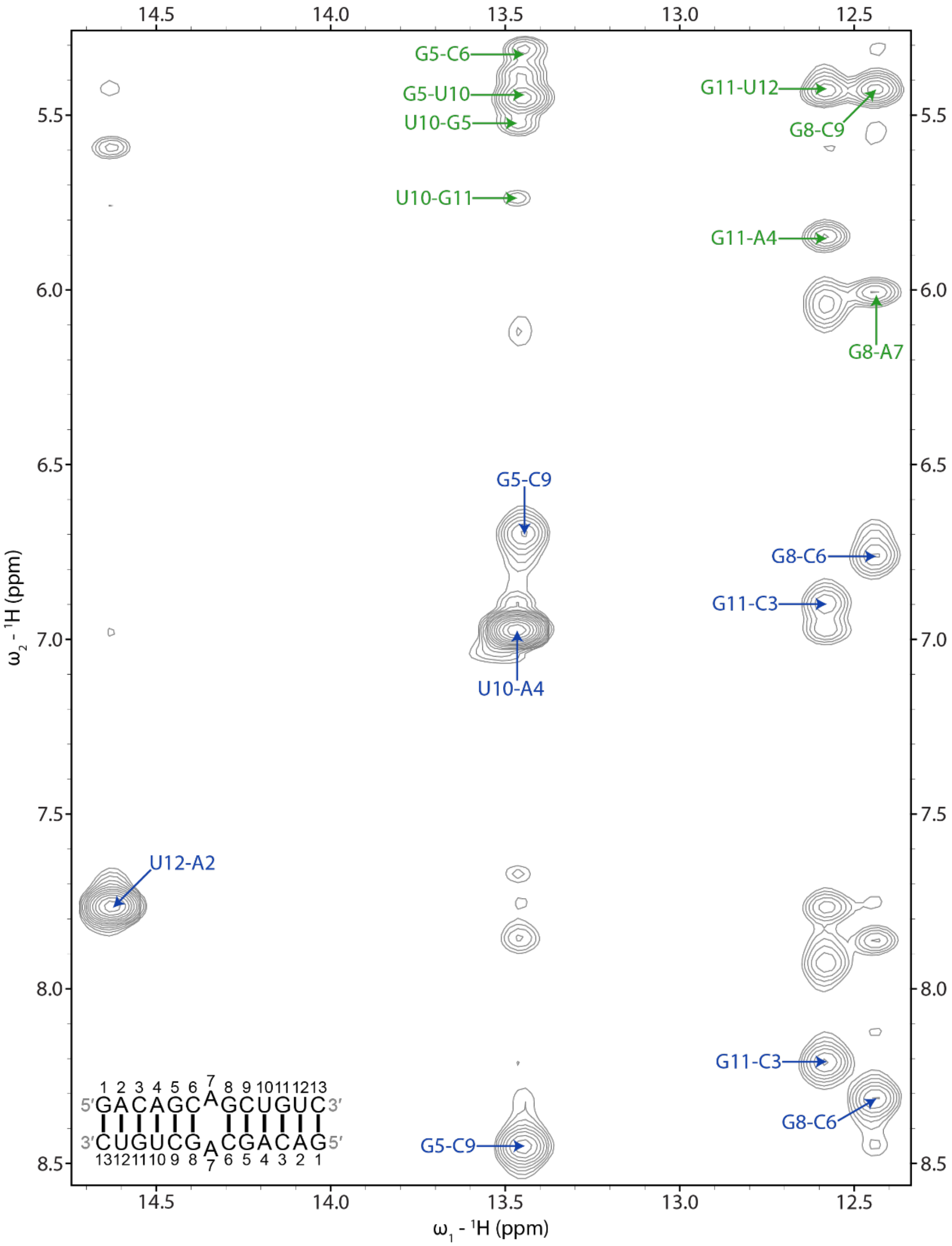


**Figure S2: Imino proton region of a 2D $^{1}$H NOESY spectrum of unbound r(CAG).** Blue labels correspond to UH3-AH2 and GH1 to C amino NOEs within base pairs. Green labels correspond to NOEs between UH3 or GH1 and the H1′ of a 3′ adjacent or cross-strand residue. In each label, the first residue corresponds to UH3 or GH1 and the second label corresponds to AH2, a C amino proton, or H1′. The spectrum was acquired at 5 °C with 125 ms mixing time and 0.7 mM of RNA.

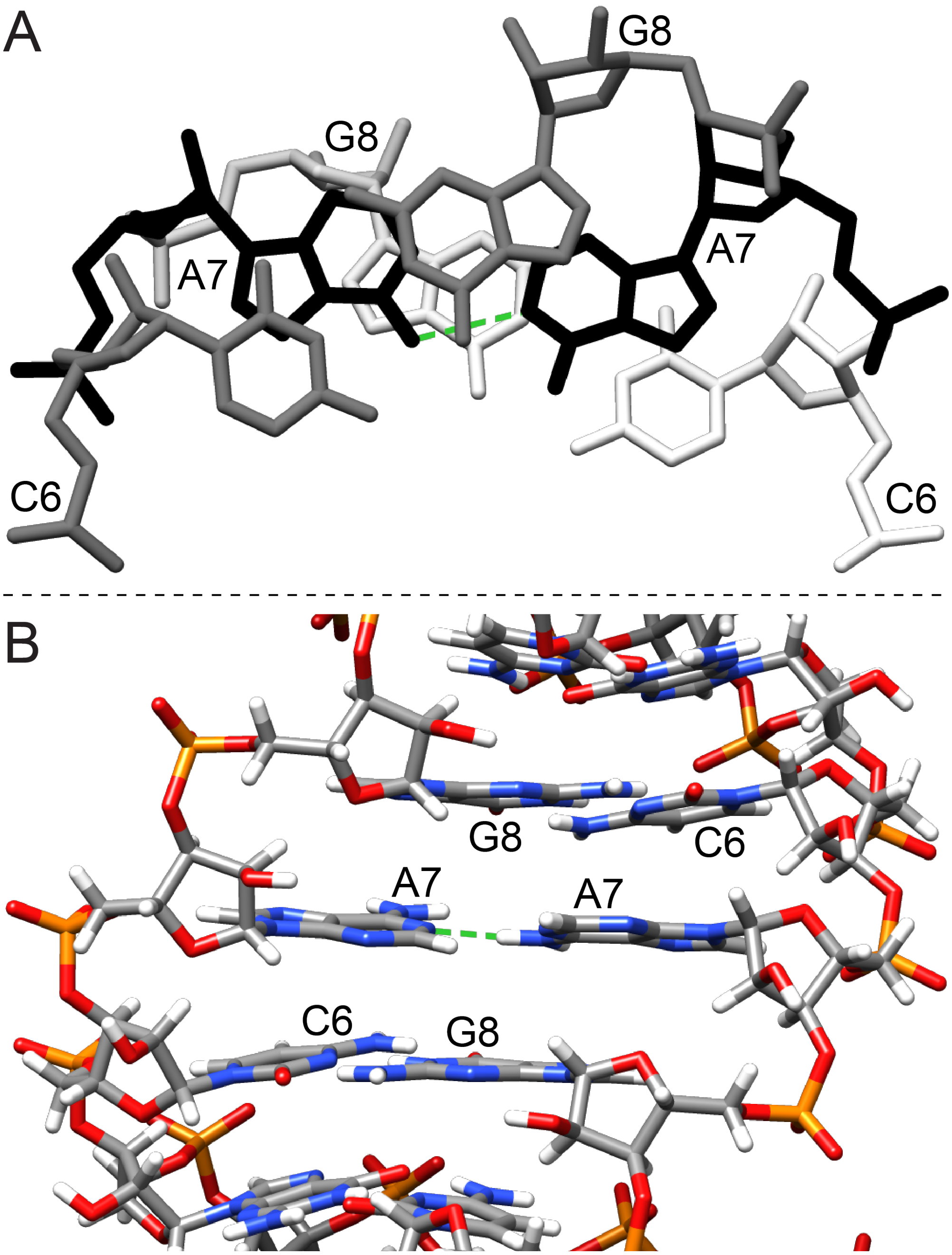


**Figure S3: Structure of the unbound r(CAG) motif.** (A) Major groove view showing overtwisting and undertwisting of the helix. The AA pair is colored black. (B) Minor groove view showing the stacking of the AA mismatch in the helix. The AA pair in each view contains one interbase hydrogen bond, as indicated with a green dashed line.

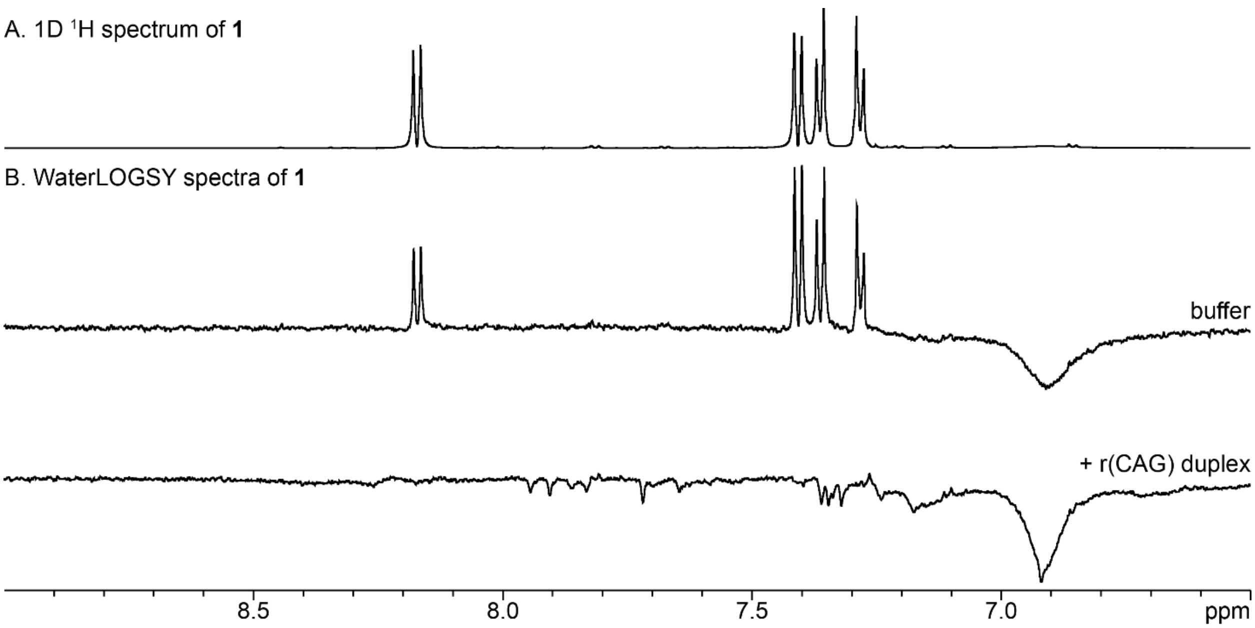


**Figure S4: 1D $^{1}$H and WaterLOGSY NMR spectra of 1.** (A) 1D $^{1}$H NMR spectrum of 300 μM of **1** in NMR Buffer. (B) WaterLOGSY spectra of 300 μM of **1** in NMR Buffer with or without 10 μM of r(CAG). Spectra were acquired at 25 °C.

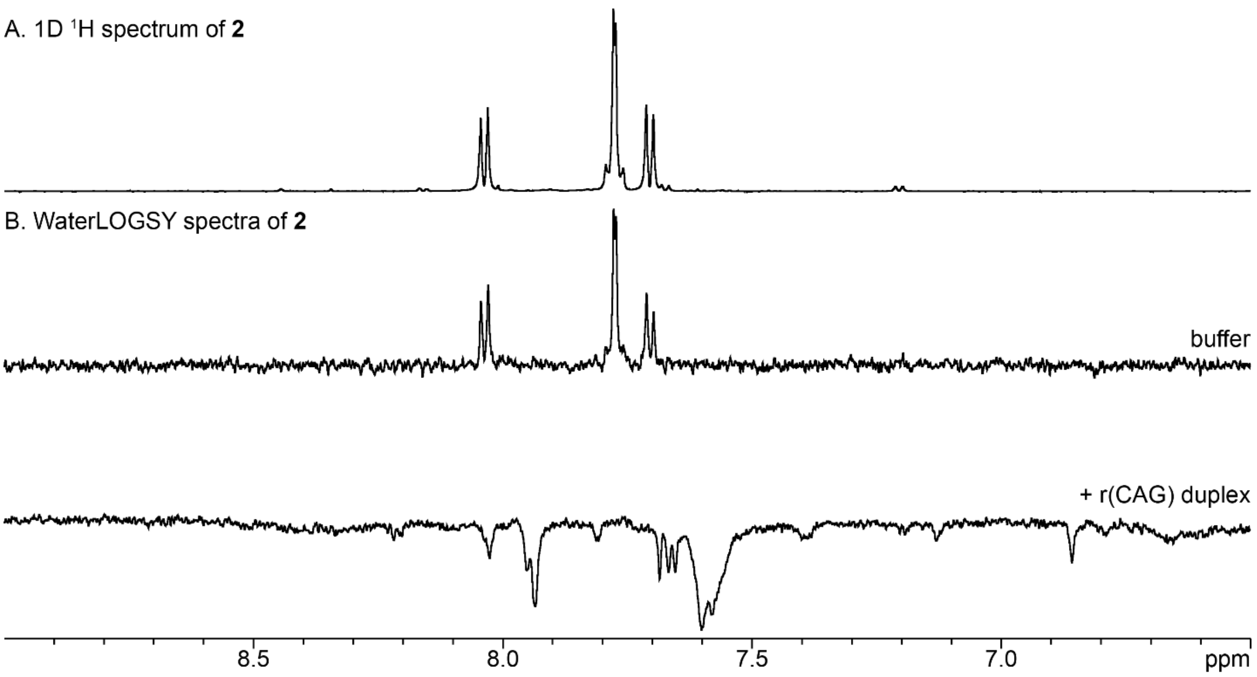


**Figure S5: 1D $^1$H and WaterLOGSY NMR spectra of 2.** (A) 1D $^1$H NMR spectrum of 300 μM of **2** in NMR Buffer. (B) WaterLOGSY spectra of 300 μM of **2** in NMR Buffer with or without 10 μM of r(CAG). Spectra were acquired at 25 °C.

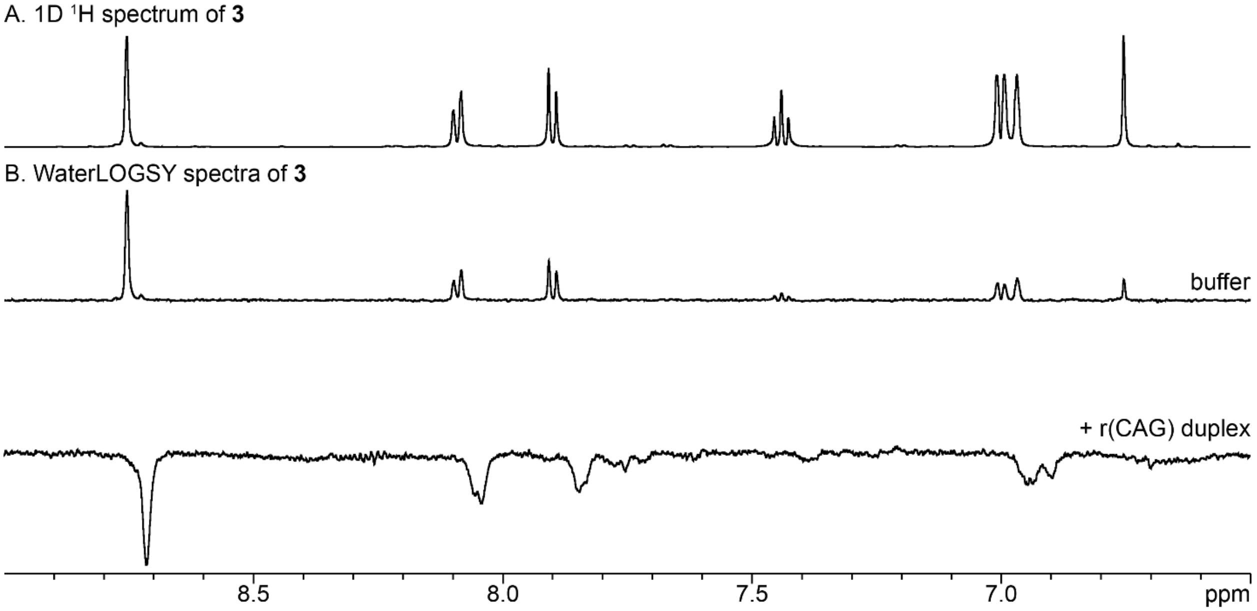


**Figure S6: 1D $^1$H and WaterLOGSY NMR spectra of 3.** (A) 1D $^1$H NMR spectrum of 300 μM of **3** in NMR Buffer. (B) WaterLOGSY spectra of 300 μM of **3** in NMR Buffer with or without 10 μM of r(CAG). Spectra were acquired at 25 °C.

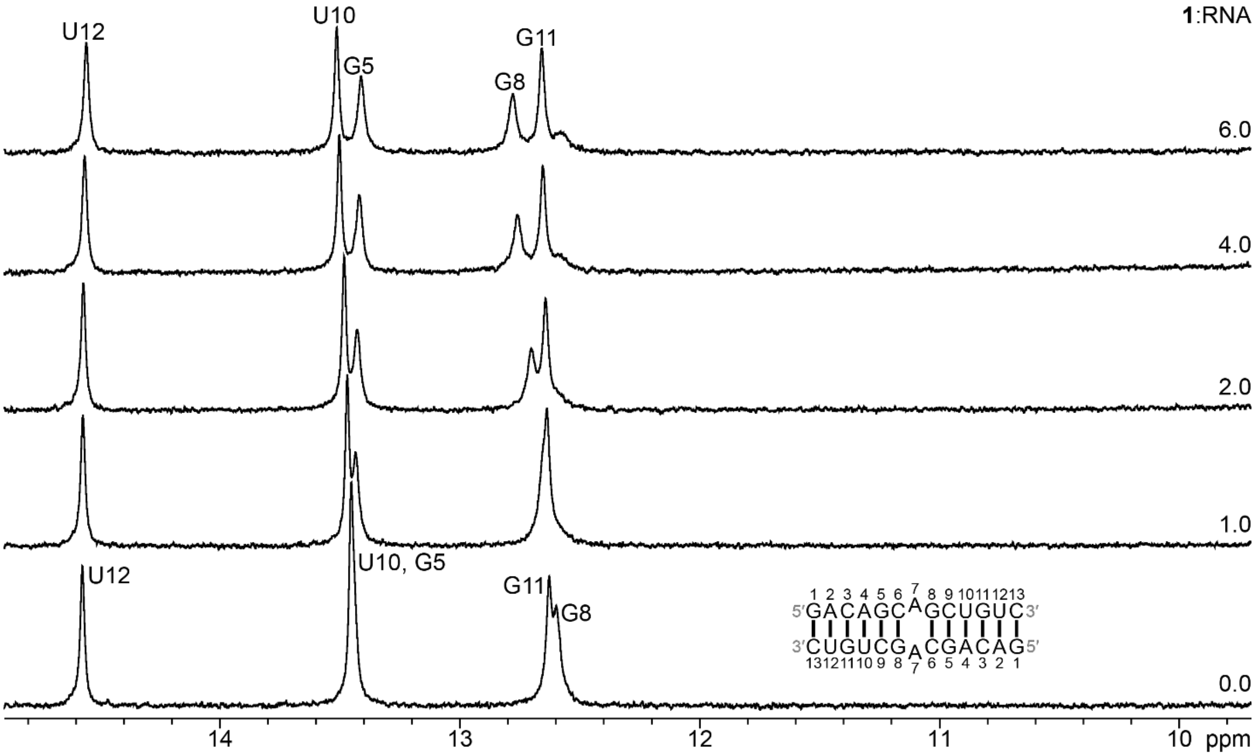


**Figure S7: 1D $^1$H spectra of the imino region of the r(CAG) duplex upon titration with 1.** Spectra were acquired in NMR Buffer at 5 °C with 50 μM of RNA and 0 to 300 μM of **1**. The molar ratio of **1** : RNA is indicated on the right.

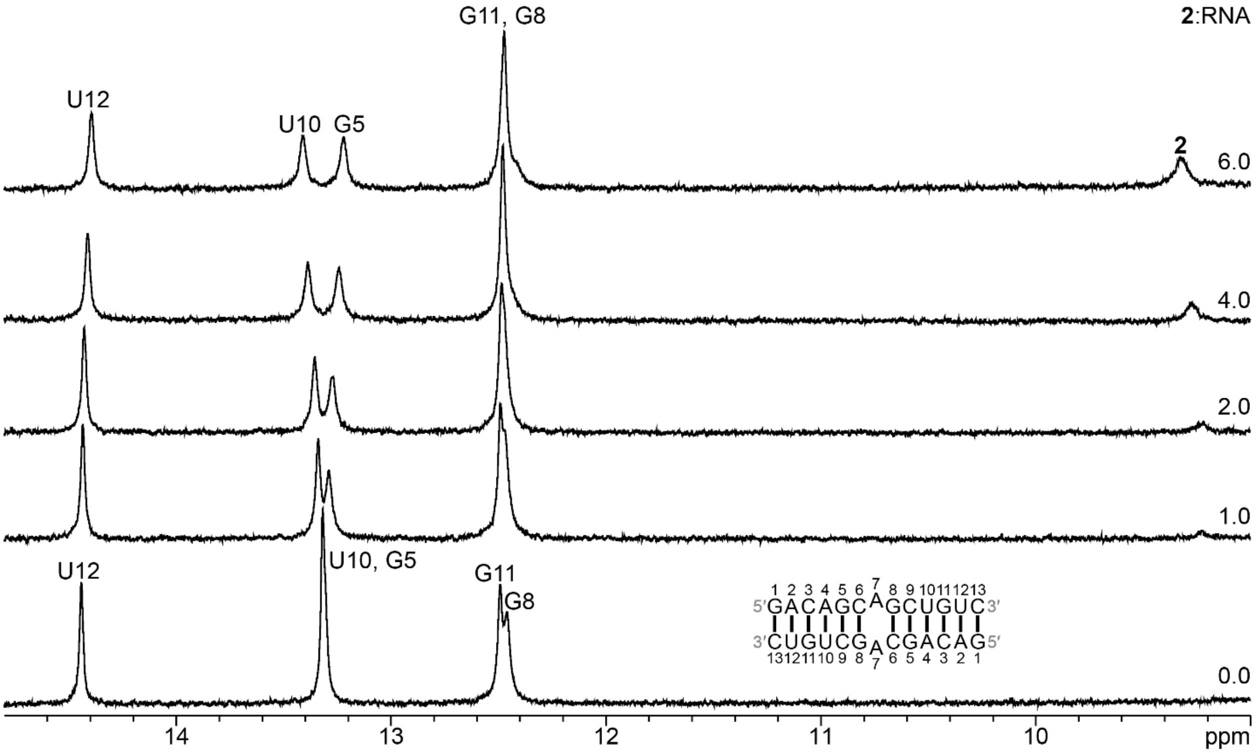


**Figure S8: 1D ¹H spectra of the imino region of the r(CAG) duplex upon titration with 2.** Spectra were acquired in NMR Buffer at 5 °C with 50 μM of RNA and 0 to 300 μM of **2**. The molar ratio of **2** : RNA is indicated on the right.

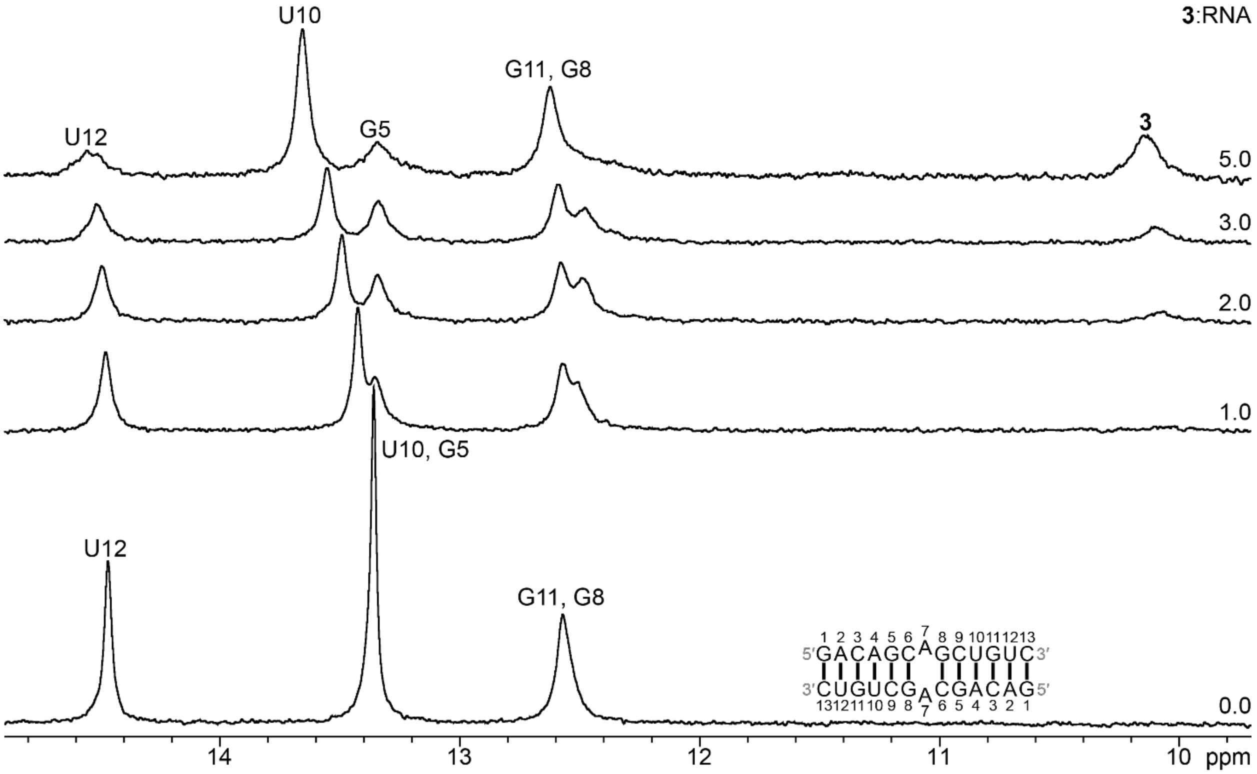


**Figure S9: 1D $^{1}$H spectra of the imino region of the r(CAG) duplex upon titration with 3.** Spectra were acquired in NMR Buffer at 5 °C with 50 μM of RNA and 0 to 300 μM of **3**. The molar ratio of **3** : RNA is indicated on the right.

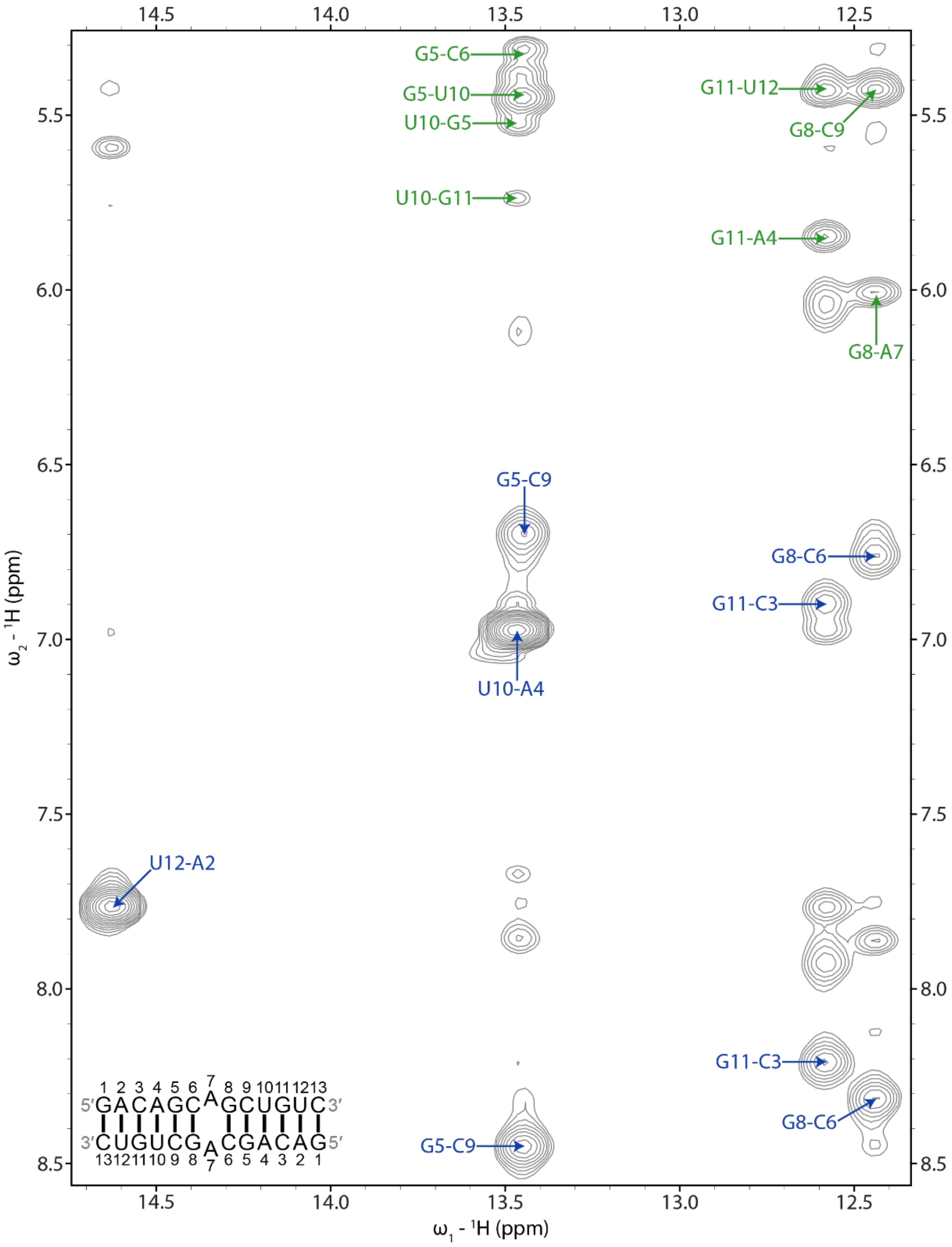


**Figure S10: Imino proton region of a 2D $^1$H NOESY spectrum of the r(CAG)-1 complex.** Blue labels correspond to UH3-AH2 and GH1 to C amino NOEs within base pairs. Green labels correspond to NOEs between UH3 or GH1 and the H1′ of a 3′ adjacent or cross-strand residue. In each label, the first residue corresponds to UH3 or GH1 and the second label corresponds to AH2, a C amino proton, or H1′ of a nearby residue. The spectrum was acquired in NMR Buffer at 5 °C with 125 ms mixing time and 0.7 mM of r(CAG) and 0.7 mM of **1**.

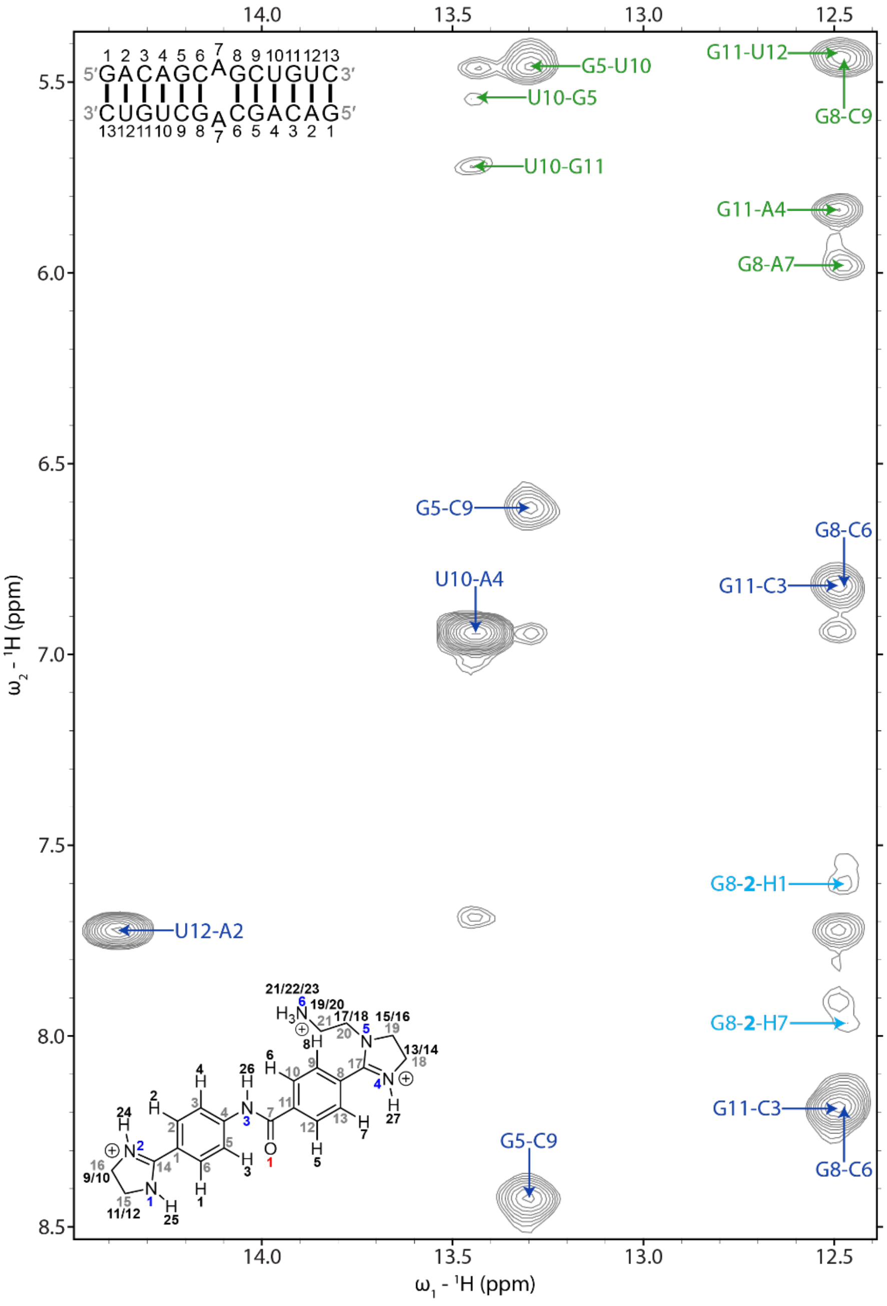


**Figure S11: Imino proton region of a 2D $^{1}$H NOESY spectrum of the r(CAG)-2 complex.** Blue labels correspond to UH3-AH2 and GH1 to C amino NOEs within base pairs. Green labels correspond to NOEs between UH3 or GH1 and the H1′ of a 3′ adjacent or cross-strand residue. Light blue labels correspond to NOEs between G8H1 and **2**. In each label, the first residue corresponds to UH3 or GH1 and the second label corresponds to AH2, a C amino proton, or H1′ of a nearby residue. In the chemical structure of **2**, black numbers correspond to hydrogens, gray numbers correspond to carbons, blue numbers correspond to nitrogen, and red numbers correspond to oxygen. The spectrum was acquired in NMR Buffer at 15 °C with 125 ms mixing time and 0.3 mM of r(CAG) and 0.6 mM of **2**.

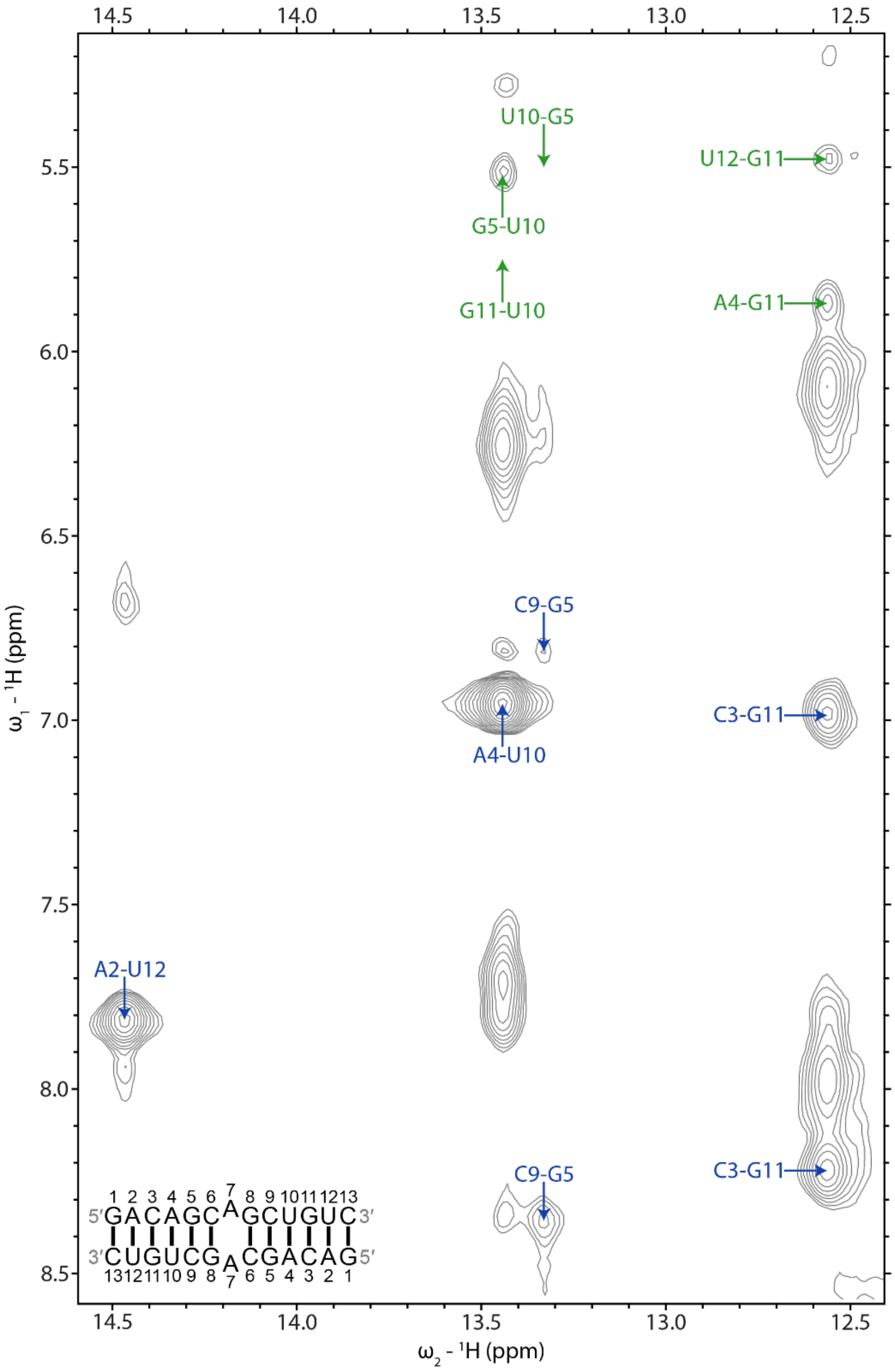


**Figure S12: Imino proton region of a 2D $^1$H NOESY spectrum of the r(CAG)-3 complex.** Blue labels correspond to UH3-AH2 and GH1 to C amino NOEs within base pairs. Green labels correspond to NOEs between UH3 or GH1 and the H1′ of a 3′ adjacent or cross-strand residue. In each label, the first residue corresponds to UH3 or GH1 and the second residue corresponds to AH2, a C amino proton, or H1′ of a nearby residue. The spectrum was acquired in NMR Buffer at 5 °C with 125 ms mixing time and 0.4 mM of r(CAG) and 0.6 mM of **3**.

A B C

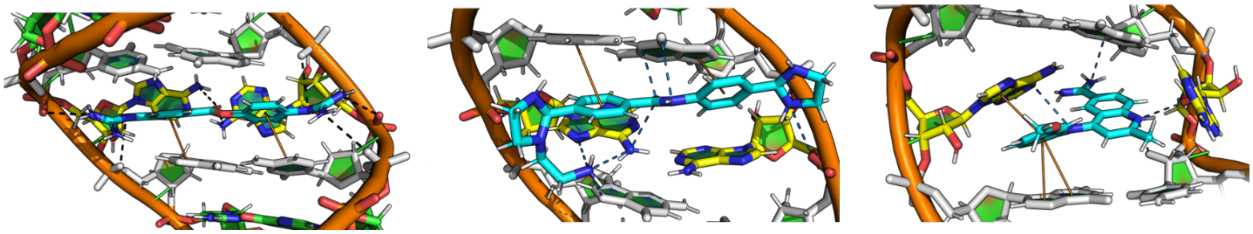

**Figure S13. Stabilizing interactions between a r(CAG) repeat duplex model and compounds, extracted for the lowest energy NMR-restrained structures.** (A) A combination of hydrogen bonds and stacking interactions with the neighboring base pairs stabilizes the bound pose of **1**. (B) **2** is stabilized by multiple hydrogen bonds to the A nucleotide of the A/A loop and the neighboring base pairs as well as hydrogen bonds to the backbone. It also forms stacking interactions with the neighboring base pairs. (D) **3** makes stacking interactions with A nucleotide of the A/A loop and the neighboring base pairs as well as hydrogen bonds. Hydrogen bonds are indicated by the dashed lines.

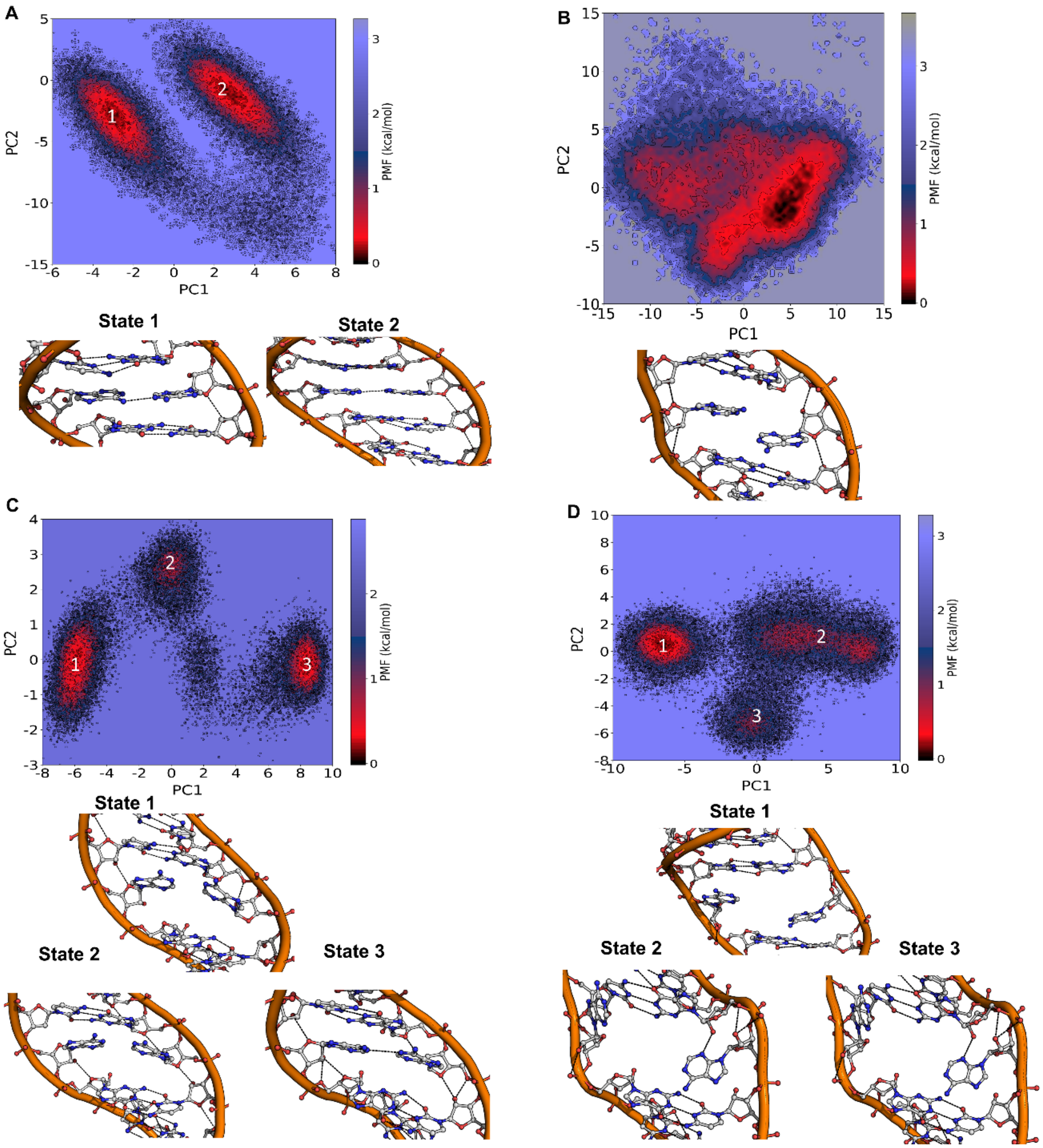


**Figure S14. Potential of mean force (PMF) with respect to PC1 and PC2 of the RNA complex with 1, 2, and 3.** (A) Analysis of the free energy landscape with respect to PC1 and PC2 of the A/A loop of the Apo-form shows two minima (left) corresponding to two observed clusters (bottom). (B) RNA bound to **1** just shows one minimum (bottom) where there are no hydrogen bonds between the A/A nucleotides (right). (C) **3** induces drastic conformational changes as observed by three distinct minima along the free energy landscape (bottom) corresponding to three different clusters. (D) Although **3** is the only compounds which pushes the A nucleotides out of the helical axis the conformational changes are not as strong as observed with **2**, creating one distinct minimum and two low populated one corresponding to observed clusters (bottom).

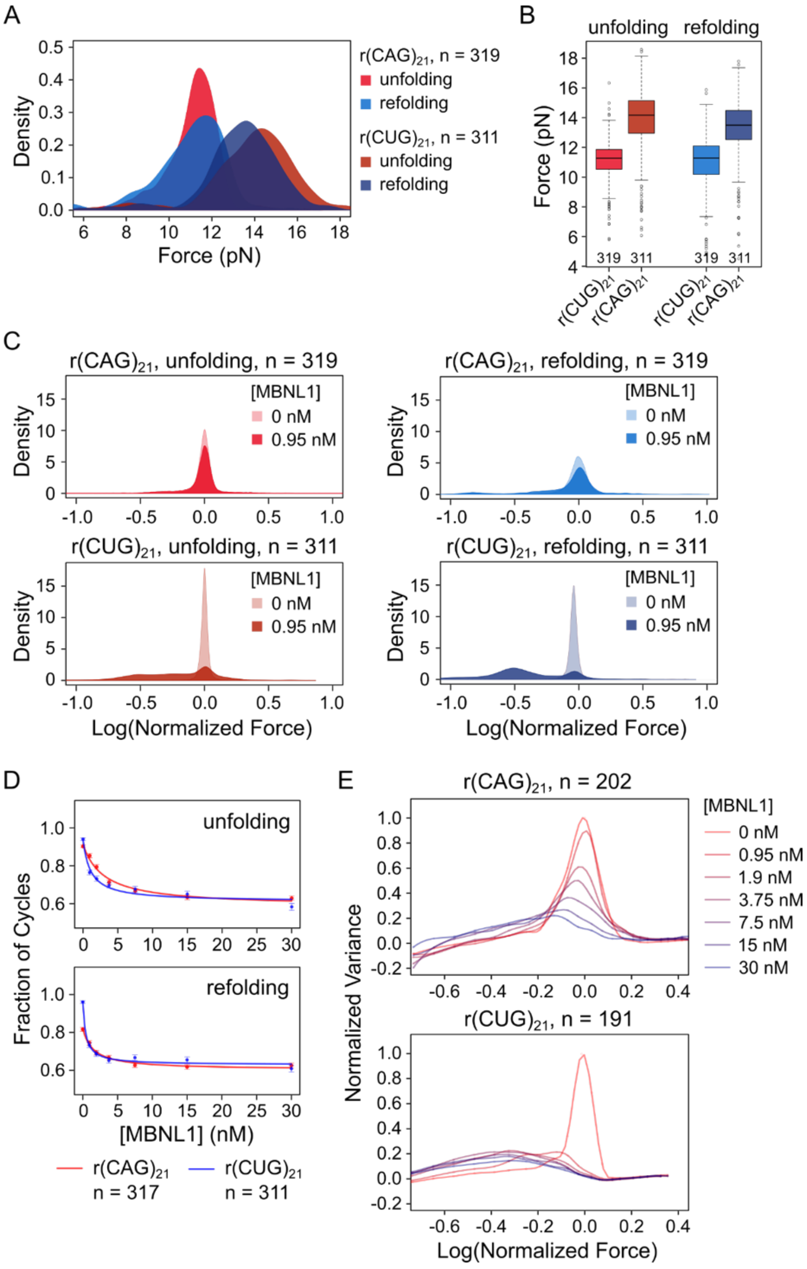


**Figure S15. Comparison between r(CAG)$_{21}$ and r(CUG)$_{21}$ RNA behavior by magnetic force microscopy**. (A) the raw unfolding and refolding force distribution for ramp experiments comparing r(CAG)$_{21}$ and r(CUG)$_{21}$. The median unfolding force of each molecule is used to plot the force distribution, and the number of molecules used is indicated as 'n'. (B) Boxplot comparing the unfolding and refolding force distribution, using the median force of each molecule. (C) Unfolding and refolding force distribution, comparing r(CAG)$_{21}$ and r(CUG)$_{21}$ in the presence of 0.95 nM of MBNL1 protein. (D) Unfolding and refolding probability comparing the same concentration of MBNL1 protein binding to r(CUG)$_{21}$ or r(CUG)$_{21}$. (E) Graph showing the normalized variance height against normalized force in stepped force experiments, testing different MBNL1 protein concentrations binding to r(CAG)$_{21}$ (left) or r(CUG)$_{21}$ (right) molecules.

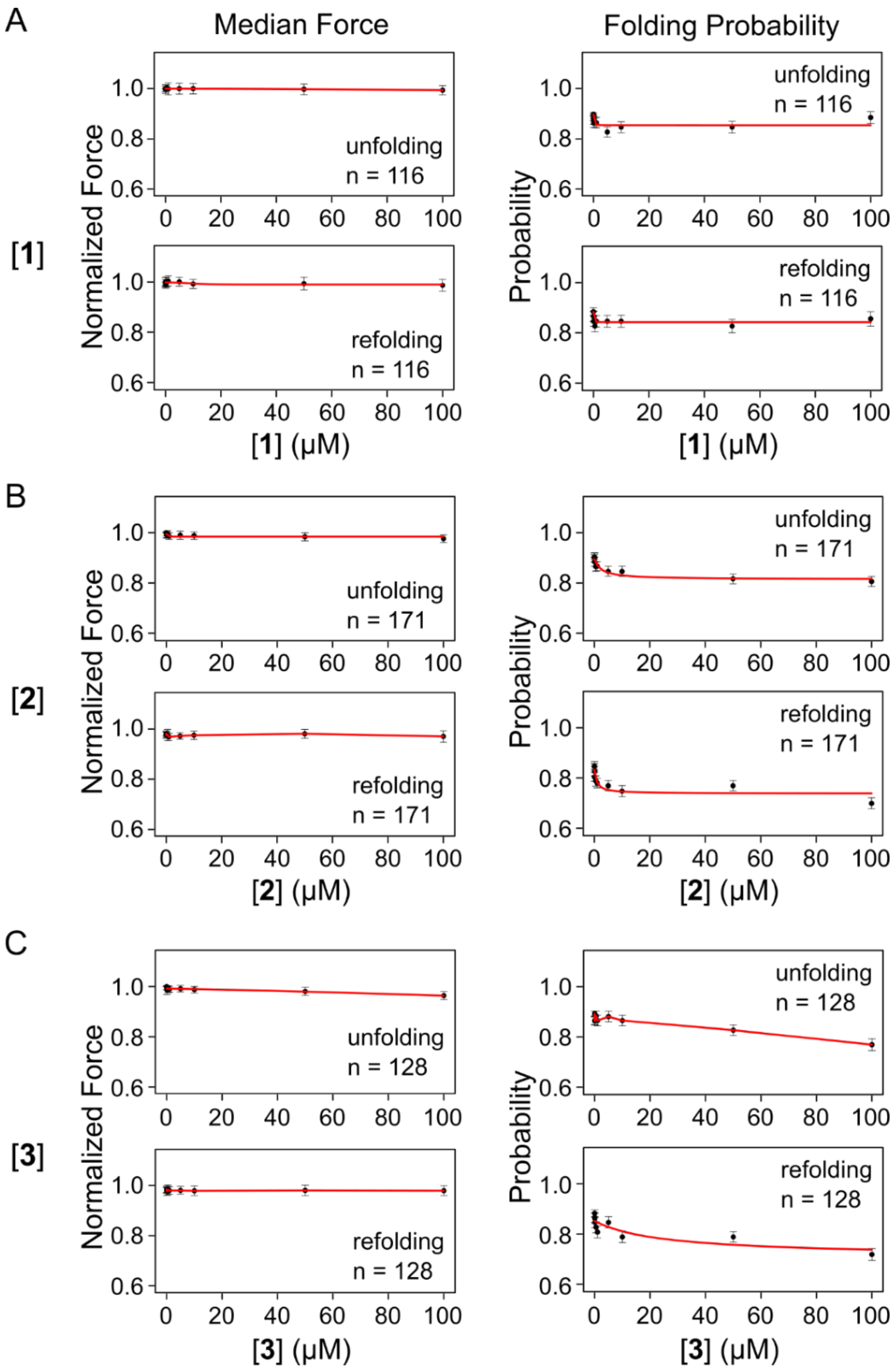


**Figure S16. The effect of different compounds binding to the r(CAG)$_{21}$ RNA in force ramp and stepped force experiments**. The normalized median unfolding and refolding (left) forces and probabilities (right) obtained in force ramp experiments were plotted with increasing concentrations of **1** (A), **2** (B), and **3** (C). n is the number of molecules analyzed.

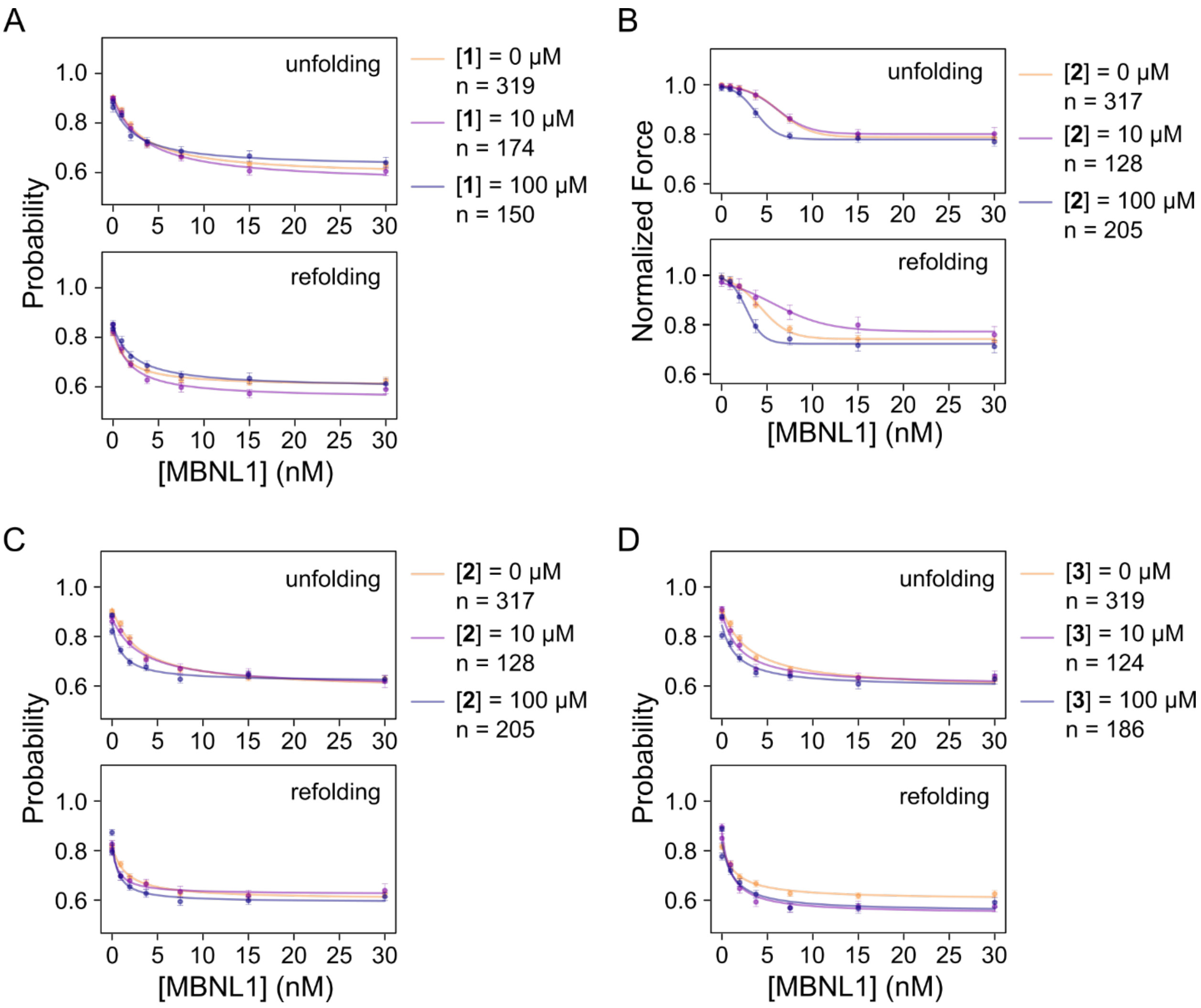


**Figure S17. The effects of 1, 2, and 3 on MBNL1 protein binding, as determined from single molecule studies**. (A) The unfolding and refolding probabilities with increasing concentrations of MBNL1 protein in the presence of 0, 10, and 100 μM of **1** in force ramp experiments. The graph shows the median of all molecules ± SEM, and the fitted curves. (B) The median unfolding and refolding force as a function of increasing MBNL1 protein concentration in the presence of 0, 10, and 100 μM of **2**. (C) The unfolding and refolding probability effect of MBNL1 protein with 0, 10, and 100 μM of **2**. (D) The unfolding and refolding probability effect of MBNL1 protein with 0, 10, and 100 μM of **3**.

**Table S1**: $^{1}$H NMR chemical shifts of the *apo* r(CAG) duplex.

| Residue | H1′ | H2′ | H3′ | H4′ | H2/H5 | H6/H8 | H1/H3 | H21/41/61, H22/42/62 amino |
|---|---|---|---|---|---|---|---|---|
| G1 | 5.691 | 4.814 | 4.642 | 4.38 | n/a | 8.031 | - | - |
| A2 | 6.063 | 4.512 | 4.777 | 4.54 | 7.82 | 8.177 | n/a | - |
| C3 | 5.419 | 4.317 | 4.512 | 4.406 | 5.198 | 7.599 | n/a | 8.210, 6.900 |
| A4 | 5.883 | 4.622 | 4.71 | 4.478 | 7.03 | 7.971 | n/a | - |
| G5 | 5.545 | 4.341 | 4.388 | - | n/a | 7.206 | 13.44 | - |
| C6 | 5.349 | 4.284 | 4.496 | - | 5.1 | 7.423 | n/a | 8.313, 6.763 |
| A7 | 6.036 | 4.62 | 4.654 | 4.495 | 7.891 | 8.151 | n/a | - |
| G8 | 5.549 | 4.466 | 4.289 | - | n/a | 7.094 | 12.44 | - |
| C9 | 5.45 | 4.289 | - | - | 5.041 | 7.752 | n/a | 8.450, 6.701 |
| U10 | 5.488 | 4.623 | 4.585 | 4.423 | 5.378 | 7.875 | 13.47 | n/a |
| G11 | 5.762 | 4.408 | 4.552 | 4.471 | n/a | 7.702 | 12.58 | - |
| U12 | 5.471 | 4.22 | 4.452 | 4.372 | 5.088 | 7.789 | 14.62 | n/a |
| C13 | 5.822 | 3.962 | 4.173 | - | 5.622 | 7.732 | n/a | 8.348, 6.985 |

Chemical shifts are in ppm units. Non-exchangeable protons were assigned at 25 °C. Exchangeable protons were assigned at 5 °C.

Table S2: $^{1}$H NMR chemical shifts of r(CAG)-**1** complex.

$^{1}$H NMR chemical shifts of RNA in the r(CAG)-**1** complex.

| Residue | H1′ | H2′ | H3′ | H4′ | H2/H5 | H6/H8 | H1/H3 | H21/41/61, H22/42/62 amino |
|---|---|---|---|---|---|---|---|---|
| G1 | 5.641 | 4.786 | 4.625 | 4.363 | n/a | 8.020 | - | - |
| A2 | 6.045 | 4.499 | - | 4.525 | 7.798 | 8.158 | n/a | - |
| C3 | 5.405 | 4.306 | 4.499 | 4.396 | 5.184 | 7.592 | n/a | 8.184, 6.844 |
| A4 | 5.871 | 4.582 | 4.704 | 4.467 | 7.025 | 7.96 | n/a | - |
| G5 | 5.552 | 4.332 | 4.406 | - | n/a | 7.221 | 13.41 | - |
| C6 | 5.371 | 4.304 | 4.441 | - | 5.077 | 7.394 | n/a | 8.176, 6.689 |
| A7 | 6.031 | 4.58 | 4.632 | - | 7.944 | 8.072 | n/a | - |
| G8 | 5.464 | 4.465 | 4.332 | - | n/a | 7.212 | 12.49 | - |
| C9 | 5.471 | 4.314 | - | - | 5.052 | 7.745 | n/a | 8.427, 6.65 |
| U10 | 5.493 | 4.621 | 4.582 | 4.425 | 5.376 | 7.883 | 13.46 | n/a |
| G11 | 5.756 | 4.402 | 4.550 | 4.467 | n/a | 7.714 | 12.56 | - |
| U12 | 5.458 | 4.205 | 4.449 | 4.364 | 5.083 | 7.786 | 14.57 | n/a |
| C13 | 5.801 | 3.929 | 4.132 | - | 5.587 | 7.706 | n/a | - |

$^{1}$H NMR chemical shifts of **1** in the r(CAG)-**1** complex.

| Hydrogen | Chemical shift |
|---|---|
| H1/H2 | 7.163 |
| H3/H4 | 7.267 |
| H5/H6 | 7.886 |
| H7/H8 | 7.202 |

Chemical shifts are in ppm units. Non-exchangeable protons were assigned at 25 °C. Exchangeable protons were assigned at 5 °C.

Table S3: $^1$H NMR chemical shifts of r(CAG)-**2** complex.

$^1$H NMR chemical shifts of RNA in the r(CAG)-**2** complex.

| Residue | H1′ | H2′ | H3′ | H4′ | H2/H5 | H6/H8 | H1/H3 | H21/41/61, H22/42/62 amino |
|---|---|---|---|---|---|---|---|---|
| G1 | 5.599 | 4.724 | 4.616 | 4.341 | n/a | 8.000 | - | - |
| A2 | 6.02 | 4.473 | - | 4.515 | 7.755 | 8.153 | n/a | - |
| C3 | 5.382 | 4.282 | - | 4.384 | 5.168 | 7.573 | n/a | 8.190, 6.820 |
| A4 | 5.852 | 4.565 | 4.689 | 4.458 | 6.994 | 7.938 | n/a | - |
| G5 | 5.54 | 4.275 | 4.392 | - | n/a | 7.202 | 13.3 | - |
| C6 | 5.391 | 4.248 | - | - | 5.042 | 7.371 | n/a | 8.184, 6.824 |
| A7 | 5.998 | 4.599 | 4.657 | - | 7.857 | 8.129 | n/a | - |
| G8 | 5.454 | 4.461 | 4.346 | - | n/a | 7.244 | 12.47 | - |
| C9 | 5.455 | 4.294 | - | - | 5.049 | 7.732 | n/a | 8.425, 6.616 |
| U10 | 5.477 | 4.608 | 4.576 | 4.417 | 5.346 | 7.878 | 13.44 | n/a |
| G11 | 5.741 | 4.376 | 4.543 | 4.461 | n/a | 7.693 | 12.49 | - |
| U12 | 5.441 | 4.184 | 4.422 | 4.362 | 5.03 | 7.761 | 14.38 | n/a |
| C13 | 5.772 | 3.944 | 4.044 | 4.13 | 5.525 | 7.657 | n/a | - |

$^1$H NMR chemical shifts of **2** in the r(CAG)-**2** complex.

| Hydrogen | Chemical shift |
|---|---|
| H1/H2 | 7.507 |
| H3/H4 | 7.594 |
| H5/H6 | 7.961 |
| H7/H8 | 7.711 |

Chemical shifts are in ppm units. Non-exchangeable protons were assigned at 25 °C. Exchangeable protons were assigned at 15 °C.

**Table S4**: $^1$H NMR chemical shifts of r(CAG)-**3** complex.

$^1$H NMR chemical shifts of RNA in the r(CAG)-**3** complex.

| Residue | H1′ | H2′ | H3′ | H4′ | H2/H5 | H6/H8 | H1/H3 | H21/41/61, H22/42/62 amino |
|---|---|---|---|---|---|---|---|---|
| G1 | 5.649 | 4.773 | 4.637 | - | n/a | 7.974 | - | - |
| A2 | 6.052 | 4.497 | 4.783 | 4.545 | 7.827 | 8.182 | n/a | - |
| C3 | 5.441 | 4.333 | 4.538 | 4.415 | 5.186 | 7.641 | n/a | 8.221, 6.986 |
| A4 | 5.896 | 4.601 | - | 4.488 | 7.055 | 7.968 | n/a | - |
| G5 | 5.563 | 4.316 | 4.418 | 4.495 | n/a | 7.218 | 13.33 | - |
| C6 | 5.429 | 4.24 | - | - | 5.092 | 7.409 | n/a | - |
| A7 | 6.004 | 4.626 | - | 4.509 | 7.837 | 8.182 | n/a | - |
| G8 | 5.496 | 4.501 | 4.401 | - | n/a | 7.325 | - | - |
| C9 | 5.493 | 4.322 | 4.485 | - | 5.081 | 7.781 | n/a | 8.358, 6.813 |
| U10 | 5.518 | - | 4.623 | - | 5.336 | 7.926 | 13.44 | n/a |
| G11 | 5.783 | 4.428 | 4.581 | 4.483 | n/a | 7.751 | 12.56 | - |
| U12 | 5.484 | 4.21 | 4.473 | 4.389 | 5.121 | 7.798 | 14.47 | n/a |
| C13 | 5.842 | 3.943 | 4.178 | - | 5.641 | 7.76 | n/a | 8.429, 7.298 |

$^1$H NMR chemical shifts of **3** in the r(CAG)-**3** complex.

| Hydrogen | Chemical shift |
|---|---|
| H1 | 8.708 |
| H2 | 8.034 |
| H3 | 7.773 |
| H4 | 6.692 |
| H5 | 6.878 |
| H6 | 6.917 |
| H7 | 7.378 |
| H8 | 6.927 |
| H15/16/17 | 2.473 |
| H18/19/20 | 3.773 |

Chemical shifts are in ppm units. Non-exchangeable protons were assigned at 35 °C. Exchangeable protons were assigned at 10 °C.

**Table S5:** NOE restraints used for modeling of the *apo* r(CAG) duplex.

| | | | | | | | |
|---|---|---|---|---|---|---|---|
| 1 | G | H1 | 26 | C | N3 | 1.8 | 2.4 |
| 1 | G | H1' | 1 | G | H3' | 3.21 | 5.13 |
| 1 | G | H1' | 1 | G | H4' | 2.98 | 4.75 |
| 1 | G | H22 | 26 | C | O2 | 1.8 | 2.4 |
| 1 | G | H8 | 1 | G | H1' | 2.65 | 4.23 |
| 1 | G | H8 | 1 | G | H2' | 2.62 | 4.18 |
| 1 | G | H8 | 1 | G | H3' | 2.31 | 3.69 |
| 1 | G | H8 | 1 | G | H4' | 3.06 | 4.88 |
| 1 | G | O6 | 26 | C | H41 | 1.8 | 2.4 |
| 2 | A | H1' | 1 | G | H2' | 3.44 | 5.48 |
| 2 | A | H1' | 2 | A | H3' | 3.83 | 6.11 |
| 2 | A | H2 | 2 | A | H1' | 3.82 | 6.1 |
| 2 | A | H2 | 3 | C | H1' | 2.74 | 4.37 |
| 2 | A | H2 | 26 | C | H1' | 2.89 | 4.61 |
| 2 | A | H61 | 25 | U | O4 | 1.8 | 2.4 |
| 2 | A | H8 | 1 | G | H2' | 2.25 | 3.59 |
| 2 | A | H8 | 2 | A | H1' | 3.06 | 4.88 |
| 2 | A | H8 | 3 | C | H5 | 4.12 | 6.56 |
| 2 | A | N1 | 25 | U | H3 | 1.8 | 2.4 |
| 3 | C | H1' | 3 | C | H2' | 2.23 | 3.55 |
| 3 | C | H1' | 3 | C | H4' | 2.8 | 4.47 |
| 3 | C | H41 | 24 | G | H1 | 1.21 | 2.81 |
| 3 | C | H41 | 24 | G | O6 | 1.8 | 2.4 |
| 3 | C | H41 | 25 | U | H3 | 1.55 | 3.61 |
| 3 | C | H5 | 2 | A | H2' | 3.38 | 5.39 |
| 3 | C | H6 | 2 | A | H2' | 1.71 | 2.72 |
| 3 | C | H6 | 2 | A | H3' | 3.25 | 5.19 |
| 3 | C | H6 | 3 | C | H1' | 2.63 | 4.2 |
| 3 | C | H6 | 3 | C | H2' | 2.72 | 4.34 |
| 3 | C | N3 | 24 | G | H1 | 1.8 | 2.4 |
| 3 | C | O2 | 24 | G | H22 | 1.8 | 2.4 |
| 4 | A | H1' | 3 | C | H2' | 3.58 | 5.71 |
| 4 | A | H1' | 4 | A | H2' | 2.27 | 3.62 |
| 4 | A | H1' | 4 | A | H3' | 3.84 | 6.12 |
| 4 | A | H1' | 4 | A | H4' | 3.39 | 5.41 |
| 4 | A | H1' | 24 | G | H1 | 3 | 6 |
| 4 | A | H2 | 4 | A | H1' | 4.08 | 6.51 |
| 4 | A | H2 | 5 | G | H1' | 2.6 | 4.14 |
| 4 | A | H2 | 23 | U | H1' | 4.44 | 7.08 |
| 4 | A | H2 | 23 | U | H3 | 1.06 | 2.48 |
| 4 | A | H2 | 24 | G | H1' | 2.59 | 4.13 |

| | | | | | | | |
|---|---|---|---|---|---|---|---|
| 4 | A | H61 | 23 | U | O4 | 1.8 | 2.4 |
| 4 | A | H8 | 3 | C | H2' | 1.83 | 2.92 |
| 4 | A | H8 | 4 | A | H1' | 2.95 | 4.7 |
| 4 | A | H8 | 4 | A | H2' | 2.93 | 4.68 |
| 4 | A | H8 | 4 | A | H3' | 2.61 | 4.17 |
| 4 | A | N1 | 23 | U | H3 | 1.8 | 2.4 |
| 5 | G | H1 | 22 | C | N3 | 1.8 | 2.4 |
| 5 | G | H1' | 4 | A | H2' | 3.08 | 4.91 |
| 5 | G | H1' | 5 | G | H3' | 2.88 | 4.59 |
| 5 | G | H22 | 22 | C | O2 | 1.8 | 2.4 |
| 5 | G | H8 | 4 | A | H2' | 1.87 | 2.97 |
| 5 | G | H8 | 4 | A | H3' | 3.03 | 4.84 |
| 5 | G | H8 | 5 | G | H1' | 2.94 | 4.69 |
| 5 | G | H8 | 5 | G | H2' | 2.93 | 4.68 |
| 5 | G | H8 | 5 | G | H3' | 2.35 | 3.76 |
| 5 | G | H8 | 6 | C | H5 | 3.28 | 5.23 |
| 5 | G | O6 | 22 | C | H41 | 1.8 | 2.4 |
| 6 | C | H1' | 5 | G | H1 | 3 | 6 |
| 6 | C | H1' | 6 | C | H2' | 2.06 | 3.29 |
| 6 | C | H1' | 6 | C | H3' | 3.67 | 5.86 |
| 6 | C | H41 | 21 | G | H1 | 1.2 | 2.8 |
| 6 | C | H41 | 21 | G | O6 | 1.8 | 2.4 |
| 6 | C | H5 | 5 | G | H2' | 3 | 4.79 |
| 6 | C | H6 | 5 | G | H2' | 1.78 | 2.85 |
| 6 | C | H6 | 5 | G | H3' | 2.29 | 3.65 |
| 6 | C | H6 | 6 | C | H1' | 2.59 | 4.13 |
| 6 | C | H6 | 6 | C | H2' | 2.63 | 4.2 |
| 6 | C | H6 | 6 | C | H3' | 1.95 | 3.11 |
| 6 | C | N3 | 21 | G | H1 | 1.8 | 2.4 |
| 6 | C | O2 | 21 | G | H22 | 1.8 | 2.4 |
| 7 | A | H1' | 6 | C | H2' | 3.11 | 4.95 |
| 7 | A | H1' | 7 | A | H2' | 2.09 | 3.34 |
| 7 | A | H1' | 7 | A | H3' | 2.99 | 4.76 |
| 7 | A | H1' | 7 | A | H4' | 2.98 | 4.75 |
| 7 | A | H1' | 21 | G | H1 | 3 | 6 |
| 7 | A | H2 | 7 | A | H1' | 3.75 | 5.99 |
| 7 | A | H2 | 8 | G | H1 | 3 | 6 |
| 7 | A | H2 | 8 | G | H1' | 2.73 | 4.36 |
| 7 | A | H2 | 21 | G | H1 | 3 | 6 |
| 7 | A | H2 | 21 | G | H1' | 2.78 | 4.43 |
| 7 | A | H8 | 6 | C | H2' | 1.85 | 2.95 |
| 7 | A | H8 | 7 | A | H1' | 2.58 | 4.12 |
| 8 | G | H1 | 18 | G | H1 | 1.53 | 3.55 |

| | | | | | | | |
|---|---|---|---|---|---|---|---|
| 8 | G | H1 | 19 | C | N3 | 1.8 | 2.4 |
| 8 | G | H1' | 8 | G | H3' | 2.82 | 4.5 |
| 8 | G | H22 | 19 | C | O2 | 1.8 | 2.4 |
| 8 | G | H8 | 7 | A | H2' | 2.01 | 3.21 |
| 8 | G | H8 | 8 | G | H1' | 2.77 | 4.42 |
| 8 | G | H8 | 8 | G | H2' | 2.91 | 4.65 |
| 8 | G | H8 | 8 | G | H3' | 2.28 | 3.64 |
| 8 | G | H8 | 9 | C | H5 | 3.49 | 5.57 |
| 8 | G | O6 | 19 | C | H41 | 1.8 | 2.4 |
| 9 | C | H1' | 8 | G | H1 | 3 | 6 |
| 9 | C | H1' | 8 | G | H2' | 2.88 | 4.6 |
| 9 | C | H1' | 9 | C | H2' | 2.04 | 3.25 |
| 9 | C | H41 | 18 | G | H1 | 2 | 4.5 |
| 9 | C | H41 | 18 | G | O6 | 1.8 | 2.4 |
| 9 | C | H5 | 8 | G | H2' | 3.63 | 5.8 |
| 9 | C | H6 | 9 | C | H1' | 2.64 | 4.21 |
| 9 | C | H6 | 9 | C | H2' | 2.2 | 3.5 |
| 9 | C | H6 | 10 | U | H5 | 3.2 | 5.1 |
| 9 | C | N3 | 18 | G | H1 | 1.8 | 2.4 |
| 9 | C | O2 | 18 | G | H22 | 1.8 | 2.4 |
| 10 | U | H1' | 9 | C | H2' | 3.29 | 5.25 |
| 10 | U | H1' | 10 | U | H2' | 2.2 | 3.5 |
| 10 | U | H1' | 10 | U | H3' | 2.88 | 4.6 |
| 10 | U | H1' | 10 | U | H4' | 2.69 | 4.29 |
| 10 | U | H1' | 18 | G | H1 | 3 | 6 |
| 10 | U | H3 | 11 | G | H1 | 2.39 | 5.59 |
| 10 | U | H3 | 17 | A | N1 | 1.8 | 2.4 |
| 10 | U | H5 | 10 | U | H3' | 4.09 | 6.51 |
| 10 | U | H6 | 9 | C | H2' | 1.72 | 2.75 |
| 10 | U | H6 | 10 | U | H1' | 2.52 | 4.02 |
| 10 | U | H6 | 10 | U | H3' | 1.94 | 3.1 |
| 10 | U | O4 | 17 | A | H61 | 1.8 | 2.4 |
| 11 | G | H1 | 16 | C | N3 | 1.8 | 2.4 |
| 11 | G | H1' | 10 | U | H2' | 3.42 | 5.46 |
| 11 | G | H1' | 10 | U | H3 | 3 | 7 |
| 11 | G | H1' | 11 | G | H2' | 2.12 | 3.39 |
| 11 | G | H1' | 11 | G | H3' | 3 | 4.79 |
| 11 | G | H1' | 11 | G | H4' | 2.92 | 4.65 |
| 11 | G | H22 | 16 | C | O2 | 1.8 | 2.4 |
| 11 | G | H8 | 10 | U | H2' | 1.87 | 2.97 |
| 11 | G | H8 | 11 | G | H1' | 2.99 | 4.77 |
| 11 | G | O6 | 16 | C | H41 | 1.8 | 2.4 |
| 12 | U | H1' | 11 | G | H1 | 3 | 6 |

| | | | | | | | |
|---|---|---|---|---|---|---|---|
| 12 | U | H1' | 12 | U | H2' | 2.14 | 3.42 |
| 12 | U | H1' | 12 | U | H4' | 2.61 | 4.17 |
| 12 | U | H3 | 11 | G | H1 | 1.46 | 3.4 |
| 12 | U | H3 | 15 | A | H2 | 1.08 | 2.52 |
| 12 | U | H3 | 15 | A | N1 | 1.8 | 2.4 |
| 12 | U | H6 | 11 | G | H2' | 1.84 | 2.94 |
| 12 | U | H6 | 12 | U | H1' | 2.62 | 4.18 |
| 12 | U | H6 | 12 | U | H2' | 2.71 | 4.32 |
| 12 | U | H6 | 12 | U | H3' | 2.01 | 3.21 |
| 12 | U | H6 | 13 | C | H5 | 3.13 | 4.99 |
| 12 | U | O4 | 15 | A | H61 | 1.8 | 2.4 |
| 13 | C | H1' | 12 | U | H2' | 3.24 | 5.17 |
| 13 | C | H1' | 13 | C | H2' | 2.08 | 3.31 |
| 13 | C | H41 | 14 | G | O6 | 1.8 | 2.4 |
| 13 | C | H5 | 12 | U | H2' | 2.69 | 4.29 |
| 13 | C | H5 | 12 | U | H3' | 3.4 | 5.42 |
| 13 | C | H5 | 12 | U | H3' | 3.4 | 5.42 |
| 13 | C | H6 | 12 | U | H2' | 1.88 | 3 |
| 13 | C | H6 | 13 | C | H1' | 2.71 | 4.32 |
| 13 | C | H6 | 13 | C | H2' | 2.47 | 3.94 |
| 13 | C | N3 | 14 | G | H1 | 1.8 | 2.4 |
| 13 | C | O2 | 14 | G | H22 | 1.8 | 2.4 |
| 14 | G | H1' | 14 | G | H3' | 3.21 | 5.13 |
| 14 | G | H1' | 14 | G | H4' | 2.98 | 4.75 |
| 14 | G | H8 | 14 | G | H1' | 2.65 | 4.23 |
| 14 | G | H8 | 14 | G | H2' | 2.62 | 4.18 |
| 14 | G | H8 | 14 | G | H3' | 2.31 | 3.69 |
| 14 | G | H8 | 14 | G | H4' | 3.06 | 4.88 |
| 15 | A | H1' | 14 | G | H2' | 3.44 | 5.48 |
| 15 | A | H1' | 15 | A | H3' | 3.83 | 6.11 |
| 15 | A | H2 | 13 | C | H1' | 2.89 | 4.61 |
| 15 | A | H2 | 15 | A | H1' | 3.82 | 6.1 |
| 15 | A | H2 | 16 | C | H1' | 2.74 | 4.37 |
| 15 | A | H8 | 14 | G | H2' | 2.25 | 3.59 |
| 15 | A | H8 | 15 | A | H1' | 3.06 | 4.88 |
| 15 | A | H8 | 16 | C | H5 | 4.12 | 6.56 |
| 16 | C | H1' | 16 | C | H2' | 2.23 | 3.55 |
| 16 | C | H1' | 16 | C | H4' | 2.8 | 4.47 |
| 16 | C | H41 | 11 | G | H1 | 1.21 | 2.81 |
| 16 | C | H41 | 12 | U | H3 | 1.55 | 3.61 |
| 16 | C | H5 | 15 | A | H2' | 3.38 | 5.39 |
| 16 | C | H6 | 15 | A | H2' | 1.71 | 2.72 |
| 16 | C | H6 | 15 | A | H3' | 3.25 | 5.19 |

| | | | | | | | |
|---|---|---|---|---|---|---|---|
| 16 | C | H6 | 16 | C | H1' | 2.63 | 4.2 |
| 16 | C | H6 | 16 | C | H2' | 2.72 | 4.34 |
| 17 | A | H1' | 11 | G | H1 | 3 | 6 |
| 17 | A | H1' | 16 | C | H2' | 3.58 | 5.71 |
| 17 | A | H1' | 17 | A | H2' | 2.27 | 3.62 |
| 17 | A | H1' | 17 | A | H3' | 3.84 | 6.12 |
| 17 | A | H1' | 17 | A | H4' | 3.39 | 5.41 |
| 17 | A | H2 | 10 | U | H1' | 4.44 | 7.08 |
| 17 | A | H2 | 10 | U | H3 | 1.06 | 2.48 |
| 17 | A | H2 | 11 | G | H1' | 2.59 | 4.13 |
| 17 | A | H2 | 17 | A | H1' | 4.08 | 6.51 |
| 17 | A | H2 | 18 | G | H1' | 2.6 | 4.14 |
| 17 | A | H8 | 16 | C | H2' | 1.83 | 2.92 |
| 17 | A | H8 | 17 | A | H1' | 2.95 | 4.7 |
| 17 | A | H8 | 17 | A | H2' | 2.93 | 4.68 |
| 17 | A | H8 | 17 | A | H3' | 2.61 | 4.17 |
| 18 | G | H1' | 17 | A | H2' | 3.08 | 4.91 |
| 18 | G | H1' | 18 | G | H3' | 2.88 | 4.59 |
| 18 | G | H8 | 17 | A | H2' | 1.87 | 2.97 |
| 18 | G | H8 | 17 | A | H3' | 3.03 | 4.84 |
| 18 | G | H8 | 18 | G | H1' | 2.94 | 4.69 |
| 18 | G | H8 | 18 | G | H2' | 2.93 | 4.68 |
| 18 | G | H8 | 18 | G | H3' | 2.35 | 3.76 |
| 18 | G | H8 | 19 | C | H5 | 3.28 | 5.23 |
| 19 | C | H1' | 18 | G | H1 | 3 | 6 |
| 19 | C | H1' | 19 | C | H2' | 2.06 | 3.29 |
| 19 | C | H1' | 19 | C | H3' | 3.67 | 5.86 |
| 19 | C | H41 | 8 | G | H1 | 1.2 | 2.8 |
| 19 | C | H5 | 18 | G | H2' | 3 | 4.79 |
| 19 | C | H6 | 18 | G | H2' | 1.78 | 2.85 |
| 19 | C | H6 | 18 | G | H3' | 2.29 | 3.65 |
| 19 | C | H6 | 19 | C | H1' | 2.59 | 4.13 |
| 19 | C | H6 | 19 | C | H2' | 2.63 | 4.2 |
| 19 | C | H6 | 19 | C | H3' | 1.95 | 3.11 |
| 20 | A | H1' | 8 | G | H1 | 3 | 6 |
| 20 | A | H1' | 19 | C | H2' | 3.11 | 4.95 |
| 20 | A | H1' | 20 | A | H2' | 2.09 | 3.34 |
| 20 | A | H1' | 20 | A | H3' | 2.99 | 4.76 |
| 20 | A | H1' | 20 | A | H4' | 2.98 | 4.75 |
| 20 | A | H2 | 8 | G | H1 | 3 | 6 |
| 20 | A | H2 | 8 | G | H1' | 2.78 | 4.43 |
| 20 | A | H2 | 20 | A | H1' | 3.75 | 5.99 |
| 20 | A | H2 | 21 | G | H1 | 3 | 6 |

| | | | | | | | |
|---|---|---|---|---|---|---|---|
| 20 | A | H2 | 21 | G | H1' | 2.73 | 4.36 |
| 20 | A | H8 | 19 | C | H2' | 1.85 | 2.95 |
| 20 | A | H8 | 20 | A | H1' | 2.58 | 4.12 |
| 21 | G | H1 | 5 | G | H1 | 1.53 | 3.55 |
| 21 | G | H1' | 21 | G | H3' | 2.82 | 4.5 |
| 21 | G | H8 | 20 | A | H2' | 2.01 | 3.21 |
| 21 | G | H8 | 21 | G | H1' | 2.77 | 4.42 |
| 21 | G | H8 | 21 | G | H2' | 2.91 | 4.65 |
| 21 | G | H8 | 21 | G | H3' | 2.28 | 3.64 |
| 21 | G | H8 | 22 | C | H5 | 3.49 | 5.57 |
| 22 | C | H1' | 21 | G | H1 | 3 | 6 |
| 22 | C | H1' | 21 | G | H2' | 2.88 | 4.6 |
| 22 | C | H1' | 22 | C | H2' | 2.04 | 3.25 |
| 22 | C | H41 | 5 | G | H1 | 2 | 4.5 |
| 22 | C | H5 | 21 | G | H2' | 3.63 | 5.8 |
| 22 | C | H6 | 22 | C | H1' | 2.64 | 4.21 |
| 22 | C | H6 | 22 | C | H2' | 2.2 | 3.5 |
| 22 | C | H6 | 23 | U | H5 | 3.2 | 5.1 |
| 23 | U | H1' | 5 | G | H1 | 3 | 6 |
| 23 | U | H1' | 22 | C | H2' | 3.29 | 5.25 |
| 23 | U | H1' | 23 | U | H2' | 2.2 | 3.5 |
| 23 | U | H1' | 23 | U | H3' | 2.88 | 4.6 |
| 23 | U | H1' | 23 | U | H4' | 2.69 | 4.29 |
| 23 | U | H3 | 24 | G | H1 | 2.39 | 5.59 |
| 23 | U | H5 | 23 | U | H3' | 4.09 | 6.51 |
| 23 | U | H6 | 22 | C | H2' | 1.72 | 2.75 |
| 23 | U | H6 | 23 | U | H1' | 2.52 | 4.02 |
| 23 | U | H6 | 23 | U | H3' | 1.94 | 3.1 |
| 24 | G | H1' | 23 | U | H2' | 3.42 | 5.46 |
| 24 | G | H1' | 23 | U | H3 | 3 | 7 |
| 24 | G | H1' | 24 | G | H2' | 2.12 | 3.39 |
| 24 | G | H1' | 24 | G | H3' | 3 | 4.79 |
| 24 | G | H1' | 24 | G | H4' | 2.92 | 4.65 |
| 24 | G | H8 | 23 | U | H2' | 1.87 | 2.97 |
| 24 | G | H8 | 24 | G | H1' | 2.99 | 4.77 |
| 25 | U | H1' | 24 | G | H1 | 3 | 6 |
| 25 | U | H1' | 25 | U | H2' | 2.14 | 3.42 |
| 25 | U | H1' | 25 | U | H4' | 2.61 | 4.17 |
| 25 | U | H3 | 2 | A | H2 | 1.08 | 2.52 |
| 25 | U | H3 | 24 | G | H1 | 1.46 | 3.4 |
| 25 | U | H6 | 24 | G | H2' | 1.84 | 2.94 |
| 25 | U | H6 | 25 | U | H1' | 2.62 | 4.18 |
| 25 | U | H6 | 25 | U | H2' | 2.71 | 4.32 |

| | | | | | | | |
|---|---|---|---|---|---|---|---|
| 25 | U | H6 | 25 | U | H3' | 2.01 | 3.21 |
| 25 | U | H6 | 26 | C | H5 | 3.13 | 4.99 |
| 26 | C | H1' | 25 | U | H2' | 3.24 | 5.17 |
| 26 | C | H1' | 26 | C | H2' | 2.08 | 3.31 |
| 26 | C | H5 | 25 | U | H2' | 2.69 | 4.29 |
| 26 | C | H5 | 25 | U | H3' | 3.4 | 5.42 |
| 26 | C | H5 | 25 | U | H3' | 3.4 | 5.42 |
| 26 | C | H6 | 25 | U | H2' | 1.88 | 3 |
| 26 | C | H6 | 26 | C | H1' | 2.71 | 4.32 |
| 26 | C | H6 | 26 | C | H2' | 2.47 | 3.94 |

**Table S6:** Dihedral angle restraints used for modeling of the *apo* r(CAG) duplex.

| | | | |
|---|---|---|---|
| ALPHA | (1 RG5 O3')-(2 RA P)-(2 RA O5')-(2 RA C5') | -155.0 | 25.0 |
| ALPHA | (2 RA O3')-(3 RC P)-(3 RC O5')-(3 RC C5') | -155.0 | 25.0 |
| ALPHA | (3 RC O3')-(4 RA P)-(4 RA O5')-(4 RA C5') | -155.0 | 25.0 |
| ALPHA | (4 RA O3')-(5 RG P)-(5 RG O5')-(5 RG C5') | -155.0 | 25.0 |
| ALPHA | (5 RG O3')-(6 RC P)-(6 RC O5')-(6 RC C5') | -155.0 | 25.0 |
| ALPHA | (7 RA O3')-(8 RG P)-(8 RG O5')-(8 RG C5') | -155.0 | 25.0 |
| ALPHA | (8 RG O3')-(9 RC P)-(9 RC O5')-(9 RC C5') | -155.0 | 25.0 |
| ALPHA | (9 RC O3')-(10 RU P)-(10 RU O5')-(10 RU C5') | -155.0 | 25.0 |
| ALPHA | (10 RU O3')-(11 RG P)-(11 RG O5')-(11 RG C5') | -155.0 | 25.0 |
| ALPHA | (11 RG O3')-(12 RU P)-(12 RU O5')-(12 RU C5') | -155.0 | 25.0 |
| ALPHA | (14 RG5 O3')-(15 RA P)-(15 RA O5')-(15 RA C5') | -155.0 | 25.0 |
| ALPHA | (15 RA O3')-(16 RC P)-(16 RC O5')-(16 RC C5') | -155.0 | 25.0 |
| ALPHA | (16 RC O3')-(17 RA P)-(17 RA O5')-(17 RA C5') | -155.0 | 25.0 |
| ALPHA | (17 RA O3')-(18 RG P)-(18 RG O5')-(18 RG C5') | -155.0 | 25.0 |
| ALPHA | (18 RG O3')-(19 RC P)-(19 RC O5')-(19 RC C5') | -155.0 | 25.0 |
| ALPHA | (20 RA O3')-(21 RG P)-(21 RG O5')-(21 RG C5') | -155.0 | 25.0 |
| ALPHA | (21 RG O3')-(22 RC P)-(22 RC O5')-(22 RC C5') | -155.0 | 25.0 |
| ALPHA | (22 RC O3')-(23 RU P)-(23 RU O5')-(23 RU C5') | -155.0 | 25.0 |
| ALPHA | (23 RU O3')-(24 RG P)-(24 RG O5')-(24 RG C5') | -155.0 | 25.0 |
| ALPHA | (24 RG O3')-(25 RU P)-(25 RU O5')-(25 RU C5') | -155.0 | 25.0 |
| BETA | (2 RA P)-(2 RA O5')-(2 RA C5')-(2 RA C4') | 90.0 | 240.0 |
| BETA | (3 RC P)-(3 RC O5')-(3 RC C5')-(3 RC C4') | 90.0 | 240.0 |
| BETA | (4 RA P)-(4 RA O5')-(4 RA C5')-(4 RA C4') | 90.0 | 240.0 |
| BETA | (5 RG P)-(5 RG O5')-(5 RG C5')-(5 RG C4') | 90.0 | 240.0 |
| BETA | (6 RC P)-(6 RC O5')-(6 RC C5')-(6 RC C4') | 90.0 | 240.0 |
| BETA | (8 RG P)-(8 RG O5')-(8 RG C5')-(8 RG C4') | 90.0 | 240.0 |
| BETA | (9 RC P)-(9 RC O5')-(9 RC C5')-(9 RC C4') | 90.0 | 240.0 |
| BETA | (10 RU P)-(10 RU O5')-(10 RU C5')-(10 RU C4') | 90.0 | 240.0 |
| BETA | (11 RG P)-(11 RG O5')-(11 RG C5')-(11 RG C4') | 90.0 | 240.0 |
| BETA | (12 RU P)-(12 RU O5')-(12 RU C5')-(12 RU C4') | 90.0 | 240.0 |
| BETA | (15 RA P)-(15 RA O5')-(15 RA C5')-(15 RA C4') | 90.0 | 240.0 |
| BETA | (16 RC P)-(16 RC O5')-(16 RC C5')-(16 RC C4') | 90.0 | 240.0 |
| BETA | (17 RA P)-(17 RA O5')-(17 RA C5')-(17 RA C4') | 90.0 | 240.0 |
| BETA | (18 RG P)-(18 RG O5')-(18 RG C5')-(18 RG C4') | 90.0 | 240.0 |
| BETA | (19 RC P)-(19 RC O5')-(19 RC C5')-(19 RC C4') | 90.0 | 240.0 |
| BETA | (21 RG P)-(21 RG O5')-(21 RG C5')-(21 RG C4') | 90.0 | 240.0 |
| BETA | (22 RC P)-(22 RC O5')-(22 RC C5')-(22 RC C4') | 90.0 | 240.0 |
| BETA | (23 RU P)-(23 RU O5')-(23 RU C5')-(23 RU C4') | 90.0 | 240.0 |
| BETA | (24 RG P)-(24 RG O5')-(24 RG C5')-(24 RG C4') | 90.0 | 240.0 |
| BETA | (25 RU P)-(25 RU O5')-(25 RU C5')-(25 RU C4') | 90.0 | 240.0 |
| GAMMA | (2 RA O5')-(2 RA C5')-(2 RA C4')-(2 RA C3') | 0.0 | 120.0 |

| | | | |
|---|---|---|---|
| GAMMA | (3 RC O5')-(3 RC C5')-(3 RC C4')-(3 RC C3') | 0.0 | 120.0 |
| GAMMA | (4 RA O5')-(4 RA C5')-(4 RA C4')-(4 RA C3') | 0.0 | 120.0 |
| GAMMA | (5 RG O5')-(5 RG C5')-(5 RG C4')-(5 RG C3') | 0.0 | 120.0 |
| GAMMA | (6 RC O5')-(6 RC C5')-(6 RC C4')-(6 RC C3') | 0.0 | 120.0 |
| GAMMA | (8 RG O5')-(8 RG C5')-(8 RG C4')-(8 RG C3') | 0.0 | 120.0 |
| GAMMA | (9 RC O5')-(9 RC C5')-(9 RC C4')-(9 RC C3') | 0.0 | 120.0 |
| GAMMA | (10 RU O5')-(10 RU C5')-(10 RU C4')-(10 RU C3') | 0.0 | 120.0 |
| GAMMA | (11 RG O5')-(11 RG C5')-(11 RG C4')-(11 RG C3') | 0.0 | 120.0 |
| GAMMA | (12 RU O5')-(12 RU C5')-(12 RU C4')-(12 RU C3') | 0.0 | 120.0 |
| GAMMA | (15 RA O5')-(15 RA C5')-(15 RA C4')-(15 RA C3') | 0.0 | 120.0 |
| GAMMA | (16 RC O5')-(16 RC C5')-(16 RC C4')-(16 RC C3') | 0.0 | 120.0 |
| GAMMA | (17 RA O5')-(17 RA C5')-(17 RA C4')-(17 RA C3') | 0.0 | 120.0 |
| GAMMA | (18 RG O5')-(18 RG C5')-(18 RG C4')-(18 RG C3') | 0.0 | 120.0 |
| GAMMA | (19 RC O5')-(19 RC C5')-(19 RC C4')-(19 RC C3') | 0.0 | 120.0 |
| GAMMA | (21 RG O5')-(21 RG C5')-(21 RG C4')-(21 RG C3') | 0.0 | 120.0 |
| GAMMA | (22 RC O5')-(22 RC C5')-(22 RC C4')-(22 RC C3') | 0.0 | 120.0 |
| GAMMA | (23 RU O5')-(23 RU C5')-(23 RU C4')-(23 RU C3') | 0.0 | 120.0 |
| GAMMA | (24 RG O5')-(24 RG C5')-(24 RG C4')-(24 RG C3') | 0.0 | 120.0 |
| GAMMA | (25 RU O5')-(25 RU C5')-(25 RU C4')-(25 RU C3') | 0.0 | 120.0 |
| DELTA | (2 RA C5')-(2 RA C4')-(2 RA C3')-(2 RA O3') | 45.0 | 115.0 |
| DELTA | (3 RC C5')-(3 RC C4')-(3 RC C3')-(3 RC O3') | 45.0 | 115.0 |
| DELTA | (4 RA C5')-(4 RA C4')-(4 RA C3')-(4 RA O3') | 45.0 | 115.0 |
| DELTA | (5 RG C5')-(5 RG C4')-(5 RG C3')-(5 RG O3') | 45.0 | 115.0 |
| DELTA | (6 RC C5')-(6 RC C4')-(6 RC C3')-(6 RC O3') | 45.0 | 115.0 |
| DELTA | (8 RG C5')-(8 RG C4')-(8 RG C3')-(8 RG O3') | 45.0 | 115.0 |
| DELTA | (9 RC C5')-(9 RC C4')-(9 RC C3')-(9 RC O3') | 45.0 | 115.0 |
| DELTA | (10 RU C5')-(10 RU C4')-(10 RU C3')-(10 RU O3') | 45.0 | 115.0 |
| DELTA | (11 RG C5')-(11 RG C4')-(11 RG C3')-(11 RG O3') | 45.0 | 115.0 |
| DELTA | (12 RU C5')-(12 RU C4')-(12 RU C3')-(12 RU O3') | 45.0 | 115.0 |
| DELTA | (15 RA C5')-(15 RA C4')-(15 RA C3')-(15 RA O3') | 45.0 | 115.0 |
| DELTA | (16 RC C5')-(16 RC C4')-(16 RC C3')-(16 RC O3') | 45.0 | 115.0 |
| DELTA | (17 RA C5')-(17 RA C4')-(17 RA C3')-(17 RA O3') | 45.0 | 115.0 |
| DELTA | (18 RG C5')-(18 RG C4')-(18 RG C3')-(18 RG O3') | 45.0 | 115.0 |
| DELTA | (19 RC C5')-(19 RC C4')-(19 RC C3')-(19 RC O3') | 45.0 | 115.0 |
| DELTA | (21 RG C5')-(21 RG C4')-(21 RG C3')-(21 RG O3') | 45.0 | 115.0 |
| DELTA | (22 RC C5')-(22 RC C4')-(22 RC C3')-(22 RC O3') | 45.0 | 115.0 |
| DELTA | (23 RU C5')-(23 RU C4')-(23 RU C3')-(23 RU O3') | 45.0 | 115.0 |
| DELTA | (24 RG C5')-(24 RG C4')-(24 RG C3')-(24 RG O3') | 45.0 | 115.0 |
| DELTA | (25 RU C5')-(25 RU C4')-(25 RU C3')-(25 RU O3') | 45.0 | 115.0 |
| EPSILON | (2 RA C4')-(2 RA C3')-(2 RA O3')-(3 RC P) | -240.0 | 10.0 |
| EPSILON | (3 RC C4')-(3 RC C3')-(3 RC O3')-(4 RA P) | -240.0 | 10.0 |
| EPSILON | (4 RA C4')-(4 RA C3')-(4 RA O3')-(5 RG P) | -240.0 | 10.0 |
| EPSILON | (5 RG C4')-(5 RG C3')-(5 RG O3')-(6 RC P) | -240.0 | 10.0 |

| | | | |
|---|---|---|---|
| EPSILON | (6 RC C4')-(6 RC C3')-(6 RC O3')-(7 RA P) | -240.0 | 10.0 |
| EPSILON | (8 RG C4')-(8 RG C3')-(8 RG O3')-(9 RC P) | -240.0 | 10.0 |
| EPSILON | (9 RC C4')-(9 RC C3')-(9 RC O3')-(10 RU P) | -240.0 | 10.0 |
| EPSILON | (10 RU C4')-(10 RU C3')-(10 RU O3')-(11 RG P) | -240.0 | 10.0 |
| EPSILON | (11 RG C4')-(11 RG C3')-(11 RG O3')-(12 RU P) | -240.0 | 10.0 |
| EPSILON | (12 RU C4')-(12 RU C3')-(12 RU O3')-(13 RC3 P) | -240.0 | 10.0 |
| EPSILON | (15 RA C4')-(15 RA C3')-(15 RA O3')-(16 RC P) | -240.0 | 10.0 |
| EPSILON | (16 RC C4')-(16 RC C3')-(16 RC O3')-(17 RA P) | -240.0 | 10.0 |
| EPSILON | (17 RA C4')-(17 RA C3')-(17 RA O3')-(18 RG P) | -240.0 | 10.0 |
| EPSILON | (18 RG C4')-(18 RG C3')-(18 RG O3')-(19 RC P) | -240.0 | 10.0 |
| EPSILON | (19 RC C4')-(19 RC C3')-(19 RC O3')-(20 RA P) | -240.0 | 10.0 |
| EPSILON | (21 RG C4')-(21 RG C3')-(21 RG O3')-(22 RC P) | -240.0 | 10.0 |
| EPSILON | (22 RC C4')-(22 RC C3')-(22 RC O3')-(23 RU P) | -240.0 | 10.0 |
| EPSILON | (23 RU C4')-(23 RU C3')-(23 RU O3')-(24 RG P) | -240.0 | 10.0 |
| EPSILON | (24 RG C4')-(24 RG C3')-(24 RG O3')-(25 RU P) | -240.0 | 10.0 |
| EPSILON | (25 RU C4')-(25 RU C3')-(25 RU O3')-(26 RC3 P) | -240.0 | 10.0 |
| ZETA | (2 RA C3')-(2 RA O3')-(3 RC P)-(3 RC O5') | -160.0 | 20.0 |
| ZETA | (3 RC C3')-(3 RC O3')-(4 RA P)-(4 RA O5') | -160.0 | 20.0 |
| ZETA | (4 RA C3')-(4 RA O3')-(5 RG P)-(5 RG O5') | -160.0 | 20.0 |
| ZETA | (5 RG C3')-(5 RG O3')-(6 RC P)-(6 RC O5') | -160.0 | 20.0 |
| ZETA | (6 RC C3')-(6 RC O3')-(7 RA P)-(7 RA O5') | -160.0 | 20.0 |
| ZETA | (8 RG C3')-(8 RG O3')-(9 RC P)-(9 RC O5') | -160.0 | 20.0 |
| ZETA | (9 RC C3')-(9 RC O3')-(10 RU P)-(10 RU O5') | -160.0 | 20.0 |
| ZETA | (10 RU C3')-(10 RU O3')-(11 RG P)-(11 RG O5') | -160.0 | 20.0 |
| ZETA | (11 RG C3')-(11 RG O3')-(12 RU P)-(12 RU O5') | -160.0 | 20.0 |
| ZETA | (12 RU C3')-(12 RU O3')-(13 RC3 P)-(13 RC3 O5') | -160.0 | 20.0 |
| ZETA | (15 RA C3')-(15 RA O3')-(16 RC P)-(16 RC O5') | -160.0 | 20.0 |
| ZETA | (16 RC C3')-(16 RC O3')-(17 RA P)-(17 RA O5') | -160.0 | 20.0 |
| ZETA | (17 RA C3')-(17 RA O3')-(18 RG P)-(18 RG O5') | -160.0 | 20.0 |
| ZETA | (18 RG C3')-(18 RG O3')-(19 RC P)-(19 RC O5') | -160.0 | 20.0 |
| ZETA | (19 RC C3')-(19 RC O3')-(20 RA P)-(20 RA O5') | -160.0 | 20.0 |
| ZETA | (21 RG C3')-(21 RG O3')-(22 RC P)-(22 RC O5') | -160.0 | 20.0 |
| ZETA | (22 RC C3')-(22 RC O3')-(23 RU P)-(23 RU O5') | -160.0 | 20.0 |
| ZETA | (23 RU C3')-(23 RU O3')-(24 RG P)-(24 RG O5') | -160.0 | 20.0 |
| ZETA | (24 RG C3')-(24 RG O3')-(25 RU P)-(25 RU O5') | -160.0 | 20.0 |
| ZETA | (25 RU C3')-(25 RU O3')-(26 RC3 P)-(26 RC3 O5') | -160.0 | 20.0 |
| CHI | (2 RA O4')-(2 RA C1')-(2 RA N9)-(2 RA C4) | 170.0 | 340.0 |
| CHI | (3 RC O4')-(3 RC C1')-(3 RC N1)-(3 RC C2) | 170.0 | 340.0 |
| CHI | (4 RA O4')-(4 RA C1')-(4 RA N9)-(4 RA C4) | 170.0 | 340.0 |
| CHI | (5 RG O4')-(5 RG C1')-(5 RG N9)-(5 RG C4) | 170.0 | 340.0 |
| CHI | (6 RC O4')-(6 RC C1')-(6 RC N1)-(6 RC C2) | 170.0 | 340.0 |
| CHI | (8 RG O4')-(8 RG C1')-(8 RG N9)-(8 RG C4) | 170.0 | 340.0 |
| CHI | (9 RC O4')-(9 RC C1')-(9 RC N1)-(9 RC C2) | 170.0 | 340.0 |

| | | | |
|---|---|---|---|
| CHI | (10 RU O4')-(10 RU C1')-(10 RU N1)-(10 RU C2) | 170.0 | 340.0 |
| CHI | (11 RG O4')-(11 RG C1')-(11 RG N9)-(11 RG C4) | 170.0 | 340.0 |
| CHI | (12 RU O4')-(12 RU C1')-(12 RU N1)-(12 RU C2) | 170.0 | 340.0 |
| CHI | (15 RA O4')-(15 RA C1')-(15 RA N9)-(15 RA C4) | 170.0 | 340.0 |
| CHI | (16 RC O4')-(16 RC C1')-(16 RC N1)-(16 RC C2) | 170.0 | 340.0 |
| CHI | (17 RA O4')-(17 RA C1')-(17 RA N9)-(17 RA C4) | 170.0 | 340.0 |
| CHI | (18 RG O4')-(18 RG C1')-(18 RG N9)-(18 RG C4) | 170.0 | 340.0 |
| CHI | (19 RC O4')-(19 RC C1')-(19 RC N1)-(19 RC C2) | 170.0 | 340.0 |
| CHI | (21 RG O4')-(21 RG C1')-(21 RG N9)-(21 RG C4) | 170.0 | 340.0 |
| CHI | (22 RC O4')-(22 RC C1')-(22 RC N1)-(22 RC C2) | 170.0 | 340.0 |
| CHI | (23 RU O4')-(23 RU C1')-(23 RU N1)-(23 RU C2) | 170.0 | 340.0 |
| CHI | (24 RG O4')-(24 RG C1')-(24 RG N9)-(24 RG C4) | 170.0 | 340.0 |
| CHI | (25 RU O4')-(25 RU C1')-(25 RU N1)-(25 RU C2) | 170.0 | 340.0 |

**Table S7:** NOE restraints used for modeling of the r(CAG)-**1** complex.

| | | | | | | | |
|---|---|---|---|---|---|---|---|
| 1 | G | H1 | 26 | C | N3 | 1.8 | 2.4 |
| 1 | G | H1' | 1 | G | H2' | 2.42 | 3.86 |
| 1 | G | H1' | 1 | G | H3' | 2.84 | 4.52 |
| 1 | G | H1' | 1 | G | H4' | 2.56 | 4.08 |
| 1 | G | H22 | 26 | C | O2 | 1.8 | 2.4 |
| 1 | G | H8 | 1 | G | H1' | 2.57 | 4.1 |
| 1 | G | O6 | 26 | C | H41 | 1.8 | 2.4 |
| 2 | A | H1' | 1 | G | H2' | 3.18 | 5.06 |
| 2 | A | H1' | 2 | A | H2' | 2 | 3.19 |
| 2 | A | H2 | 2 | A | H1' | 3.8 | 6.06 |
| 2 | A | H2 | 3 | C | H1' | 2.73 | 4.35 |
| 2 | A | H2 | 26 | C | H1' | 2.67 | 4.26 |
| 2 | A | H61 | 25 | U | O4 | 1.8 | 2.4 |
| 2 | A | H8 | 1 | G | H1' | 3.11 | 4.96 |
| 2 | A | H8 | 1 | G | H2' | 2.45 | 3.91 |
| 2 | A | H8 | 2 | A | H1' | 3.07 | 4.89 |
| 2 | A | N1 | 25 | U | H3 | 1.8 | 2.4 |
| 3 | C | H1' | 2 | A | H2' | 2.59 | 4.13 |
| 3 | C | H1' | 3 | C | H2' | 2.01 | 3.2 |
| 3 | C | H1' | 3 | C | H4' | 2.74 | 4.37 |
| 3 | C | H41 | 24 | G | O6 | 1.8 | 2.4 |
| 3 | C | H5 | 2 | A | H2' | 2.5 | 3.99 |
| 3 | C | H6 | 2 | A | H1' | 2.86 | 4.56 |
| 3 | C | H6 | 3 | C | H1' | 2.67 | 4.26 |
| 3 | C | N3 | 24 | G | H1 | 1.8 | 2.4 |
| 3 | C | O2 | 24 | G | H22 | 1.8 | 2.4 |
| 4 | A | H1' | 3 | C | H2' | 3.14 | 5.01 |
| 4 | A | H1' | 4 | A | H2' | 2.1 | 3.35 |
| 4 | A | H1' | 4 | A | H3' | 2.44 | 3.89 |
| 4 | A | H1' | 4 | A | H4' | 2.55 | 4.07 |
| 4 | A | H2 | 4 | A | H1' | 3.75 | 5.99 |
| 4 | A | H2 | 5 | G | H1' | 2.53 | 4.03 |
| 4 | A | H2 | 23 | U | H1' | 4.01 | 6.4 |
| 4 | A | H2 | 24 | G | H1' | 2.53 | 4.03 |
| 4 | A | H61 | 23 | U | O4 | 1.8 | 2.4 |
| 4 | A | H8 | 3 | C | H1' | 3 | 4.79 |
| 4 | A | H8 | 4 | A | H1' | 2.91 | 4.64 |
| 4 | A | N1 | 23 | U | H3 | 1.8 | 2.4 |
| 5 | G | H1 | 22 | C | N3 | 1.8 | 2.4 |
| 5 | G | H1' | 4 | A | H2' | 2.82 | 4.5 |
| 5 | G | H1' | 5 | G | H2' | 2.05 | 3.28 |

| | | | | | | | |
|---|---|---|---|---|---|---|---|
| 5 | G | H22 | 22 | C | O2 | 1.8 | 2.4 |
| 5 | G | H8 | 4 | A | H1' | 2.97 | 4.74 |
| 5 | G | H8 | 5 | G | H1' | 2.84 | 4.52 |
| 5 | G | O6 | 22 | C | H41 | 1.8 | 2.4 |
| 6 | C | H1' | 5 | G | H2' | 3 | 6 |
| 6 | C | H1' | 6 | C | H2' | 2.01 | 3.21 |
| 6 | C | H1' | 6 | C | H3' | 2.42 | 3.85 |
| 6 | C | H41 | 21 | G | O6 | 1.8 | 2.4 |
| 6 | C | H5 | 5 | G | H2' | 2.63 | 4.2 |
| 6 | C | H6 | 5 | G | H1' | 3 | 6 |
| 6 | C | H6 | 6 | C | H1' | 2.62 | 4.18 |
| 6 | C | N3 | 21 | G | H1 | 1.8 | 2.4 |
| 6 | C | O2 | 21 | G | H22 | 1.8 | 2.4 |
| 7 | A | H1' | 6 | C | H2' | 3 | 6 |
| 7 | A | H1' | 7 | A | H2' | 2.06 | 3.29 |
| 7 | A | H1' | 7 | A | H3' | 2.67 | 4.26 |
| 7 | A | H2 | 7 | A | H1' | 3.47 | 5.53 |
| 7 | A | H2 | 8 | G | H1' | 1.8 | 4.5 |
| 7 | A | H8 | 6 | C | H1' | 3 | 6 |
| 7 | A | H8 | 6 | C | H6 | 3 | 6 |
| 7 | A | H8 | 7 | A | H1' | 2.63 | 4.2 |
| 7 | A | H8 | 8 | G | H8 | 3 | 6 |
| 8 | G | H1 | 19 | C | N3 | 1.8 | 2.4 |
| 8 | G | H1' | 8 | G | H2' | 1.86 | 2.96 |
| 8 | G | H22 | 19 | C | O2 | 1.8 | 2.4 |
| 8 | G | H8 | 7 | A | H1' | 3 | 6 |
| 8 | G | H8 | 8 | G | H1' | 2.73 | 4.35 |
| 8 | G | H8 | 9 | C | H5 | 3.11 | 4.97 |
| 8 | G | O6 | 19 | C | H41 | 1.8 | 2.4 |
| 9 | C | H1' | 9 | C | H2' | 1.92 | 3.06 |
| 9 | C | H41 | 18 | G | O6 | 1.8 | 2.4 |
| 9 | C | H5 | 8 | G | H2' | 3 | 6 |
| 9 | C | H5 | 9 | C | H2' | 3 | 6 |
| 9 | C | H6 | 9 | C | H1' | 2.36 | 3.77 |
| 9 | C | H6 | 10 | U | H5 | 3.31 | 5.28 |
| 9 | C | N3 | 18 | G | H1 | 1.8 | 2.4 |
| 9 | C | O2 | 18 | G | H22 | 1.8 | 2.4 |
| 10 | U | H1' | 9 | C | H2' | 3 | 6 |
| 10 | U | H1' | 10 | U | H2' | 2.11 | 3.36 |
| 10 | U | H1' | 10 | U | H3' | 3.03 | 4.84 |
| 10 | U | H1' | 10 | U | H4' | 2.32 | 3.69 |
| 10 | U | H3 | 17 | A | N1 | 1.8 | 2.4 |
| 10 | U | H6 | 9 | C | H1' | 3 | 6 |

| | | | | | | | |
|---|---|---|---|---|---|---|---|
| 10 | U | H6 | 10 | U | H1' | 2.6 | 4.14 |
| 10 | U | O4 | 17 | A | H61 | 1.8 | 2.4 |
| 11 | G | H1 | 16 | C | N3 | 1.8 | 2.4 |
| 11 | G | H1' | 10 | U | H2' | 3.18 | 5.07 |
| 11 | G | H1' | 11 | G | H2' | 2.03 | 3.24 |
| 11 | G | H1' | 11 | G | H3' | 2.66 | 4.25 |
| 11 | G | H22 | 16 | C | O2 | 1.8 | 2.4 |
| 11 | G | H8 | 10 | U | H1' | 3 | 6 |
| 11 | G | H8 | 11 | G | H1' | 2.73 | 4.36 |
| 11 | G | O6 | 16 | C | H41 | 1.8 | 2.4 |
| 12 | U | H1' | 12 | U | H2' | 1.99 | 3.18 |
| 12 | U | H3 | 15 | A | N1 | 1.8 | 2.4 |
| 12 | U | H5 | 11 | G | H2' | 2.55 | 4.07 |
| 12 | U | H5 | 12 | U | H3' | 2.92 | 4.65 |
| 12 | U | H6 | 11 | G | H1' | 3 | 6 |
| 12 | U | H6 | 12 | U | H1' | 2.55 | 4.07 |
| 12 | U | H6 | 13 | C | H5 | 2.91 | 4.64 |
| 12 | U | O4 | 15 | A | H61 | 1.8 | 2.4 |
| 13 | C | H1' | 12 | U | H2' | 2.93 | 4.67 |
| 13 | C | H1' | 13 | C | H2' | 2.01 | 3.21 |
| 13 | C | H41 | 14 | G | O6 | 1.8 | 2.4 |
| 13 | C | H5 | 12 | U | H2' | 2.52 | 4.02 |
| 13 | C | H5 | 12 | U | H3' | 2.59 | 4.13 |
| 13 | C | H6 | 12 | U | H1' | 3 | 6 |
| 13 | C | H6 | 13 | C | H1' | 2.5 | 3.99 |
| 13 | C | H6 | 13 | C | H2' | 2.35 | 3.74 |
| 13 | C | N3 | 14 | G | H1 | 1.8 | 2.4 |
| 13 | C | O2 | 14 | G | H22 | 1.8 | 2.4 |
| 14 | G | H1' | 14 | G | H2' | 2.42 | 3.86 |
| 14 | G | H1' | 14 | G | H3' | 2.84 | 4.52 |
| 14 | G | H1' | 14 | G | H4' | 2.56 | 4.08 |
| 14 | G | H8 | 14 | G | H1' | 2.57 | 4.1 |
| 15 | A | H1' | 14 | G | H2' | 3.18 | 5.06 |
| 15 | A | H1' | 15 | A | H2' | 2 | 3.19 |
| 15 | A | H2 | 13 | C | H1' | 2.67 | 4.26 |
| 15 | A | H2 | 15 | A | H1' | 3.8 | 6.06 |
| 15 | A | H2 | 16 | C | H1' | 2.73 | 4.35 |
| 15 | A | H8 | 14 | G | H1' | 3.11 | 4.96 |
| 15 | A | H8 | 14 | G | H2' | 2.45 | 3.91 |
| 15 | A | H8 | 15 | A | H1' | 3.07 | 4.89 |
| 16 | C | H1' | 15 | A | H2' | 2.59 | 4.13 |
| 16 | C | H1' | 16 | C | H2' | 2.01 | 3.2 |
| 16 | C | H1' | 16 | C | H4' | 2.74 | 4.37 |

| | | | | | | | |
|---|---|---|---|---|---|---|---|
| 16 | C | H5 | 15 | A | H2' | 2.5 | 3.99 |
| 16 | C | H6 | 15 | A | H1' | 2.86 | 4.56 |
| 16 | C | H6 | 16 | C | H1' | 2.67 | 4.26 |
| 17 | A | H1' | 16 | C | H2' | 3.14 | 5.01 |
| 17 | A | H1' | 17 | A | H2' | 2.1 | 3.35 |
| 17 | A | H1' | 17 | A | H3' | 2.44 | 3.89 |
| 17 | A | H1' | 17 | A | H4' | 2.55 | 4.07 |
| 17 | A | H2 | 10 | U | H1' | 4.01 | 6.4 |
| 17 | A | H2 | 11 | G | H1' | 2.53 | 4.03 |
| 17 | A | H2 | 17 | A | H1' | 3.75 | 5.99 |
| 17 | A | H2 | 18 | G | H1' | 2.53 | 4.03 |
| 17 | A | H8 | 16 | C | H1' | 3 | 4.79 |
| 17 | A | H8 | 17 | A | H1' | 2.91 | 4.64 |
| 18 | G | H1' | 17 | A | H2' | 2.82 | 4.5 |
| 18 | G | H1' | 18 | G | H2' | 2.05 | 3.28 |
| 18 | G | H8 | 17 | A | H1' | 2.97 | 4.74 |
| 18 | G | H8 | 18 | G | H1' | 2.84 | 4.52 |
| 19 | C | H1' | 18 | G | H2' | 3 | 6 |
| 19 | C | H1' | 19 | C | H2' | 2.01 | 3.21 |
| 19 | C | H1' | 19 | C | H3' | 2.42 | 3.85 |
| 19 | C | H5 | 18 | G | H2' | 3 | 6 |
| 19 | C | H6 | 18 | G | H1' | 3 | 6 |
| 19 | C | H6 | 19 | C | H1' | 2.62 | 4.18 |
| 20 | A | H1' | 19 | C | H2' | 3 | 6 |
| 20 | A | H1' | 20 | A | H2' | 2.06 | 3.29 |
| 20 | A | H1' | 20 | A | H3' | 2.67 | 4.26 |
| 20 | A | H2 | 20 | A | H1' | 3.47 | 5.53 |
| 20 | A | H2 | 21 | G | H1' | 1.8 | 4.5 |
| 20 | A | H8 | 19 | C | H1' | 3 | 6 |
| 20 | A | H8 | 19 | C | H6 | 3 | 6 |
| 20 | A | H8 | 20 | A | H1' | 2.63 | 4.2 |
| 20 | A | H8 | 21 | G | H8 | 3 | 6 |
| 21 | G | H1' | 21 | G | H2' | 1.86 | 2.96 |
| 21 | G | H8 | 20 | A | H1' | 3 | 6 |
| 21 | G | H8 | 21 | G | H1' | 2.73 | 4.35 |
| 21 | G | H8 | 22 | C | H5 | 3.11 | 4.97 |
| 22 | C | H1' | 22 | C | H2' | 1.92 | 3.06 |
| 22 | C | H5 | 21 | G | H2' | 3 | 6 |
| 22 | C | H5 | 22 | C | H2' | 3 | 6 |
| 22 | C | H6 | 22 | C | H1' | 2.36 | 3.77 |
| 22 | C | H6 | 23 | U | H5 | 3.31 | 5.28 |
| 23 | U | H1' | 22 | C | H2' | 3 | 6 |
| 23 | U | H1' | 23 | U | H2' | 2.11 | 3.36 |

| | | | | | | | |
|---|---|---|---|---|---|---|---|
| 23 | U | H1' | 23 | U | H3' | 3.03 | 4.84 |
| 23 | U | H1' | 23 | U | H4' | 2.32 | 3.69 |
| 23 | U | H6 | 22 | C | H1' | 3 | 6 |
| 23 | U | H6 | 23 | U | H1' | 2.6 | 4.14 |
| 24 | G | H1' | 23 | U | H2' | 3.18 | 5.07 |
| 24 | G | H1' | 24 | G | H2' | 2.03 | 3.24 |
| 24 | G | H1' | 24 | G | H3' | 2.66 | 4.25 |
| 24 | G | H8 | 23 | U | H1' | 3 | 6 |
| 24 | G | H8 | 24 | G | H1' | 2.73 | 4.36 |
| 25 | U | H1' | 25 | U | H2' | 1.99 | 3.18 |
| 25 | U | H5 | 24 | G | H2' | 2.55 | 4.07 |
| 25 | U | H5 | 25 | U | H3' | 2.92 | 4.65 |
| 25 | U | H6 | 24 | G | H1' | 3 | 6 |
| 25 | U | H6 | 25 | U | H1' | 2.55 | 4.07 |
| 25 | U | H6 | 26 | C | H5 | 2.91 | 4.64 |
| 26 | C | H1' | 25 | U | H2' | 2.93 | 4.67 |
| 26 | C | H1' | 26 | C | H2' | 2.01 | 3.21 |
| 26 | C | H5 | 25 | U | H2' | 2.52 | 4.02 |
| 26 | C | H5 | 25 | U | H3' | 2.59 | 4.13 |
| 26 | C | H6 | 25 | U | H1' | 3 | 6 |
| 26 | C | H6 | 26 | C | H1' | 2.5 | 3.99 |
| 26 | C | H6 | 26 | C | H2' | 2.35 | 3.74 |
| 27 | D6D | H1 | 7 | A | H1' | 3 | 6 |
| 27 | D6D | H1 | 7 | A | H2' | 2 | 4 |
| 27 | D6D | H1 | 7 | A | H3' | 3 | 6 |
| 27 | D6D | H1 | 8 | G | H1' | 3 | 6 |
| 27 | D6D | H3 | 7 | A | H2 | 3 | 6 |
| 27 | D6D | H3 | 8 | G | H1' | 3 | 6 |
| 27 | D6D | H5 | 20 | A | H2 | 3 | 6 |
| 27 | D6D | H5 | 20 | A | H8 | 3 | 6 |
| 27 | D6D | H5 | 21 | G | H1' | 2 | 4 |
| 27 | D6D | H7 | 20 | A | H1' | 3 | 6 |
| 27 | D6D | H7 | 20 | A | H2 | 3 | 6 |
| 27 | D6D | H7 | 20 | A | H8 | 3 | 6 |
| 27 | D6D | H7 | 21 | G | H1' | 3 | 6 |

**Table S8:** NOE restraints used for modeling of the r(CAG)-**2** complex.

| | | | | | | | |
|---|---|---|---|---|---|---|---|
| 1 | G | H1 | 26 | C | N3 | 1.8 | 2.4 |
| 1 | G | H1' | 1 | G | H3' | 3.24 | 5.17 |
| 1 | G | H22 | 26 | C | O2 | 1.8 | 2.4 |
| 1 | G | H8 | 1 | G | H1' | 2.55 | 4.07 |
| 1 | G | H8 | 1 | G | H3' | 2.24 | 3.58 |
| 1 | G | O6 | 26 | C | H41 | 1.8 | 2.4 |
| 2 | A | H1' | 2 | A | H2' | 2.14 | 3.41 |
| 2 | A | H1' | 2 | A | H4' | 2.63 | 4.2 |
| 2 | A | H2 | 2 | A | H1' | 4.3 | 6.86 |
| 2 | A | H2 | 3 | C | H1' | 2.72 | 4.33 |
| 2 | A | H2 | 26 | C | H1' | 2.58 | 4.11 |
| 2 | A | H61 | 25 | U | O4 | 1.8 | 2.4 |
| 2 | A | H8 | 1 | G | H3' | 3.18 | 5.08 |
| 2 | A | H8 | 2 | A | H1' | 2.86 | 4.56 |
| 2 | A | N1 | 25 | U | H3 | 1.8 | 2.4 |
| 3 | C | H41 | 24 | G | O6 | 1.8 | 2.4 |
| 3 | C | H5 | 2 | A | H2' | 3.31 | 5.28 |
| 3 | C | H6 | 2 | A | H2' | 1.67 | 2.66 |
| 3 | C | H6 | 3 | C | H1' | 2.79 | 4.45 |
| 3 | C | H6 | 3 | C | H2' | 2.62 | 4.18 |
| 3 | C | N3 | 24 | G | H1 | 1.8 | 2.4 |
| 3 | C | O2 | 24 | G | H22 | 1.8 | 2.4 |
| 4 | A | H1' | 3 | C | H2' | 3.55 | 5.66 |
| 4 | A | H1' | 4 | A | H2' | 2.09 | 3.33 |
| 4 | A | H1' | 4 | A | H3' | 2.88 | 4.6 |
| 4 | A | H1' | 4 | A | H4' | 2.51 | 4.01 |
| 4 | A | H2 | 4 | A | H1' | 3.44 | 5.48 |
| 4 | A | H2 | 5 | G | H1' | 2.44 | 3.89 |
| 4 | A | H2 | 24 | G | H1' | 2.46 | 3.92 |
| 4 | A | H61 | 23 | U | O4 | 1.8 | 2.4 |
| 4 | A | H8 | 3 | C | H2' | 1.83 | 2.91 |
| 4 | A | H8 | 4 | A | H1' | 2.86 | 4.56 |
| 4 | A | H8 | 4 | A | H2' | 2.83 | 4.51 |
| 4 | A | H8 | 4 | A | H3' | 2.15 | 3.43 |
| 4 | A | N1 | 23 | U | H3 | 1.8 | 2.4 |
| 5 | G | H1 | 22 | C | N3 | 1.8 | 2.4 |
| 5 | G | H1' | 4 | A | H2' | 3.4 | 5.42 |
| 5 | G | H1' | 5 | G | H2' | 2.01 | 3.2 |
| 5 | G | H1' | 5 | G | H3' | 1.8 | 4.5 |
| 5 | G | H22 | 22 | C | O2 | 1.8 | 2.4 |

| | | | | | | | |
|---|---|---|---|---|---|---|---|
| 5 | G | H8 | 4 | A | H2' | 1.82 | 2.91 |
| 5 | G | H8 | 4 | A | H3' | 2.5 | 3.99 |
| 5 | G | H8 | 5 | G | H1' | 2.88 | 4.59 |
| 5 | G | H8 | 5 | G | H2' | 3.14 | 5.02 |
| 5 | G | H8 | 5 | G | H3' | 2.05 | 3.26 |
| 5 | G | H8 | 6 | C | H5 | 3.24 | 5.17 |
| 5 | G | O6 | 22 | C | H41 | 1.8 | 2.4 |
| 6 | C | H1' | 6 | C | H2' | 2 | 3.19 |
| 6 | C | H41 | 21 | G | O6 | 1.8 | 2.4 |
| 6 | C | H5 | 5 | G | H2' | 3.33 | 5.32 |
| 6 | C | H6 | 5 | G | H2' | 1.85 | 2.95 |
| 6 | C | H6 | 5 | G | H3' | 2.31 | 3.68 |
| 6 | C | H6 | 6 | C | H1' | 2.95 | 4.71 |
| 6 | C | H6 | 6 | C | H2' | 1.8 | 4.5 |
| 6 | C | N3 | 21 | G | H1 | 1.8 | 2.4 |
| 6 | C | O2 | 21 | G | H22 | 1.8 | 2.4 |
| 7 | A | H1' | 6 | C | H2' | 3.24 | 5.17 |
| 7 | A | H1' | 7 | A | H2' | 2.12 | 3.39 |
| 7 | A | H1' | 7 | A | H3' | 3.02 | 4.83 |
| 7 | A | H2 | 8 | G | H1' | 1.8 | 4.5 |
| 7 | A | H2 | 21 | G | H1' | 1.8 | 4.5 |
| 7 | A | H2 | 27 | D6L | H3 | 3 | 6 |
| 7 | A | H8 | 6 | C | H2' | 2.09 | 3.34 |
| 7 | A | H8 | 7 | A | H1' | 2.85 | 4.55 |
| 7 | A | H8 | 7 | A | H2' | 1.8 | 4.5 |
| 7 | A | H8 | 7 | A | H3' | 2.17 | 3.47 |
| 8 | G | H1 | 19 | C | N3 | 1.8 | 2.4 |
| 8 | G | H22 | 19 | C | O2 | 1.8 | 2.4 |
| 8 | G | H8 | 7 | A | H2' | 2.25 | 3.59 |
| 8 | G | H8 | 7 | A | H3' | 2.93 | 4.68 |
| 8 | G | H8 | 8 | G | H2' | 1.8 | 4.5 |
| 8 | G | H8 | 27 | D6L | H3 | 3 | 6 |
| 8 | G | O6 | 19 | C | H41 | 1.8 | 2.4 |
| 9 | C | H41 | 18 | G | O6 | 1.8 | 2.4 |
| 9 | C | H5 | 8 | G | H2' | 2.78 | 4.44 |
| 9 | C | H6 | 8 | G | H2' | 1.74 | 2.77 |
| 9 | C | H6 | 10 | U | H5 | 2.72 | 4.35 |
| 9 | C | N3 | 18 | G | H1 | 1.8 | 2.4 |
| 9 | C | O2 | 18 | G | H22 | 1.8 | 2.4 |
| 10 | U | H1' | 10 | U | H2' | 2.1 | 3.35 |
| 10 | U | H3 | 17 | A | N1 | 1.8 | 2.4 |
| 10 | U | H5 | 9 | C | H2' | 2.4 | 3.83 |
| 10 | U | H5 | 10 | U | H3' | 3.28 | 5.23 |

| | | | | | | | |
|---|---|---|---|---|---|---|---|
| 10 | U | H6 | 9 | C | H2' | 1.75 | 2.8 |
| 10 | U | O4 | 17 | A | H61 | 1.8 | 2.4 |
| 11 | G | H1 | 16 | C | N3 | 1.8 | 2.4 |
| 11 | G | H1' | 10 | U | H2' | 2.88 | 4.59 |
| 11 | G | H1' | 11 | G | H2' | 1.94 | 3.1 |
| 11 | G | H1' | 11 | G | H3' | 2.5 | 3.98 |
| 11 | G | H22 | 16 | C | O2 | 1.8 | 2.4 |
| 11 | G | H8 | 10 | U | H2' | 1.93 | 3.07 |
| 11 | G | H8 | 11 | G | H1' | 2.92 | 4.66 |
| 11 | G | H8 | 11 | G | H2' | 1.8 | 4.5 |
| 11 | G | H8 | 12 | U | H5 | 2.87 | 4.57 |
| 11 | G | O6 | 16 | C | H41 | 1.8 | 2.4 |
| 12 | U | H1' | 12 | U | H2' | 1.93 | 3.07 |
| 12 | U | H3 | 15 | A | N1 | 1.8 | 2.4 |
| 12 | U | H5 | 11 | G | H3' | 3.21 | 5.12 |
| 12 | U | H6 | 11 | G | H2' | 1.71 | 2.72 |
| 12 | U | H6 | 11 | G | H3' | 1.99 | 3.17 |
| 12 | U | H6 | 12 | U | H1' | 2.46 | 3.92 |
| 12 | U | H6 | 12 | U | H2' | 2.35 | 3.74 |
| 12 | U | H6 | 12 | U | H3' | 1.9 | 3.02 |
| 12 | U | O4 | 15 | A | H61 | 1.8 | 2.4 |
| 13 | C | H1' | 12 | U | H2' | 2.84 | 4.52 |
| 13 | C | H1' | 13 | C | H3' | 2.5 | 4 |
| 13 | C | H41 | 14 | G | O6 | 1.8 | 2.4 |
| 13 | C | H5 | 12 | U | H2' | 2.67 | 4.26 |
| 13 | C | H5 | 12 | U | H3' | 2.56 | 4.08 |
| 13 | C | H5 | 13 | C | H3' | 2.76 | 4.4 |
| 13 | C | H6 | 12 | U | H2' | 1.78 | 2.83 |
| 13 | C | H6 | 12 | U | H3' | 1.98 | 3.16 |
| 13 | C | H6 | 13 | C | H1' | 2.39 | 3.8 |
| 13 | C | H6 | 13 | C | H2' | 2.21 | 3.53 |
| 13 | C | H6 | 13 | C | H3' | 1.76 | 2.81 |
| 13 | C | N3 | 14 | G | H1 | 1.8 | 2.4 |
| 13 | C | O2 | 14 | G | H22 | 1.8 | 2.4 |
| 14 | G | H1' | 14 | G | H3' | 3.24 | 5.17 |
| 14 | G | H8 | 14 | G | H1' | 2.55 | 4.07 |
| 14 | G | H8 | 14 | G | H3' | 2.24 | 3.58 |
| 15 | A | H1' | 15 | A | H2' | 2.14 | 3.41 |
| 15 | A | H1' | 15 | A | H4' | 2.63 | 4.2 |
| 15 | A | H2 | 13 | C | H1' | 2.58 | 4.11 |
| 15 | A | H2 | 15 | A | H1' | 4.3 | 6.86 |
| 15 | A | H2 | 16 | C | H1' | 2.72 | 4.33 |
| 15 | A | H8 | 14 | G | H3' | 3.18 | 5.08 |

| | | | | | | | |
|---|---|---|---|---|---|---|---|
| 15 | A | H8 | 15 | A | H1' | 2.86 | 4.56 |
| 16 | C | H5 | 15 | A | H2' | 3.31 | 5.28 |
| 16 | C | H6 | 15 | A | H2' | 1.67 | 2.66 |
| 16 | C | H6 | 16 | C | H1' | 2.79 | 4.45 |
| 16 | C | H6 | 16 | C | H2' | 2.62 | 4.18 |
| 17 | A | H1' | 16 | C | H2' | 3.55 | 5.66 |
| 17 | A | H1' | 17 | A | H2' | 2.09 | 3.33 |
| 17 | A | H1' | 17 | A | H3' | 2.88 | 4.6 |
| 17 | A | H1' | 17 | A | H4' | 2.51 | 4.01 |
| 17 | A | H2 | 11 | G | H1' | 2.46 | 3.92 |
| 17 | A | H2 | 17 | A | H1' | 3.44 | 5.48 |
| 17 | A | H2 | 18 | G | H1' | 2.44 | 3.89 |
| 17 | A | H8 | 16 | C | H2' | 1.83 | 2.91 |
| 17 | A | H8 | 17 | A | H1' | 2.86 | 4.56 |
| 17 | A | H8 | 17 | A | H2' | 2.83 | 4.51 |
| 17 | A | H8 | 17 | A | H3' | 2.15 | 3.43 |
| 18 | G | H1' | 17 | A | H2' | 3.4 | 5.42 |
| 18 | G | H1' | 18 | G | H2' | 2.01 | 3.2 |
| 18 | G | H1' | 18 | G | H3' | 1.8 | 4.5 |
| 18 | G | H8 | 17 | A | H2' | 1.82 | 2.91 |
| 18 | G | H8 | 17 | A | H3' | 2.5 | 3.99 |
| 18 | G | H8 | 18 | G | H1' | 2.88 | 4.59 |
| 18 | G | H8 | 18 | G | H2' | 3.14 | 5.02 |
| 18 | G | H8 | 18 | G | H3' | 2.05 | 3.26 |
| 18 | G | H8 | 19 | C | H5 | 3.24 | 5.17 |
| 19 | C | H1' | 19 | C | H2' | 2 | 3.19 |
| 19 | C | H5 | 18 | G | H2' | 3.33 | 5.32 |
| 19 | C | H6 | 18 | G | H2' | 1.85 | 2.95 |
| 19 | C | H6 | 18 | G | H3' | 2.31 | 3.68 |
| 19 | C | H6 | 19 | C | H1' | 2.95 | 4.71 |
| 19 | C | H6 | 19 | C | H2' | 1.8 | 4.5 |
| 19 | C | H6 | 27 | D6L | H7 | 3.5 | 6 |
| 20 | A | H1' | 19 | C | H2' | 3.24 | 5.17 |
| 20 | A | H1' | 20 | A | H2' | 2.12 | 3.39 |
| 20 | A | H1' | 20 | A | H3' | 3.02 | 4.83 |
| 20 | A | H2 | 8 | G | H1' | 1.8 | 4.5 |
| 20 | A | H2 | 21 | G | H1' | 1.8 | 4.5 |
| 20 | A | H8 | 19 | C | H2' | 2.09 | 3.34 |
| 20 | A | H8 | 20 | A | H1' | 2.85 | 4.55 |
| 20 | A | H8 | 20 | A | H2' | 2.8 | 4.46 |
| 20 | A | H8 | 20 | A | H3' | 2.17 | 3.47 |
| 21 | G | H8 | 20 | A | H2' | 2.25 | 3.59 |
| 21 | G | H8 | 20 | A | H3' | 2.93 | 4.68 |

| | | | | | | | |
|---|---|---|---|---|---|---|---|
| 21 | G | H8 | 21 | G | H2' | 1.8 | 4.5 |
| 22 | C | H5 | 21 | G | H2' | 2.78 | 4.44 |
| 22 | C | H6 | 21 | G | H2' | 1.74 | 2.77 |
| 22 | C | H6 | 23 | U | H5 | 2.72 | 4.35 |
| 23 | U | H1' | 23 | U | H2' | 2.1 | 3.35 |
| 23 | U | H5 | 22 | C | H2' | 2.4 | 3.83 |
| 23 | U | H5 | 23 | U | H3' | 3.28 | 5.23 |
| 23 | U | H6 | 22 | C | H2' | 1.75 | 2.8 |
| 24 | G | H1' | 23 | U | H2' | 2.88 | 4.59 |
| 24 | G | H1' | 24 | G | H2' | 1.94 | 3.1 |
| 24 | G | H1' | 24 | G | H3' | 2.5 | 3.98 |
| 24 | G | H8 | 23 | U | H2' | 1.93 | 3.07 |
| 24 | G | H8 | 24 | G | H1' | 2.92 | 4.66 |
| 24 | G | H8 | 24 | G | H2' | 1.8 | 4.5 |
| 24 | G | H8 | 25 | U | H5 | 2.87 | 4.57 |
| 25 | U | H1' | 25 | U | H2' | 1.93 | 3.07 |
| 25 | U | H5 | 24 | G | H3' | 3.21 | 5.12 |
| 25 | U | H6 | 24 | G | H2' | 1.71 | 2.72 |
| 25 | U | H6 | 24 | G | H3' | 1.99 | 3.17 |
| 25 | U | H6 | 25 | U | H1' | 2.46 | 3.92 |
| 25 | U | H6 | 25 | U | H2' | 2.35 | 3.74 |
| 25 | U | H6 | 25 | U | H3' | 1.9 | 3.02 |
| 26 | C | H1' | 25 | U | H2' | 2.84 | 4.52 |
| 26 | C | H1' | 26 | C | H3' | 2.5 | 4 |
| 26 | C | H5 | 25 | U | H2' | 2.67 | 4.26 |
| 26 | C | H5 | 25 | U | H3' | 2.56 | 4.08 |
| 26 | C | H5 | 26 | C | H3' | 2.76 | 4.4 |
| 26 | C | H6 | 25 | U | H2' | 1.78 | 2.83 |
| 26 | C | H6 | 25 | U | H3' | 1.98 | 3.16 |
| 26 | C | H6 | 26 | C | H1' | 2.39 | 3.8 |
| 26 | C | H6 | 26 | C | H2' | 2.21 | 3.53 |
| 26 | C | H6 | 26 | C | H3' | 1.76 | 2.81 |
| 27 | D6L | H1 | 7 | A | H2 | 3 | 6 |
| 27 | D6L | H3 | 6 | C | H2' | 3 | 6.2 |
| 27 | D6L | H3 | 7 | A | H2' | 3 | 6 |
| 27 | D6L | H5 | 19 | C | H2' | 3 | 6.2 |
| 27 | D6L | H5 | 19 | C | H5 | 3 | 6 |
| 27 | D6L | H5 | 19 | C | H6 | 3.5 | 6 |
| 27 | D6L | H5 | 20 | A | H2' | 3 | 6 |
| 27 | D6L | H5 | 20 | A | H8 | 3 | 6 |
| 27 | D6L | H7 | 20 | A | H8 | 3 | 6 |

**Table S9**: NOE restraints used for modeling of the r(CAG)-**3** complex.

| | | | | | | | |
|---|---|---|---|---|---|---|---|
| 1 | G | H1 | 26 | C | N3 | 1.8 | 2.4 |
| 1 | G | H1' | 1 | G | H2' | 2.04 | 3.25 |
| 1 | G | H1' | 1 | G | H3' | 3 | 6 |
| 1 | G | H22 | 26 | C | O2 | 1.8 | 2.4 |
| 1 | G | H8 | 1 | G | H1' | 2.41 | 3.84 |
| 1 | G | H8 | 1 | G | H2' | 2.48 | 3.96 |
| 1 | G | H8 | 1 | G | H2' | 2.48 | 3.96 |
| 1 | G | H8 | 1 | G | H3' | 2.7 | 4.31 |
| 1 | G | O6 | 26 | C | H41 | 1.8 | 2.4 |
| 2 | A | H1' | 2 | A | H2' | 2.17 | 3.45 |
| 2 | A | H1' | 2 | A | H3' | 3 | 6 |
| 2 | A | H1' | 2 | A | H4' | 2.48 | 3.96 |
| 2 | A | H2 | 2 | A | H1' | 3 | 6 |
| 2 | A | H2 | 3 | C | H1' | 2.54 | 4.04 |
| 2 | A | H2 | 3 | C | H5 | 3 | 6 |
| 2 | A | H2 | 26 | C | H1' | 2.58 | 4.11 |
| 2 | A | H61 | 25 | U | O4 | 1.8 | 2.4 |
| 2 | A | H8 | 1 | G | H2' | 1.8 | 4.5 |
| 2 | A | H8 | 2 | A | H1' | 2.94 | 4.69 |
| 2 | A | H8 | 2 | A | H2' | 1.8 | 4.5 |
| 2 | A | H8 | 2 | A | H3' | 1.8 | 4.5 |
| 2 | A | H8 | 3 | C | H5 | 3 | 6 |
| 2 | A | N1 | 25 | U | H3 | 1.8 | 2.4 |
| 3 | C | H1' | 3 | C | H2' | 1.8 | 4.5 |
| 3 | C | H1' | 3 | C | H3' | 2.7 | 4.31 |
| 3 | C | H1' | 3 | C | H4' | 2.56 | 4.08 |
| 3 | C | H41 | 24 | G | O6 | 1.8 | 2.4 |
| 3 | C | H5 | 2 | A | H2' | 3 | 6 |
| 3 | C | H5 | 3 | C | H2' | 3 | 6 |
| 3 | C | H5 | 3 | C | H3' | 3 | 6 |
| 3 | C | H6 | 2 | A | H2' | 1.88 | 3 |
| 3 | C | H6 | 2 | A | H3' | 2.51 | 4.01 |
| 3 | C | H6 | 3 | C | H1' | 2.79 | 4.45 |
| 3 | C | H6 | 3 | C | H2' | 2.6 | 4.15 |
| 3 | C | H6 | 3 | C | H3' | 1.91 | 3.05 |
| 3 | C | N3 | 24 | G | H1 | 1.8 | 2.4 |
| 3 | C | O2 | 24 | G | H22 | 1.8 | 2.4 |
| 4 | A | H1' | 3 | C | H2' | 3 | 6 |
| 4 | A | H1' | 4 | A | H2' | 2.42 | 3.86 |
| 4 | A | H1' | 4 | A | H4' | 3.19 | 5.09 |
| 4 | A | H2 | 4 | A | H1' | 3 | 6 |
| 4 | A | H2 | 5 | G | H1' | 2.39 | 3.8 |
| 4 | A | H2 | 24 | G | H1' | 2.93 | 4.67 |
| 4 | A | H61 | 23 | U | O4 | 1.8 | 2.4 |
| 4 | A | H8 | 3 | C | H2' | 1.83 | 2.92 |
| 4 | A | H8 | 3 | C | H3' | 2.39 | 3.8 |
| 4 | A | H8 | 4 | A | H1' | 3.21 | 5.12 |
| 4 | A | H8 | 4 | A | H2' | 1.8 | 4.5 |
| 4 | A | N1 | 23 | U | H3 | 1.8 | 2.4 |

| | | | | | | | |
|---|---|---|---|---|---|---|---|
| 5 | G | H1 | 22 | C | N3 | 1.8 | 2.4 |
| 5 | G | H1' | 4 | A | H2' | 3 | 6 |
| 5 | G | H1' | 5 | G | H2' | 1.93 | 3.07 |
| 5 | G | H1' | 5 | G | H4' | 3 | 6 |
| 5 | G | H22 | 22 | C | O2 | 1.8 | 2.4 |
| 5 | G | H8 | 4 | A | H2' | 2.09 | 3.34 |
| 5 | G | H8 | 5 | G | H1' | 1.8 | 4.5 |
| 5 | G | H8 | 5 | G | H2' | 1.8 | 4.5 |
| 5 | G | H8 | 5 | G | H3' | 2.32 | 3.69 |
| 5 | G | H8 | 6 | C | H5 | 3 | 6 |
| 5 | G | O6 | 22 | C | H41 | 1.8 | 2.4 |
| 6 | C | H1' | 6 | C | H2' | 2.02 | 3.23 |
| 6 | C | H41 | 21 | G | O6 | 1.8 | 2.4 |
| 6 | C | H5 | 5 | G | H2' | 3.09 | 4.93 |
| 6 | C | H5 | 6 | C | H2' | 3 | 6 |
| 6 | C | H6 | 5 | G | H2' | 1.91 | 3.05 |
| 6 | C | H6 | 5 | G | H3' | 2.15 | 3.43 |
| 6 | C | H6 | 6 | C | H1' | 2.78 | 4.43 |
| 6 | C | H6 | 6 | C | H2' | 2.37 | 3.78 |
| 6 | C | N3 | 21 | G | H1 | 1.8 | 2.4 |
| 6 | C | O2 | 21 | G | H22 | 1.8 | 2.4 |
| 7 | A | H1' | 6 | C | H2' | 3 | 6 |
| 7 | A | H1' | 7 | A | H4' | 2.66 | 4.25 |
| 7 | A | H8 | 6 | C | H2' | 3 | 6 |
| 7 | A | H8 | 6 | C | H6 | 3 | 7 |
| 7 | A | H8 | 7 | A | H1' | 1.8 | 4.5 |
| 8 | G | H1 | 19 | C | N3 | 1.8 | 2.4 |
| 8 | G | H1' | 8 | G | H2' | 1.8 | 4.5 |
| 8 | G | H1' | 8 | G | H3' | 3 | 6 |
| 8 | G | H22 | 19 | C | O2 | 1.8 | 2.4 |
| 8 | G | H8 | 7 | A | H1' | 2.5 | 6 |
| 8 | G | H8 | 8 | G | H1' | 1.8 | 4.5 |
| 8 | G | H8 | 8 | G | H2' | 2.66 | 4.25 |
| 8 | G | H8 | 8 | G | H3' | 2.7 | 4.31 |
| 8 | G | H8 | 9 | C | H5 | 3 | 6 |
| 8 | G | O6 | 19 | C | H41 | 1.8 | 2.4 |
| 9 | C | H1' | 9 | C | H2' | 1.95 | 3.11 |
| 9 | C | H41 | 18 | G | O6 | 1.8 | 2.4 |
| 9 | C | H5 | 8 | G | H2' | 3 | 6 |
| 9 | C | H5 | 8 | G | H3' | 3.18 | 5.23 |
| 9 | C | H6 | 9 | C | H2' | 2.42 | 3.86 |
| 9 | C | H6 | 10 | U | H5 | 3 | 6 |
| 9 | C | N3 | 18 | G | H1 | 1.8 | 2.4 |
| 9 | C | O2 | 18 | G | H22 | 1.8 | 2.4 |
| 10 | U | H1' | 10 | U | H3' | 2.82 | 4.5 |
| 10 | U | H3 | 17 | A | N1 | 1.8 | 2.4 |
| 10 | U | H5 | 9 | C | H2' | 2.65 | 4.23 |
| 10 | U | H5 | 9 | C | H3' | 3 | 6 |
| 10 | U | H5 | 10 | U | H3' | 3 | 6 |
| 10 | U | H6 | 9 | C | H2' | 1.88 | 3 |
| 10 | U | H6 | 9 | C | H3' | 2.33 | 3.72 |

| | | | | | | | |
|---|---|---|---|---|---|---|---|
| 10 | U | H6 | 9 | C | H5 | 3 | 6 |
| 10 | U | H6 | 10 | U | H1' | 2.97 | 4.74 |
| 10 | U | H6 | 10 | U | H3' | 2.3 | 3.67 |
| 10 | U | O4 | 17 | A | H61 | 1.8 | 2.4 |
| 11 | G | H1 | 16 | C | N3 | 1.8 | 2.4 |
| 11 | G | H1' | 11 | G | H2' | 2.05 | 3.27 |
| 11 | G | H1' | 11 | G | H3' | 2.94 | 4.69 |
| 11 | G | H1' | 11 | G | H4' | 2.77 | 4.41 |
| 11 | G | H22 | 16 | C | O2 | 1.8 | 2.4 |
| 11 | G | H8 | 10 | U | H3' | 2.47 | 3.94 |
| 11 | G | H8 | 10 | U | H5 | 3 | 6 |
| 11 | G | H8 | 11 | G | H1' | 1.8 | 4.5 |
| 11 | G | H8 | 11 | G | H3' | 2.46 | 3.93 |
| 11 | G | O6 | 16 | C | H41 | 1.8 | 2.4 |
| 12 | U | H1' | 12 | U | H2' | 1.9 | 3.02 |
| 12 | U | H1' | 12 | U | H4' | 2.51 | 4.01 |
| 12 | U | H3 | 15 | A | N1 | 1.8 | 2.4 |
| 12 | U | H5 | 11 | G | H2' | 2.73 | 4.35 |
| 12 | U | H5 | 11 | G | H3' | 2.96 | 4.73 |
| 12 | U | H5 | 12 | U | H2' | 3 | 6 |
| 12 | U | H5 | 12 | U | H3' | 3 | 6 |
| 12 | U | H6 | 11 | G | H2' | 1.75 | 2.8 |
| 12 | U | H6 | 11 | G | H3' | 2.29 | 3.65 |
| 12 | U | H6 | 12 | U | H1' | 1.8 | 4.5 |
| 12 | U | H6 | 12 | U | H2' | 2.64 | 4.21 |
| 12 | U | H6 | 12 | U | H3' | 1.8 | 4.5 |
| 12 | U | H6 | 13 | C | H5 | 3 | 6 |
| 12 | U | O4 | 15 | A | H61 | 1.8 | 2.4 |
| 13 | C | H1' | 12 | U | H2' | 3 | 6 |
| 13 | C | H1' | 13 | C | H2' | 1.83 | 2.92 |
| 13 | C | H1' | 13 | C | H3' | 3 | 6 |
| 13 | C | H41 | 14 | G | O6 | 1.8 | 2.4 |
| 13 | C | H5 | 12 | U | H2' | 3.19 | 5.09 |
| 13 | C | H5 | 12 | U | H3' | 2.54 | 4.04 |
| 13 | C | H6 | 12 | U | H2' | 1.63 | 2.6 |
| 13 | C | H6 | 13 | C | H1' | 2.86 | 4.56 |
| 13 | C | H6 | 13 | C | H2' | 2.29 | 3.65 |
| 13 | C | H6 | 13 | C | H3' | 1.65 | 2.63 |
| 13 | C | N3 | 14 | G | H1 | 1.8 | 2.4 |
| 13 | C | O2 | 14 | G | H22 | 1.8 | 2.4 |
| 14 | G | H1' | 14 | G | H2' | 2.04 | 3.25 |
| 14 | G | H1' | 14 | G | H3' | 3 | 6 |
| 14 | G | H8 | 14 | G | H2' | 2.48 | 3.96 |
| 14 | G | H8 | 14 | G | H3' | 2.7 | 4.31 |
| 15 | A | H1' | 15 | A | H2' | 2.17 | 3.45 |
| 15 | A | H1' | 15 | A | H3' | 3 | 6 |
| 15 | A | H1' | 15 | A | H4' | 2.48 | 3.96 |
| 15 | A | H2 | 13 | C | H1' | 2.58 | 4.11 |
| 15 | A | H2 | 15 | A | H1' | 3 | 6 |
| 15 | A | H2 | 16 | C | H1' | 2.54 | 4.04 |
| 15 | A | H2 | 16 | C | H5 | 3 | 6 |

| | | | | | | | |
|---|---|---|---|---|---|---|---|
| 15 | A | H8 | 14 | G | H2' | 1.8 | 4.5 |
| 15 | A | H8 | 15 | A | H1' | 2.94 | 4.69 |
| 15 | A | H8 | 15 | A | H2' | 1.8 | 4.5 |
| 15 | A | H8 | 15 | A | H3' | 1.8 | 4.5 |
| 15 | A | H8 | 16 | C | H5 | 3 | 6 |
| 16 | C | H1' | 16 | C | H2' | 1.8 | 4.5 |
| 16 | C | H1' | 16 | C | H3' | 2.7 | 4.31 |
| 16 | C | H1' | 16 | C | H4' | 2.56 | 4.08 |
| 16 | C | H5 | 15 | A | H2' | 3 | 6 |
| 16 | C | H5 | 16 | C | H2' | 3 | 6 |
| 16 | C | H5 | 16 | C | H3' | 3 | 6 |
| 16 | C | H6 | 15 | A | H2' | 1.88 | 3 |
| 16 | C | H6 | 15 | A | H3' | 2.51 | 4.01 |
| 16 | C | H6 | 16 | C | H1' | 2.79 | 4.45 |
| 16 | C | H6 | 16 | C | H2' | 2.6 | 4.15 |
| 16 | C | H6 | 16 | C | H3' | 1.91 | 3.05 |
| 17 | A | H1' | 16 | C | H2' | 3 | 6 |
| 17 | A | H1' | 17 | A | H2' | 2.42 | 3.86 |
| 17 | A | H1' | 17 | A | H4' | 3.19 | 5.09 |
| 17 | A | H2 | 11 | G | H1' | 2.93 | 4.67 |
| 17 | A | H2 | 17 | A | H1' | 3 | 6 |
| 17 | A | H2 | 18 | G | H1' | 2.39 | 3.8 |
| 17 | A | H8 | 16 | C | H2' | 1.83 | 2.92 |
| 17 | A | H8 | 16 | C | H3' | 2.39 | 3.8 |
| 17 | A | H8 | 17 | A | H1' | 3.21 | 5.12 |
| 17 | A | H8 | 17 | A | H2' | 1.8 | 4.5 |
| 18 | G | H1' | 17 | A | H2' | 3 | 6 |
| 18 | G | H1' | 18 | G | H2' | 1.93 | 3.07 |
| 18 | G | H1' | 18 | G | H4' | 3 | 6 |
| 18 | G | H8 | 17 | A | H2' | 2.09 | 3.34 |
| 18 | G | H8 | 18 | G | H1' | 1.8 | 4.5 |
| 18 | G | H8 | 18 | G | H2' | 1.8 | 4.5 |
| 18 | G | H8 | 18 | G | H3' | 2.32 | 3.69 |
| 18 | G | H8 | 19 | C | H5 | 3 | 6 |
| 19 | C | H1' | 19 | C | H2' | 2.02 | 3.23 |
| 19 | C | H5 | 18 | G | H2' | 3.09 | 4.93 |
| 19 | C | H5 | 19 | C | H2' | 3 | 6 |
| 19 | C | H6 | 18 | G | H2' | 1.91 | 3.05 |
| 19 | C | H6 | 18 | G | H3' | 2.15 | 3.43 |
| 19 | C | H6 | 19 | C | H1' | 2.78 | 4.43 |
| 19 | C | H6 | 19 | C | H2' | 2.37 | 3.78 |
| 20 | A | H1' | 19 | C | H2' | 3 | 6 |
| 20 | A | H1' | 20 | A | H4' | 2.66 | 4.25 |
| 20 | A | H2 | 20 | A | H1' | 3 | 6 |
| 20 | A | H8 | 19 | C | H1' | 3 | 6 |
| 20 | A | H8 | 19 | C | H2' | 2 | 5 |
| 20 | A | H8 | 19 | C | H5 | 3 | 6 |
| 20 | A | H8 | 20 | A | H1' | 1.8 | 4.5 |
| 21 | G | H1' | 21 | G | H2' | 1.8 | 4.5 |
| 21 | G | H1' | 21 | G | H3' | 3 | 6 |
| 21 | G | H8 | 20 | G | H1' | 2.5 | 6 |

| | | | | | | | |
|---|---|---|---|---|---|---|---|
| 21 | G | H8 | 21 | G | H1' | 1.8 | 4.5 |
| 21 | G | H8 | 21 | G | H3' | 2.7 | 5 |
| 21 | G | H8 | 22 | C | H5 | 3 | 6 |
| 22 | C | H1' | 22 | C | H2' | 1.95 | 3.11 |
| 22 | C | H5 | 21 | G | H2' | 3 | 6 |
| 22 | C | H6 | 22 | C | H2' | 2.42 | 3.86 |
| 22 | C | H6 | 23 | U | H5 | 3 | 6 |
| 23 | U | H1' | 23 | U | H3' | 2.82 | 4.5 |
| 23 | U | H5 | 22 | C | H2' | 2.65 | 4.23 |
| 23 | U | H5 | 22 | C | H3' | 3 | 6 |
| 23 | U | H5 | 23 | U | H3' | 3 | 6 |
| 23 | U | H6 | 22 | C | H2' | 1.88 | 3 |
| 23 | U | H6 | 22 | C | H3' | 2.33 | 3.72 |
| 23 | U | H6 | 22 | C | H5 | 3 | 6 |
| 23 | U | H6 | 23 | U | H1' | 2.97 | 4.74 |
| 23 | U | H6 | 23 | U | H3' | 2.3 | 3.67 |
| 24 | G | H1' | 24 | G | H2' | 2.05 | 3.27 |
| 24 | G | H1' | 24 | G | H3' | 2.94 | 4.69 |
| 24 | G | H1' | 24 | G | H4' | 2.77 | 4.41 |
| 24 | G | H8 | 23 | U | H3' | 2.47 | 3.94 |
| 24 | G | H8 | 23 | U | H5 | 3 | 6 |
| 24 | G | H8 | 24 | G | H1' | 1.8 | 4.5 |
| 24 | G | H8 | 24 | G | H3' | 2.46 | 3.93 |
| 25 | U | H1' | 25 | U | H2' | 1.9 | 3.02 |
| 25 | U | H1' | 25 | U | H4' | 2.51 | 4.01 |
| 25 | U | H5 | 24 | G | H2' | 2.73 | 4.35 |
| 25 | U | H5 | 24 | G | H3' | 2.96 | 4.73 |
| 25 | U | H5 | 25 | U | H2' | 3 | 6 |
| 25 | U | H5 | 25 | U | H3' | 3 | 6 |
| 25 | U | H6 | 24 | G | H2' | 1.75 | 2.8 |
| 25 | U | H6 | 24 | G | H3' | 2.29 | 3.65 |
| 25 | U | H6 | 25 | U | H1' | 1.8 | 4.5 |
| 25 | U | H6 | 25 | U | H2' | 2.64 | 4.21 |
| 25 | U | H6 | 25 | U | H3' | 1.8 | 4.5 |
| 25 | U | H6 | 26 | C | H5 | 3 | 6 |
| 26 | C | H1' | 25 | U | H2' | 3 | 6 |
| 26 | C | H1' | 26 | C | H2' | 1.83 | 2.92 |
| 26 | C | H1' | 26 | C | H3' | 3 | 6 |
| 26 | C | H5 | 25 | U | H2' | 2.42 | 3.85 |
| 26 | C | H5 | 25 | U | H3' | 2.54 | 4.04 |
| 26 | C | H5 | 26 | C | H2' | 3.19 | 5.09 |
| 26 | C | H6 | 25 | U | H2' | 1.63 | 2.6 |
| 26 | C | H6 | 26 | C | H1' | 2.86 | 4.56 |
| 26 | C | H6 | 26 | C | H2' | 2.29 | 3.65 |
| 26 | C | H6 | 26 | C | H3' | 1.65 | 2.63 |
| 27 | PFA | C15 | 20 | A | H1' | 3 | 7 |
| 27 | PFA | C15 | 20 | A | H2 | 3 | 7 |
| 27 | PFA | C15 | 21 | G | H1' | 3 | 7 |
| 27 | PFA | C7 | 19 | C | H2' | 3 | 7 |
| 27 | PFA | C7 | 19 | C | H5 | 3 | 7 |
| 27 | PFA | C7 | 20 | A | H1' | 3 | 7 |

| | | | | | | | |
|---|---|---|---|---|---|---|---|
| 27 | PFA | C7 | 20 | A | H2 | 3 | 7 |
| 27 | PFA | C7 | 20 | A | H8 | 3 | 7 |
| 27 | PFA | C7 | 21 | G | H1' | 3 | 7 |
| 27 | PFA | C7 | 27 | PFA | C15 | 3 | 6 |
| 27 | PFA | H2 | 8 | G | H1' | 3 | 7 |
| 27 | PFA | H4 | 20 | A | H2 | 3 | 7 |
| 27 | PFA | H5 | 20 | A | H2 | 3 | 7 |
| 27 | PFA | H6 | 7 | A | H1' | 3 | 7 |
| 27 | PFA | H6 | 8 | G | H1' | 3 | 7 |
| 27 | PFA | H7 | 6 | C | H2' | 3 | 7 |
| 27 | PFA | H7 | 6 | C | H2' | 3 | 7 |
| 27 | PFA | H7 | 7 | A | H1' | 3 | 7 |
| 27 | PFA | H7 | 7 | A | H2 | 3 | 7 |
| 27 | PFA | H7 | 8 | G | H1' | 3 | 7 |
| 27 | PFA | H7 | 8 | G | H8 | 3 | 7 |
| 27 | PFA | H8 | 6 | C | H2' | 3 | 7 |
| 27 | PFA | H8 | 7 | A | H1' | 3 | 7 |

**Table S10**: Summary of select helical and base pair parameters for apo- and ligand-bound RNA constructs.

| PDB ID | Helical rise average (Å) | Helical twist average (°) | C1′-C1′ distances range (Å) | Major groove widths, direct P-P distance, range (Å) | Minor groove widths, direct P-P distance, range (Å) |
|---|---|---|---|---|---|
| Fully base paired RNAs | | | | | |
| 1QCU(1) | 2.7 ± 0.2 | 33.2 ± 1.4 | 10.6-10.7 | 15.2-16.7 | 16.8-17.8 |
| 1RXB(2) | 2.8 ± 0.2 | 32.6 ± 0.9 | 10.5-10.7 | 16.6-17.3 | 17.0-17.3 |
| 3ND4(3) | 2.6 ± 0.4 | 33.0 ± 3.4 | 10.3-10.7 | 15.2-18.4 | 17.0-17.7 |
| 4MS9(4) | 2.9 ± 0.2 | 32.9 ± 1.6 | 10.5-10.7 | 15.0-15.7 | 16.5-16.6 |
| 4NFO(5) | 2.7 ± 0.3 | 33.1 ± 2.8 | 10.5-10.6 | 15.9-17.6 | 17.1-17.7 |
| 4U37(6) | 2.9 ± 0.1 | 33.1 ± 1.2 | 10.4-10.7 | 14.5-14.8 | 16.7-16.9 |
| RNAs containing unbound r(CAG) motifs | | | | | |
| 3NJ7(7) | 2.8 ± 0.4 | 28.6 ± 7.0 | 10.6-11.7 | 20.2-25.9 | 16.4-17.2 |
| 4J50(8) | 2.2 ± 1.3 | 27.7 ± 68.2 | 10.4-11.4 | 17.3-19.8 | 16.5-17.3 |
| 5VH7(9) | 2.7 ± 0.5 | 30.6 ± 3.8 | 10.6-12.6 | 17.2-25.4 | 15.7-18.5 |
| 7VFT(10) | 2.7 ± 0.2 | 33.2 ± 1.4 | 10.4-11.6 | 18.6-23.8 | 16.4-17.5 |
| apo-r(1×CAG) | 2.7 ± 0.3 | 31.5 ± 2.7 | 10.4-12.3 | 15.9-21.1 | 16.4-17.6 |
| RNAs containing ligand-bound r(CAG) motifs | | | | | |
| r(CAG)-**1** | 2.7 ± 0.9 | 30.6 ± 10.7 | 10.4-13.6 | 16.8-25.1 | 15.7-19.1 |
| r(CAG)-**2** | 2.9 ± 0.7 | 32.4 ± 2.2 | 10.3-12.5 | 15.5-25.1 | 14.7-18.8 |
| r(CAG)-**3** | 2.3 ± 1.5 | 22.2 ± 36.6 | 10.4-14.2 | 15.9-26.5 | 14.8-21.4 |
| RNAs containing unbound and ligand-bound tau RNA A-bulge motifs | | | | | |
| 6VA1(11) | 2.3 ± 1.0 | 33.7 ± 8.4 | 9.6-11.0 | 16.7-25.3 | 14.4-21.4 |
| 6VA3(11) | 2.9 ± 1.2 | 33.8 ± 6.3 | 10.4-10.8 | 17.0-21.8 | 17.3-20.4 |
| RNAs containing unbound and ligand-bound r(CUG) motifs | | | | | |
| 9CPD(12) | 2.5 ± 0.6 | 32.8 ± 11.1 | 10.5-14.8 | 15.5-21.8 | 14.4-20.1 |
| 9CPG(12) | 2.2 ± 1.0 | 31.8 ± 3.0 | 10.5-14.0 | 17.2-25.5 | 15.6-20.6 |
| 9CPI(12) | 2.8 ± 0.5 | 30.6 ± 19.9 | 7.9-11.5 | 14.7-24.6 | 14.4-23.9 |
| 9CPJ(12) | 2.8 ± 0.3 | 30.9 ± 2.0 | 9.1-10.9 | 16.7-22.2 | 16.1-18.7 |
